# *Cinnamomum cassia* Extract and Its Novel Isolated Compound Suppress Inflammation via Autophagy Induction in Sepsis

**DOI:** 10.1101/2025.07.21.665831

**Authors:** Geon Park, Tam Thi Le, Tae Kyeom Kang, Yuna Jung, Wook-Bin Lee, Sang Hoon Jung

## Abstract

Uncontrolled inflammation is central to the development of diseases such as sepsis, and autophagy has emerged as a critical regulatory mechanism in this process. The ethanol extract of *Cinnamomum cassia* (EECC) was identified as a potent autophagy inducer through high-throughput LC3 reporter screening. EECC enhanced autophagic flux, as confirmed by RFP-GFP-LC3 imaging and immunoblotting. It also suppressed Toll-like receptor signaling and reduced pro-inflammatory cytokine production in macrophages. EECC inhibited nuclear factor-κB signaling in an autophagy-dependent manner, as this effect was reversed by autophagy inhibitors. To identify active constituents, 24 compounds were isolated from EECC, including six novel structures. Among the novel compounds, Cassitamine F exhibited dual activity as an autophagy inducer and inflammation suppressor. In a lipopolysaccharide-induced sepsis model, Cassitamine F significantly reduced serum levels of pro-inflammatory cytokines. These findings suggest that EECC and Cassitamine F may hold therapeutic potential for autophagy-targeted treatment of sepsis and other inflammation-related disorders.

**Highlights:**

- Ethanol extract of *Cinnamomum cassia* (EECC) identified as a potent autophagy inducer via high-throughput LC3 HiBiT screening.
- EECC suppresses Toll-like receptor signaling and pro-inflammatory cytokines through autophagy-dependent pathways.
- Twenty-four compounds were isolated from EECC, including six novel structures (Cassitamine A–F).
- Cassitamine F exhibits dual activity: induction of autophagy and inhibition of inflammation.
- Cassitamine F significantly reduces serum cytokine levels in an LPS-induced sepsis model, suggesting therapeutic potential.

## 1. Introduction

Inflammation is a fundamental biological response that protects the body from infections, injuries, and harmful stimuli. However, excessive or uncontrolled inflammation is a major contributor to various diseases, including autoimmune disorders, metabolic syndromes, cardiovascular diseases, and neurodegenerative conditions (Duan, Rao, & Sigdel, 2019; Xiang, Zhang, Jiang, Su, & Shi, 2023). One of the most severe manifestations of uncontrolled inflammation is sepsis, a life-threatening condition characterized by a dysregulated immune response to infection, leading to systemic inflammation, organ dysfunction, and high mortality rates (Angus & Van der Poll, 2013; Cohen, 2002). Despite advances in critical care, effective therapeutic strategies for sepsis remain limited, highlighting the urgent need for novel interventions that can modulate excessive inflammatory responses.

Autophagy, an evolutionarily conserved intracellular degradation process, plays a pivotal role in regulating inflammation and immune responses (Matsuzawa-Ishimoto, Hwang, & Cadwell, 2018; Mizushima & Levine, 2020). By degrading damaged organelles, misfolded proteins, and intracellular pathogens, autophagy helps maintain cellular homeostasis and mitigate inflammatory responses (Dikic & Elazar, 2018). Studies have shown that autophagy negatively regulates inflammatory pathways by inhibiting inflammasome activation, suppressing cytokine secretion, and modulating Toll-like receptor (TLR) signaling (Deretic, 2021; Saitoh & Akira, 2010; Shibutani, Saitoh, Nowag, Münz, & Yoshimori, 2015). Furthermore, autophagy has been implicated in sepsis regulation, as it protects against excessive inflammation and tissue damage by controlling mitochondrial dysfunction and oxidative stress (Ho et al., 2016). Given its critical role in inflammation and immune homeostasis, enhancing autophagy has emerged as a potential therapeutic strategy for inflammatory diseases, including sepsis.

*Cinnamomum cassia* (Chinese cinnamon), a widely used medicinal plant, has been traditionally employed in East Asian medicine for its anti-inflammatory, antioxidant, and immune-modulating properties (Hong, Yang, Kim, Eom, Lew, & Kang, 2012)(Ranasinghe et al., 2013)(X. Li et al., 2021; Zhang et al., 2019). Several studies have suggested that *C. cassia* and its bioactive components, such as cinnamaldehyde and flavonoids, exert immunomodulatory effects by modulating key signaling pathways involved in inflammation and oxidative stress (C.-y. Li et al., 2024). However, the precise mechanism through which *C. cassia* exerts its immunomodulatory effects, particularly its potential role in autophagy induction and sepsis regulation, remains largely unexplored.

Based on these considerations, we hypothesized that *C. cassia* induces autophagy and thereby suppresses inflammatory responses and lipopolysaccharide (LPS)-induced sepsis. To test this hypothesis, we conducted a comprehensive investigation using high-throughput LC3 HiBiT reporter assays to screen for autophagy inducers and HEK-TLR4 reporter assays to evaluate anti-inflammatory activity. We isolated 24 compounds from EECC, including several novel structures, and subsequently performed in vivo experiments in an LPS-induced sepsis model to evaluate the protective effects of bioactive compounds isolated from EECC. This study provides new insights into the immunomodulatory mechanisms of *C. cassia* and its potential application as a therapeutic agent for sepsis and other inflammatory disorders.

## 2. Materials and Methods

### 2.1. Natural product extracts for screening

A total of 1,629 crude natural product extracts were obtained from the Natural Product Library of the Korea Institute of Science and Technology (KIST), Gangneung Institute, Gangneung, Republic of Korea. This library consists of extracts derived from Korean native plants and was used for high-throughput screening of autophagy-inducing activity.

### 2.2. Plant materials and extraction

Dried bark of *C. cassia* was purchased from DUSONAE HERB (Yeongcheon, Gyeongsangbuk-do, South Korea). A total of 9 kg of the dried material was extracted with 95% ethanol (EtOH) at room temperature for three days. The extract was filtered through Whatman No. 1 filter paper (Whatman, Buckinghamshire, UK), and the extraction procedure was repeated twice under the same conditions. The combined filtrates were concentrated under reduced pressure at 45°C using a rotary evaporator (Rotavapor R-100; Büchi, Flawil, Switzerland), yielding 516.3 g of extract. The EtOH extract of *C. cassia* (EECC) was stored at −20°C until further analysis.

### 2.3. Autophagy LC3 HiBiT reporter assay

HEK293 cells stably expressing HiBiT-tagged LC3B were purchased from Promega (Madison, WI, USA). The luminescence-based autophagy assay was performed using the Nano-Glo® HiBiT Lytic Detection System (Promega) following the manufacturer’s instructions. Cells were seeded in white 96-well plates (SPL Life Sciences, Pocheon, Republic of Korea) at a density of 2 × 10⁴ cells per well and incubated overnight. The next day, cells were treated with either 200 nM rapamycin (InvivoGen, Waltham, USA), 50 nM bafilomycin A1 (InvivoGen), or various concentrations of EECC (12.5, 25, and 50 μg/mL). After 24 hours of treatment, luminescence was developed using the detection reagent, with gentle shaking on a plate shaker (Thermo Scientific, Waltham, MA, USA) at 350 rpm for 2–10 minutes. Luminescence was measured using a GloMax® Navigator microplate luminometer (Promega).

### 2.4. TLR reporter assay using HEK-Blue hTLR cells

HEK-Blue™ hTLR reporter cells (TLR2/1, TLR3, TLR4, TLR5, TLR7, TLR8, and TLR9) were seeded in 96-well plates at a density of 5 × 10⁴ cells/well. Cells were pretreated with EECC (50 μg/mL) for 1 hour, followed by stimulation with the corresponding TLR agonists: Pam3CSK4 (10 ng/mL for TLR2/1), poly(I:C) (1 μg/mL for TLR3), LPS (1 μg/mL for TLR4), FLA-ST (10 ng/mL for TLR5), R848 (1 μg/mL for TLR7/8), and ODN2006 (10 μg/mL for TLR9). After 24 hours of incubation, TLR activation was assessed by measuring the activity of secreted embryonic alkaline phosphatase (SEAP) in cell supernatants using Quanti-Blue™ (InvivoGen) reagent, diluted 1:10 according to the manufacturer’s protocol. Absorbance was measured at 630 nm using a MULTISKAN Sky microplate reader (Thermo Fisher Scientific) after 20–30 minutes of incubation.

### 2.5. NF-κB and IRF Pathway Assays Using THP1 Reporter Cells

THP1-Blue™-ISG and THP1-XBlue™ cells were seeded in 96-well round-bottom plates at a density of 5 × 10⁵ cells/well. Cells were pretreated with EECC at concentrations of 12.5, 25, and 50 μg/mL for 1 hour, followed by stimulation with Pam3CSK4 (100 ng/mL) or LPS (1 μg/mL) for 24 hours. SEAP activity in the culture supernatants was then measured using the Quanti-Blue™ assay following the manufacturer’s instructions. Absorbance was read at 630 nm using the MULTISKAN Sky microplate reader. (Kang et al., 2023)

### 2.6. Cell Viability

HEK-Blue hTLRs (2/1, 3, 4, 5, 7, 8, and 9) cells were seeded at a density of 5 × 10^4^ cells/well in 96 well plate. Cells were pretreated with 50 μg/ml EECC for 24h to measure cell viability. MTT (3-4,5-dimethylthiazol-2-yl)-2,5-diphenyltetrazolium bromide) solution was added to the cells (final concentration : 0.5 mg/ml) for 1h at 37°C. Optical density was measured using a MULTISKAN Sky microplate reader with a 570 nm wavelength. Experiments were independently repeated three times. (Kang et al., 2023)

### 2.7. RFP-GFP-LC3B HeLa cell assay

HeLa-Difluo™ hLC3 cells (InvivoGen) were seeded at a density of 3 × 10⁵ cells/well onto confocal dishes (SPL Life Sciences) and incubated overnight. The following day, cells were treated with 1 μM rapamycin or EECC at concentrations of 10 and 50 μg/mL. After 24 hours, live-cell imaging was performed using a confocal laser scanning microscope (LSM 900; Zeiss, Oberkochen, Germany). Image acquisition and analysis were carried out using Zen 3.4 software (Zeiss).

### 2.8. Western blot analysis

RAW 264.7 cells were lysed in RIPA buffer (Thermo Scientific) supplemented with phosphatase and protease inhibitor cocktails (APE BIO, Boston, USA) on ice for 15 minutes. The lysates were then sonicated (Qsonica, Newtown, USA) and further incubated on ice for 30 minutes. After centrifugation at 13,000 rpm for 30 minutes at 4°C, the supernatants were collected as total protein extracts. Protein concentrations were determined using the Bradford assay. Expression levels of autophagy-related proteins, including LC3 (D11; Cell Signaling Technology, MA, USA), Beclin-1 (E-8; Santa Cruz Biotechnology, TX, USA), and p62 (EPR4844; Abcam, Cambridge, UK) were analyzed.

### 2.9. Enzyme-linked immunosorbent assay (ELISA)

Raw 264.7 cells were seeded in a 96 well plate (Thermo scientific) at a density of 1 × 10^5^ cells/well and bone-marrow derived macrophages (BMDMs) were seeded in a 96 well plate at a density of 2 × 10^5^ cells/well. After 24h incubation, the cells treated with 1 μg/ml LPS and EECC (12.5, 25, and 50 μg/ml). The cell supernatants were collected and transferred to a new Eppendorf tube and used for measuring. The levels of cytokines using the mouse TNF-α and mouse IL-6 were quantified using ELISA kits (ELISA MAX^TM^ TNF-α, IL-6; BioLegend, San Diego, USA) according to the manufacturer’s instructions.

### 2.10. Isolation and identification

The EECC was suspended in water and then partitioned with methylene chloride (MC), ethyl acetate (EA), and *n*-butanol (Bu) to obtain MC fraction (90.1 g), EA fraction (43.5 g), Bu fraction (240.7 g), and water layer. The EA-soluble fraction (CCEA) was subjected to a silica gel column using a stepwise gradient of *n*-hexane (Hx)-EA (15:0 to 100% EA) to yield 17 subfractions (CCEA 1 ̶ 17). Subfraction CCEA 8 was purified using preparative MPLC utilizing acetonitrile (ACN)/H_2_O (gradient elution, increasing from 10% ACN to 100% ACN over 50 min, flow rate of 8 mL/min, YMC C18 column) to yield coumarin (**1**, 4.6 mg), cinnamaldehyde (**4**, 6.5 mg), and (*E*)-*o*-methoxycinnamaldehyde (**5**, 3.3 mg). Subfraction CCEA 12 was separated on an open column packed with octadecylsilane (ODS) using methanol (MeOH)/H_2_O as the mobile phase, yielding 94 subfractions (CCEA 12-1−94). Subfractions CCEA 12-30, 39, 51, 73, and 94 were selected for further purification using preparative MPLC utilizing ACN/H_2_O (gradient elution, increasing from 20% ACN to 100% ACN over 60 min, flow rate of 8 mL/min, YMC C18 column) to obtain (*E*)-cinnamyl alcohol (**2**, 4.5 mg), cinnamic acid (**3**, 7.6 mg), (*E*)-3-(2-hydroxyphenyl)-propenal (**6**, 2.3 mg), 2-hydroxycinnamyl alcohol (**7**, 5.5 mg), cinncassin N (**8**, 3.3 mg), cinncassin O (**9**, 2.7 mg). Subfraction CCEA 12-90 was applied to a Sephadex LH-20 column eluted with MeOH/H_2_O (1:2 → 100% MeOH) to produce 55 subfractions (12-90-1 to 12-90-55). Subfractions 12-90-2, 28, 38, 45, and 55 underwent further purification by preparative MPLC, employing a gradient elution of ACN and water (from 20% to 100% ACN over 60 minutes) at a flow rate of 8 mL/min on a YMC C18 column, which led to the isolation of litseachromolaevane A (**12**, 1.2 mg), cinncassin L (**13**, 1.6 mg), cassitamine B (**20**, 2.1 mg), cassitamine C (**21**, 1.9 mg), and cassitamine D (**22**, 1.7 mg). Similarly, subfraction CCEA 5 was loaded on an open column packed with ODS using MeOH/H_2_O as the mobile phase, collecting 51 subfractions (CCEA 5-1−51). Subfraction CCEA 5-50 then was loaded onto a Sephadex LH-20 column eluted with MeOH/H_2_O (1:2 → 100% MeOH) to obtain 50 subfractions (5-50-1 to 5-50-50). From subfraction CCEA 5-50-18, cassitamine A (**19**, 0.5 mg), cassitamine E (**23**, 1.2 mg), and cassitamine F (**24**, 1.1 mg) were purified via preparative MPLC, employing a gradient elution of ACN and water (from 20% to 100% ACN over 40 minutes) at a flow rate of 8 mL/min on a YMC C18 column. Similarly, curcudiol-10-one (**14**, 1.3 mg), 13-oxo-(9*E*,11*E*)-octadecadienoic acid (**15**, 2.6 mg), costunolide (**16**, 5.3 mg), rhodomicranin F (**17**, 1.0 mg), and cinnaphenylpropanoid M (**18**, 1.1 mg) were isolated from subfraction CCEA 5-50-8 using preparative MPLC (gradient elution ACN/H_2_O, increasing from 50% ACN to 100% ACN over 60 min, flow rate of 8 mL/min, YMC C18 column). Finally, *β*-sitosterol (**10**, 15.3 mg) and *β*-sitosterol *β*-D-glucoside (**11**, 11.7 mg) were obtained through the crystallization of subfractions CCEA 10 and 16, respectively.

*Cassitamine A (**19**)*: yellowish gum; UV *λ*_max_ (log *ε*) (methanol): 280 (2.20), 220 (2.10), 200 (2.11) nm; ^1^H NMR (500 MHz) and ^13^C NMR (125 MHz) data in chloroform-*d*, see Table 1; HRESIMS (positive-ion mode) *m/z* 289.0837 [M + Na]^+^ (calcd. for C_17_H_14_O_3_Na, 289.0 835).

*Cassitamine B (**20**)*: brown gum; UV *λ*_max_ (log *ε*) (methanol): 340 (2.54), 284 (2.90), 222 (2.00) nm; ^1^H NMR (500 MHz) and ^13^C NMR (125 MHz) data in methanol-*d*_4_, see Table 1; HRESI MS (positive-ion mode) *m/z* 301.0839 [M + Na]^+^ (calcd. for C_18_H_14_O_3_Na, 301.0835).

*Cassitamine C (**21**)*: brown gum; UV *λ*_max_ (log *ε*) (methanol): 350 (2.40), 288 (2.52), 220 (2.53) nm; ^1^H NMR (500 MHz) and ^13^C NMR (125 MHz) data in methanol-*d*_4_, see Table 1; HRESI MS (positive-ion mode) *m/z* 317.0791 [M + Na]^+^ (calcd. for C_18_H_14_O_4_Na, 317.0784).

*Cassitamine D (**22**)*: yellowish gum; UV *λ*_max_ (log *ε*) (methanol): 280 (2.08), 215 (2.84) nm; ^1^H NMR (500 MHz) and ^13^C NMR (125 MHz) data in chloroform-*d*, see Table 2; HRESIMS (po sitive-ion mode) *m/z* 457.1623 [M + Na]^+^ (calcd. for C_26_H_26_O_6_Na, 457.1622).

*Cassitamine E (**23**)*: yellowish gum; UV *λ*_max_ (log *ε*) (methanol): 280 (2.08), 215 (2.84) nm; ^1^ H NMR (500 MHz) and ^13^C NMR (125 MHz) data in chloroform-*d*, see Table 1; HRESIMS (positive-ion mode) *m/z* 305.1154 [M + Na]^+^ (calcd. for C_18_H_18_O_3_Na, 305.1151).

*Cassitamine F (**24**)*: yellowish gum; UV *λ*_max_ (log *ε*) (methanol): 330 (3.11), 235 (2.48), 198 (2.85) nm; ^1^H NMR (500 MHz) and ^13^C NMR (125 MHz) data in chloroform-*d*, see Table 1; H RESIMS (positive-ion mode) *m/z* 287.1046 [M + Na]^+^ (calcd. for C_18_H_16_O_2_Na, 287.10 43).

### 2.11. Chemicals and apparatus

The NMR spectra (^1^H, ^13^C, DEPT135, ^1^H–^1^H COSY, HSQC, HMBC, and NOESY) were obtained using a Varian 500 MHz NMR spectrometer. The HRESIMS data were acquired on a TripleTOF 6600 (SCIEX, U.S.A.) mass spectrometer. HPLC chromatograms and ESIMS data were obtained using an Agilent 1200 system connected to a 6120 quadrupole MSD with a Hydrosphere YMC C18 column (5 μm, 250 × 4.6 mm). Preparative MPLC was carried out using a YMC MPLC system with a YMC ODS column (5 μm, 250 × 20 mm). HPLC-grade acetonitrile and distilled water were purchased from Fisher Scientific (Pittsburgh, PA, USA). Silica gel (Merck, Darmstadt, Germany) and Sephadex LH-20 (Pharmacia, Uppsala, Sweden) were used for open column chromatography.

### 2.12. LPS-Induced Sepsis Model

Animal experiments were conducted in a pathogen-free facility at the Korea Institute of Science and Technology (KIST), Gangneung Institute, in accordance with institutional guidelines approved by the KIST Animal Care and Use Committee (Approval No. KIST-IACUC-2023-061). Six-week-old female C57BL/6 mice (16–18 g) were purchased from Koatech (Gyeonggi-do, Republic of Korea) for use in the study. Sepsis was induced by intraperitoneal (i.p.) injection of LPS at a dose of 10 mg/kg body weight dissolved in sterile PBS. For treatment, compounds 21 (Cassitamine C), 23 (Cassitamine E), and 24 (Cassitamine F) were administered intraperitoneally at 10 mg/kg body weight simultaneously with LPS. The control group received an equivalent volume of PBS along with LPS. After 16 hours, mice were sacrificed, and blood samples were collected. Serum was separated and stored at –80°C until analysis. Inflammatory cytokine levels in serum were measured using the LEGENDplex™ Mouse Inflammation Panel (13-plex, BioLegend, San Diego, USA) on a V-bottom plate, according to the manufacturer’s instructions. Data were analyzed using LEGENDplex™ Data Analysis Software (BioLegend) (Lim et al., 2025)

#### Statistical analysis

All statistical evaluations were conducted using GraphPad Prism software (version 9.0; GraphPad, San Diego, CA, USA). Comparisons between two groups were made using an unpaired, two-tailed Student’s t-test, assuming a 95% confidence level. Information regarding sample sizes, replicates, and exact P values is included in the respective figure legends.

## 3. Results

### 3.1. Identification of *C. cassia* as an Autophagy-Inducing Extract

To identify natural product extracts that induce autophagy, a high-throughput screening was performed using the LC3 HiBiT reporter assay in a library of 1,679 natural product extracts (**Figure 1A**). The screening revealed that EtOH extract of *C. cassia* (EECC) exhibited significant autophagy-inducing activity, comparable to rapamycin, a well-known autophagy inducer, while Bafilomycin A, an autophagy inhibitor, suppressed LC3 reporter activity. To further confirm the autophagy-inducing potential of EECC, HEK293 Autophagy LC3 HiBiT Reporter cells were treated with increasing concentrations (12.5, 25, and 50 µg/mL) of *C. cassia* extract, resulting in a dose-dependent increase in LC3 reporter activity (**Figure 1B**). Additionally, fluorescence microscopy using HeLa-Difluo™ hLC3 reporter cells, which express the RFP-GFP-LC3 fusion protein, demonstrated autophagy activation upon EECC treatment, as indicated by the formation of RFP- and GFP-positive puncta representing autophagosomes. Furthermore, a reduction in GFP fluorescence with sustained RFP signal suggests the fusion of autophagosomes with lysosomes(Kimura, Noda, & Yoshimori, 2007; Loos, Toit, & Hofmeyr, 2014), confirming the progression of autophagic flux (**Figure 1C**). Immunoblot analysis of autophagy markers LC3-II, Beclin-1, and p62 in HEK293 cells also supported the autophagy-inducing effect of EECC, as shown by increased LC3-II and Beclin-1 levels (**Figure 1D**). Thus, these findings collectively demonstrate that *C. cassia* extract promotes autophagy induction and progression, supporting its potential as a modulator of the autophagy pathway.

**Figure 1.**
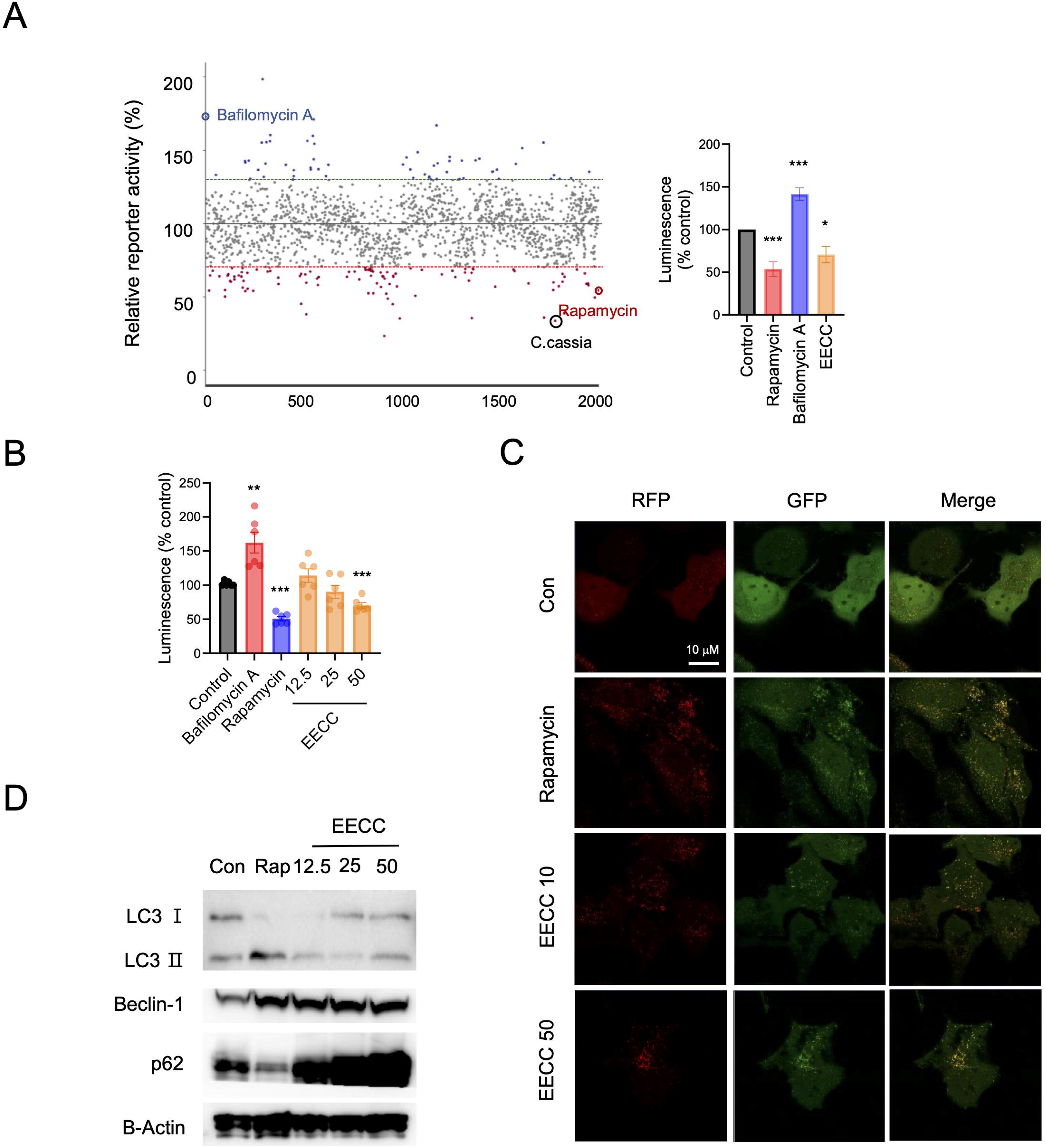
*C. cassia* extract induces autophagy in multiple cellular models. (A) Screening of 1679 natural product extracts for autophagy induction using the LC3 HiBiT reporter assay. The relative reporter activity (%) is plotted for each extract, with the control mean set at 100%. Dashed lines indicate thresholds for +30% and -70% of the control mean. Bafilomycin A (blue), rapamycin (red), the *C. cassia* extract (black) are highlighted. Each dot represents an individual natural product extract screened in the assay. The accompanying bar graph shows relative reporter activity (%) for selected conditions. (B) Dose-dependent effects of EECC on autophagy induction in HEK293 Autophagy LC3 HiBiT Reporter cells, assessed by luminescence measurements. (C) Confocal fluorescence imaging of HeLa-Difluo™ hLC3 cells expressing RFP::GFP::LC3, showing autophagosome formation following treatment with EECC. (D) Immunoblot analysis of autophagy markers LC3-II, Beclin 1, and p62 in HEK293 cells treated with EECE at increasing concentrations. Data are presented as mean ± SEM. Significant differences at ***p < 0.001, **p < 0.01, *p < 0.05 versus the Control group.

### 3.2. *C. cassia* Extract Suppresses Toll-Like Receptor Activation

Autophagy induction primarily exerts anti-inflammatory effects by modulating immune signaling pathways (Deretic & Levine, 2018; Deretic, Saitoh, & Akira, 2013). To assess the anti-inflammatory potential of EECC, we examined its impact on TLR signaling using HEK-Blue hTLR reporter cells, which express different TLRs (hTLR2, hTLR3, hTLR4, hTLR5, hTLR7, hTLR8, hTLR9). Prior to TLR stimulation, cell viability was confirmed by MTT assay, showing no cytotoxic effect of EECC at 50 μg/mL (**Figure 2A**). Cells were stimulated with specific TLR agonists (Pam3CSK4 (TLR2/1), poly(I:C) (TLR3), LPS (TLR4), FLA-ST (TLR5), R848 (TLR7, TLR8), ODN2006 (TLR9)) and treated with *C. cassia* extract. TLR activation was measured by secreted embryonic alkaline phosphatase (SEAP) activity at OD 630 nm. The results showed that EECC significantly inhibited TLR activation across multiple receptors (**Figure 2B-H**).

**Figure 2.**
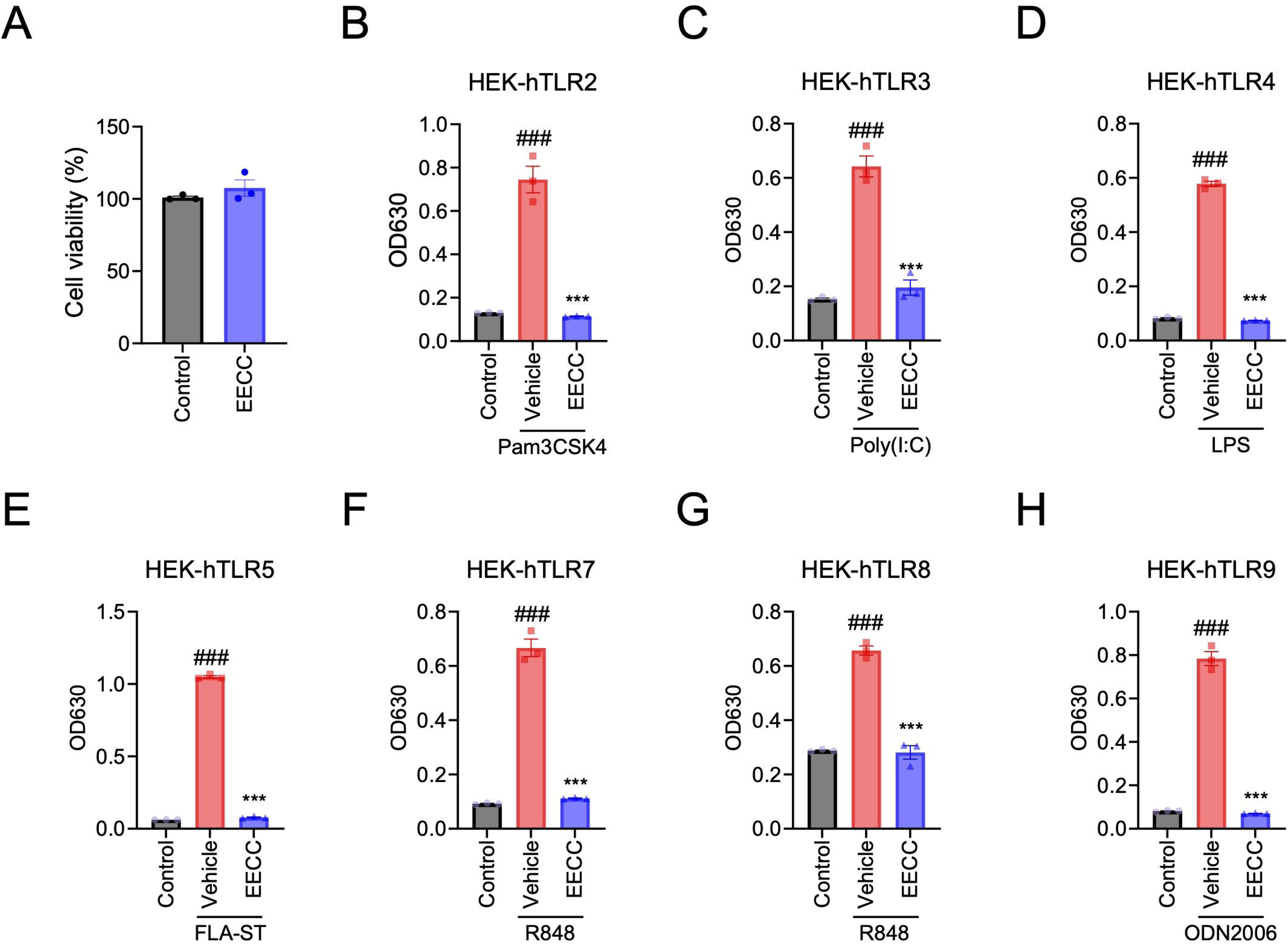
*C. cassia* extract suppresses Toll-like receptor (TLR) activation in HEK-Blue hTLR cells. (A) Cell viability of HEK-Blue hTLR2 cells treated with 50 μg/mL EECC, measured using the MTT assay at OD 570 nm. (B–H) Inhibitory effects of EECC on TLR signaling in HEK-Blue reporter cells expressing human TLR2 (B), TLR3 (C), TLR4 (D), TLR5 (E), TLR7 (F), TLR8 (G), and TLR9 (H). Cells were stimulated with the corresponding TLR agonists: Pam3CSK4, poly(I:C), LPS, FLA-ST, R848, or ODN2006, followed by EECC treatment. TLR activation was quantified by measuring secreted embryonic alkaline phosphatase (SEAP) activity at OD 630 nm. Data are presented as mean ± SEM. Significant differences at ###p < 0.001 versus the Control group, ***p < 0.001 versus the Vehicle group.

### 3.3. *C. cassia* Extract Reduces Pro-Inflammatory Cytokine Production in Macrophages

The ability of EECC to suppress pro-inflammatory cytokine secretion was assessed in RAW 264.7 macrophages and bone marrow-derived macrophages (BMDMs) stimulated with LPS. ELISA analysis revealed that EECC treatment significantly reduced IL-6 and TNF-α levels in a dose-dependent manner (**Figure 3A-D**), supporting its potential as an anti-inflammatory agent.

**Figure 3.**
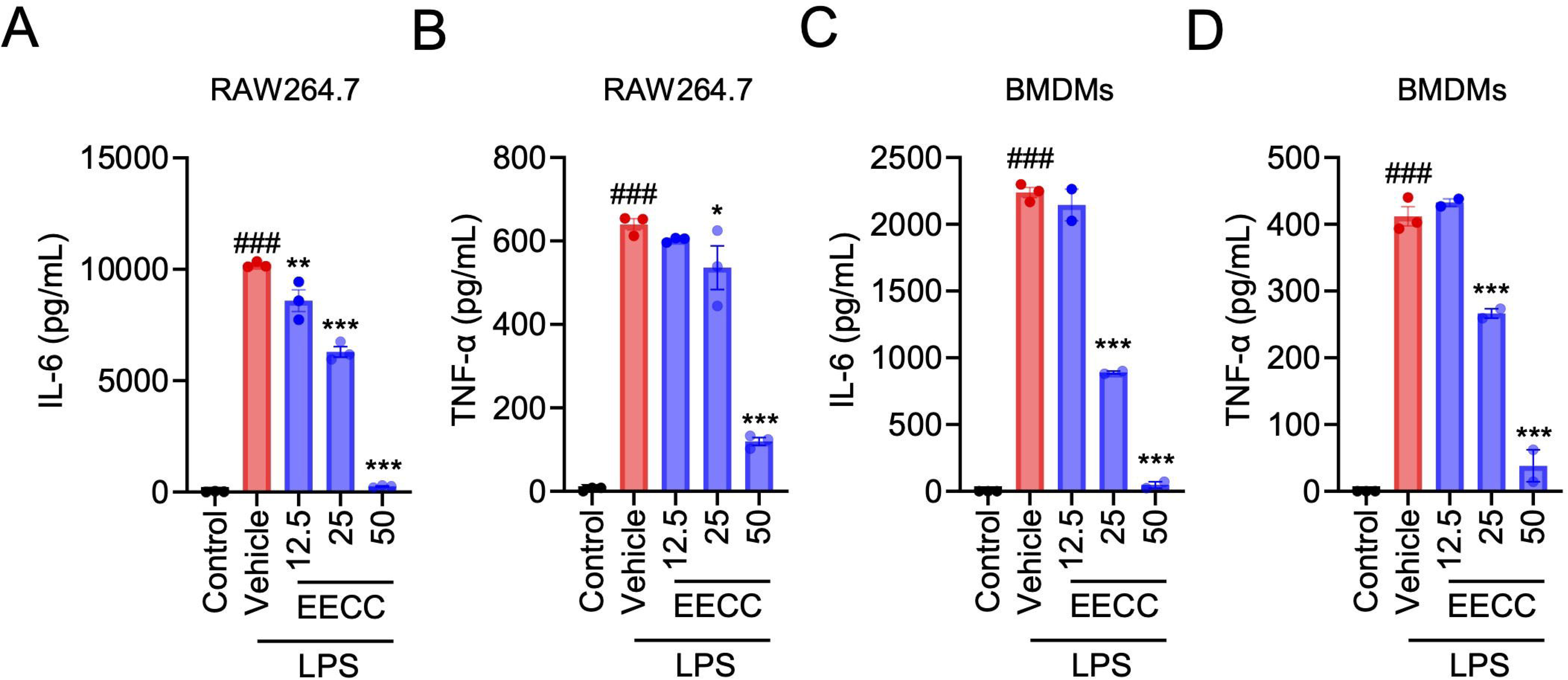
*C. cassia* extract suppresses pro-inflammatory cytokine production in macrophages. RAW 264.7 macrophages (A-B) and bone marrow-derived macrophages (BMDMs) (C-D) were stimulated with LPS and treated with EECE (12.5, 25, and 50 µg/mL). IL-6 (A, C) and TNF-α (B, D) secretion was measured by ELISA. Data are presented as mean ± SEM. Significant differences at ###p < 0.001 versus the Control group, ***p < 0.001, **p < 0.01, *p < 0.05 versus the Vehicle group.

### 3.4. *C. cassia* Extract Inhibits NFκB/AP1 and IRF Pathway Activation

To further elucidate the mechanism underlying the anti-inflammatory effects of EECC, we investigated its impact on NFκB/AP1 and IRF signaling using THP1-ISG and THP1-XBlue reporter cells. Cells were stimulated with LPS or Pam3CSK4 to activate these pathways and then treated with *C. cassia* extract. SEAP activity analysis revealed that EECC treatment led to a dose-dependent inhibition of both NFκB/AP1 and IRF activation (**Figure 4A-D**). Therefore, these findings suggest that EECC its anti-inflammatory effects by inhibiting TLR activation, suppressing pro-inflammatory cytokine production, and downregulating NFκB/AP1 and IRF signaling pathways.

**Figure 4.**
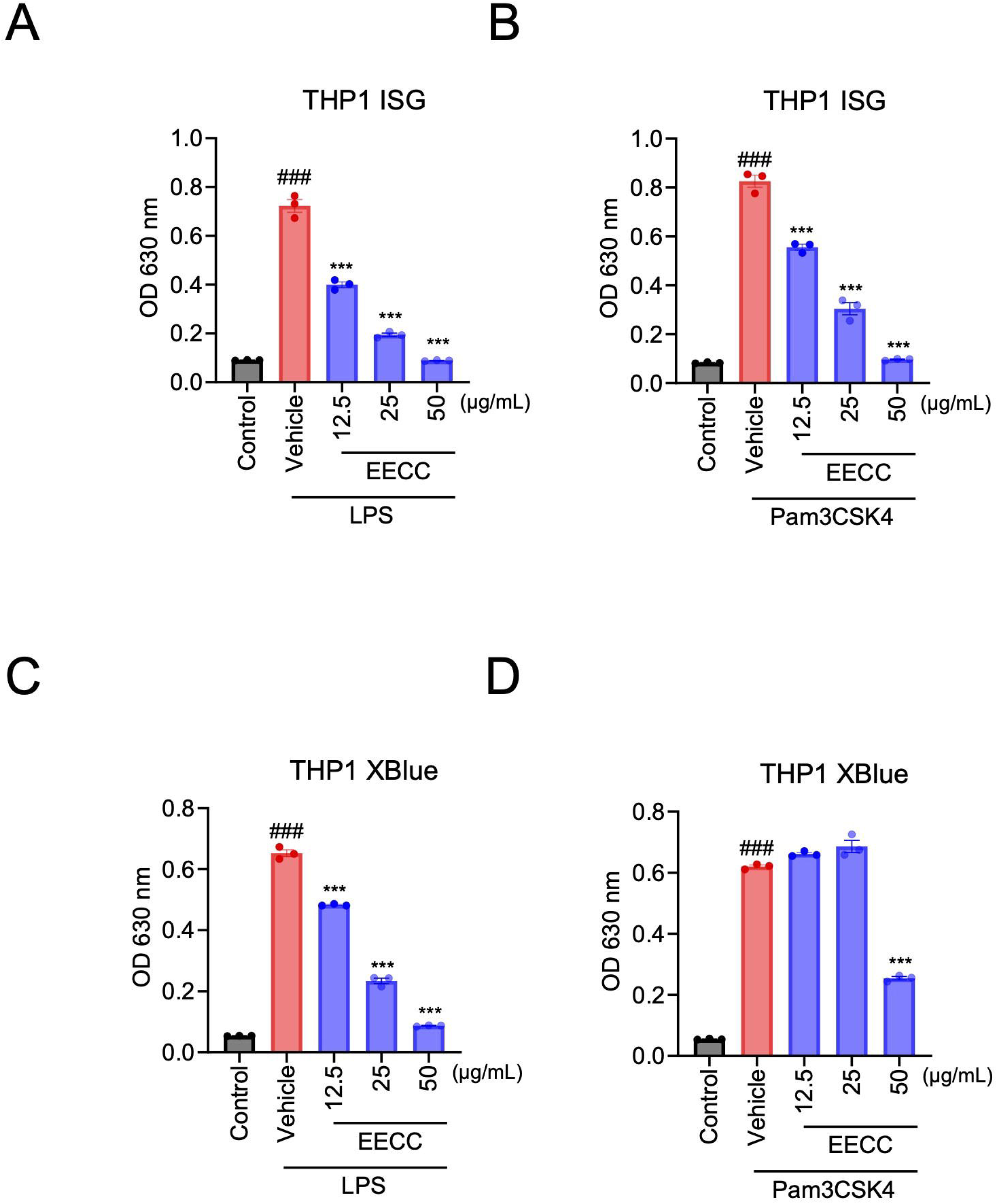
*C. cassia* extract inhibits NFκB and IRF pathway activation in THP1 reporter cells. THP1-ISG (A-B) and THP1-XBlue (C-D) reporter cells were used to evaluate the effects of EECE on NFκB/AP1 and IRF signaling. Cells were stimulated with LPS (A, C) or Pam3CSK4 (B, D) to activate the respective pathways, followed by treatment with *C. cassia* extract (12.5, 25, and 50 µg/mL). Activation of NF-κB/AP1 and IRF was assessed by monitoring the activity of secreted embryonic alkaline phosphatase (SEAP), measured at OD 630 nm. Data are presented as mean ± SEM. Significant differences at ###p < 0.001 versus the Control group, ***p < 0.001 versus the Vehicle group.

### 3.5. EECC Suppresses NF-κB Activation in an Autophagy-Dependent Manner

To determine whether this suppression was autophagy-dependent, cells were co-treated with autophagy inhibitors bafilomycin A or wortmannin. Co-treatment restored NF-κB activation, indicating that EECC-mediated inhibition of NF-κB occurs through an autophagy-dependent mechanism (**Figure 5**). These findings suggest that EECC regulates inflammatory signaling by inducing autophagy, which in turn suppresses NF-κB activation.

**Figure 5.**
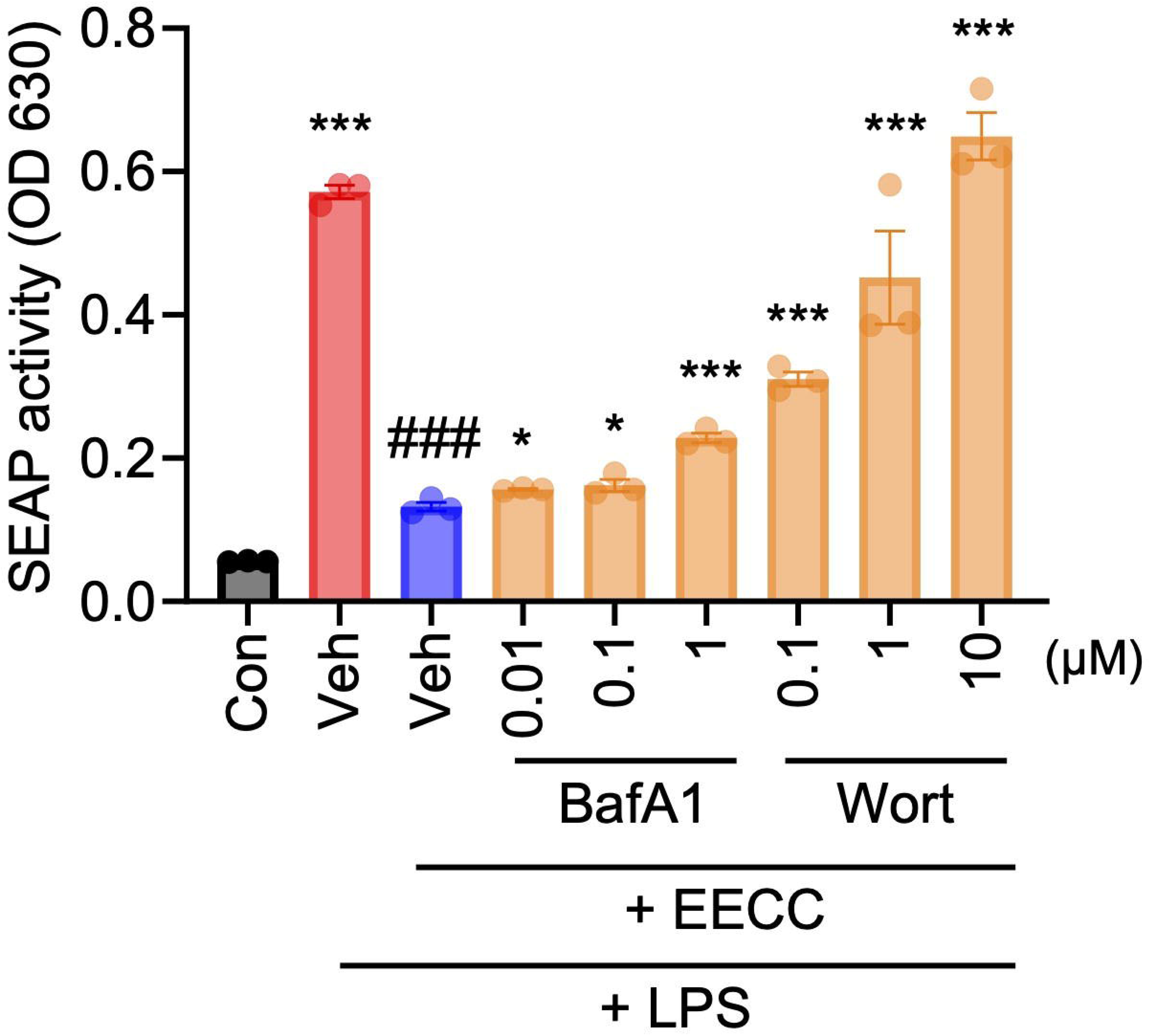
EECC inhibits NF-κB activation, but autophagy inhibition restores NF-κB activity. THP1-XBlue reporter cells were treated with EECC, bafilomycin, and wortmannin (indicated concentration) at increasing concentrations, and NF-κB activation was measured by SEAP activity at OD 630 nm. Data are presented as mean ± SEM. Significant differences at ###p < 0.001 versus the LPS group, ***p < 0.001, **p < 0.01, *p < 0.05 versus the EECC group.

### 3.6. Isolation and Structural Characterization of Compounds from *C. cassia*

To identify the bioactive components responsible for the autophagy-inducing and anti-inflammatory properties of *C. cassia*, we fractionated the EECC using high-performance liquid chromatography (HPLC). This process resulted in the isolation of 24 compounds, including six novel compounds (19–24), which were structurally characterized based on spectroscopic analysis (**Figure 6A**).

**Figure 6.**
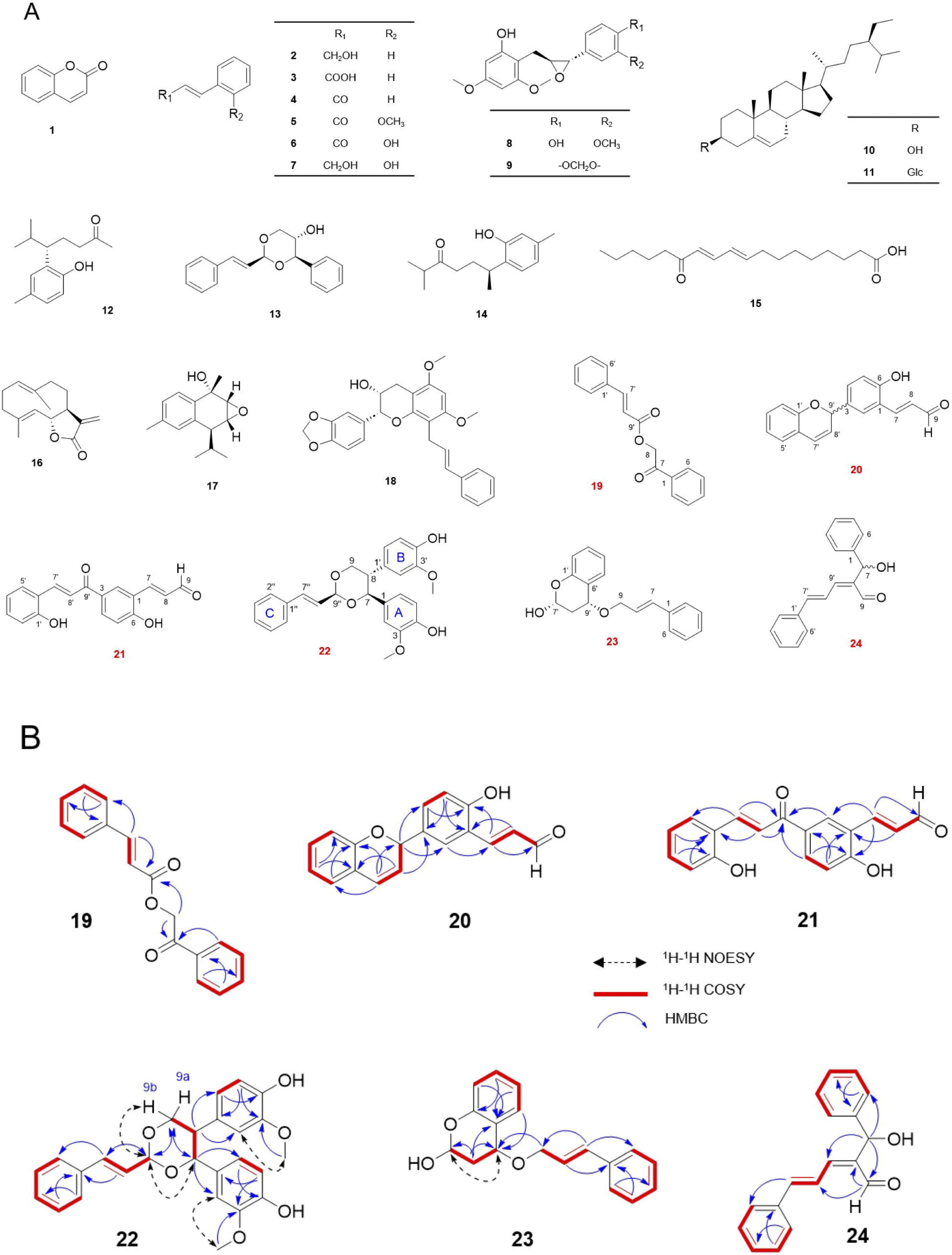
Chemical structures of compounds isolated from *C. cassia* extract and key NOESY, COSY, and HMBC correlations of representative compounds 19–24. (A) Structures of 24 compounds isolated from EECC are shown. Among them, six novel compounds (19–24, highlighted in red) were identified. (B) Structural elucidation of novel compounds 19–24 was performed using Nuclear Overhauser Effect Spectroscopy (NOESY) (dashed lines), Correlation Spectroscopy (COSY) (bold lines), and Heteronuclear Multiple Bond Correlation (HMBC) (arrows).

Repeating the process of column chromatography on EECC resulted in the isolation of a new natural product (**19**), five new compounds (**20-24**), along with other eighteen known compounds (**1**-**18**). By comparing the physical and spectroscopic data of the isolated compounds with the information recorded in existing literature, the known isolates were determined as coumarin (**1**),(Riga, Happyana, Quentmeier, Zammarelli, Kayser, & Hakim, 2021) (*E*)-cinnamyl alcohol (**2**),(Mcdonald, Turner, Gabbey, Balasubramanian, & Rousseaux, 2024) cinnamic acid (**3**),(Malik & Chakraborty, 2010) cinnamaldehyde (**4**),(Hayashi, Tanimoto, Zhang, Morimoto, Nishiyama, & Kakiuchi, 2012) (*E*)-*o*-methoxycinnamaldehyde (**5**),(Fernandes, Ranjan, & Choudhary, 2024) (*E*)-3-(2-hydroxyphenyl)propenal (**6**),(Matsuoka et al., 2023) 2-hydroxycinnamyl alcohol (**7**),(Chan, Wu, Wu, & Wu, 2002) cinncassin N (**8**),(X. Liu et al., 2018) cinncassin O (**9**),(X. Liu et al., 2018) *β*-sitosterol (**10**),(Ododo, Choudhury, & Dekebo, 2016) *β*-sitosterol *β*-D-glucoside (**11**),(Satmbekova, Srivedavyasasri, Orazbekov, Omarova, Datkhayev, & Ross, 2018) litseachromolaevane A (**12**),(Kiyotsuka, Katayama, Acharya, Hyodo, & Kobayashi, 2009) cinncassin L (**13**),(X. Liu et al., 2018) curcudiol-10-one (**14**),(Kuo & Yu, 1996) 13-oxo-(9*E*,11*E*)-octadecadienoic acid (**15**),(Trapp, Kai, Mithöfer, & Rodrigues, 2015) costunolide (**16**),(Fang, Sang, Chen, Gosslau, Ho, & Rosen, 2005) rhodomicranin F (**17**),(Yan et al., 2023) cinnaphenylpropanoid M (**18**).(Fan et al., 2024)

Cassitamine A (**19**) was yielded as a yellowish gum, and its molecular formula of C_17_H_14_O_3_ was established from the HRESIMS ion peak at m/z 289.0837 [M + Na]^+^ in positive ion mode (calcd. for C_17_H_14_O_3_Na, 289.0835). The ^1^H NMR spectrum exhibited characteristic resonances for two monosubstituted phenyl groups at [*δ*_H_ 7.99 (2H, d, *J* = 7.7 Hz, H-2,6), 7.54 (2H, t, *J* = 7.6 Hz, H-3,5), and 7.65 (1H, t, *J* = 7.4 Hz, H-4)] and [*δ*_H_ 7.59 (2H, overlapped, H-2’,6’) and 7.43 (3H, overlapped, H-2’,3’,5’)]; one disubstituted *trans*-olefin at *δ*_H_ 7.84 (1H, d, *J* = 16.0 Hz, H-7’) and 6.64 (1H, d, *J* = 16.0 Hz, H-8’); and one oxymethylene at *δ*_H_ 5.51 (2H, s, H-8). The ^13^C spectrum revealed characteristic resonances of a carboxylate group (*δ*_C_ 166.37, C-9’) and a carbonyl group (*δ*_C_ 192.26, C-7). The positions of those groups connecting two phenyl rings were determined through HMBC correlations from H-7’ (*δ*_H_ 7.84) to C-6’ (*δ*_C_ 128.28) and C-9’ (*δ*_C_ 166.37), from H-8 (*δ*_H_ 5.51) to C-7 (*δ*_C_ 192.26) and C-9’ (*δ*_C_ 166.37), and from H-6 (*δ*_H_ 7.99) to C-7 (*δ*_C_ 192.26) (**Figure 6B**). Consequently, **19** was identified as cinnamic acid 2-oxo-2-phenylethyl ester, which has been reported as a synthetic artifact.(Khamarui, Maiti, & Maiti, 2015) Its structure was shown in **Figure 6A**.

Cassitamine B (**20**) was isolated as a brown gum. Its molecular formula was determined as C_18_H_14_O_3_ by the HRESIMS ion peak at m/z 301.0839 [M + Na]^+^ in positive ion mode (calcd. for C_18_H_14_O_3_Na, 301.0835). The ^1^H NMR data displayed signals corresponding to a chromene group at *δ*_H_ 6.72 (1H, d, *J* = 8.0 Hz, H-2’), 7.10 (1H, td, *J* = 8.0, 1.7 Hz, H-3’), 6.87 (1H, t, *J* = 8.0 Hz, H-4’), 7.06 (1H, dd, *J* = 8.0, 1.7 Hz, H-5’), 6.63 (1H, d, *J* = 9.0 Hz, H-7’), and 5.88 (1H, dd, *J* = 9.0, 3.7 Hz, H-8’); a 1,3,4-trisubstituted benzene ring at *δ*_H_ 7.62 (1H, d, *J* = 2.2 Hz, H-2), 7.39 (1H, dd, *J* = 8.4, 2.2 Hz, H-4), and 6.89 (1H, d, *J* = 8.4 Hz, H-5); one disubstituted *trans*-olefin at *δ*_H_ 7.90 (1H, d, *J* = 15.9 Hz, H-7) and 6.83 (1H, d, *J* = 15.9 Hz, H-8); an aldehyde proton at *δ*_H_ 9.61 (1H, d, *J* = 7.9 Hz, H-9); and one methine at *δ*_H_ 5.86 (1H, d, *J* = 3.7 Hz, H-9’). The connection C-9’/C-3 was determined by HMBC correlations from H-8’ (*δ*_H_ 5.88) to C-3 (*δ*_C_ 133.68) and from H-9’ (*δ*_H_ 5.86) to C-2 (*δ*_C_ 129.67) and C-4 (*δ*_C_ 133.06) (**Figure 6B**). As a result, the structure of **20** was determined as shown in **Figure 6A**.

Cassitamine C (**21**) was collected as a brown gum with the molecular formula of C_18_ H_14_O_4_ from the HRESIMS ion peak at m/z 317.0791 [M + Na]^+^ in positive ion mode (calcd. for C_18_H_14_O_4_Na, 317.0784). The ^1^H NMR data showed signals of a 1,3,4-trisubstituted benzene ring at *δ*_H_ 8.36 (1H, d, *J* = 2.2 Hz, H-2), 8.07 (1H, dd, *J* = 8.6, 2.2 Hz, H-4), and 7.03 (1H, d, *J* = 8.6 Hz, H-5); a 1,2-disubstituted benzene ring at *δ*_H_ 6.90 (1H, overlapped, H-2’), 7.27 (1H, td, *J* = 7.7, 1.6 Hz, H-3’), 6.92 (1H, td, *J* = 7.7, 1.1 Hz, H-4’), and 7.73 (1H, dd, *J* = 7.7, 1.6 Hz, H-5’); two disubstituted *trans*-olefins at [*δ*_H_ 7.97 (1H, d, *J* = 16.0 Hz, H-7) and 7.04 (1H, dd, *J* = 16.0, 7.9 Hz, H-8)] and [*δ*_H_ 8.14 (1H, d, *J* = 15.7 Hz, H-7’) and 7.87 (1H, d, *J* = 15.7 Hz, H-8’)]; and an aldehyde proton at *δ*_H_ 9.68 (1H, d, *J* = 7.9 Hz, H-9). The ^13^C NMR data exhibited 18 carbon signals, corresponding to one carbonyl group (*δ*_C_ 191.13), one formyl carbon (*δ*_C_ 197.02), and sixteen *sp²* carbons (*δ*_C_ 163.76, 158.98, 150.76, 142.08, 134.43, 133.10, 132.46, 131.41, 130.70, 130.55, 123.44, 122.86, 122.30, 121.02, 117.79, and 117.24). These spectral characteristics indicated that **21** contains two 2-hydroxyphenylacrylaldehyde moeties, confirmed by HMBC correlations from H-7 (*δ*_H_ 7.97) to C-9 (*δ*_C_ 197.02), C-2 (*δ*_C_ 132.46), and C-6 (*δ*_C_ 163.76), and from H-7’ (*δ*_H_ 8.14) to C-9’ (*δ*_C_ 191.13), C-1’ (*δ*_C_ 158.98), and C-5’ (*δ*_C_ 130.55) (**Figure 6B**). The connectivity of the two 2-hydroxyphenylacrylaldehyde moeties was established through HMBC correlations from H-2 (*δ*_H_ 8.36) and H-4 (*δ*_H_ 8.07) to C-9’ (*δ*_C_ 191.13) (**Figure 6B**). Consequently, the structure of **21** was determined as displayed in **Figure 6A**.

Cassitamine D (**22**) was obtained as a yellowish gum with the molecular formula of C _26_H_26_O_6,_ as determined by the HRESIMS ion peak at m/z 457.1623 [M + Na]^+^ in positive ion mode (calcd. for C_26_H_26_O_6_Na, 457.1622). The ^1^H NMR spectrum of **22** showed signals of a monosubstituted benzene ring (ring C) at *δ*_H_ 7.41 (2H, d, *J* = 7.3 Hz, H-2’’,6’’), 7.30 (2H, d, *J* = 7.3 Hz, H-3’’,5’’), and 7.24 (1H, overlapped, H-4’’); two 1,3,4-trisubstituted benzene rings (rings A, B) at *δ*_H_ [6.70 (1H, d, *J* = 1.9 Hz, H-2), 6.71 (1H, d, *J* = 8.1 Hz, H-5), and 7.62 (1H, dd, *J* = 8.1, 1.9 Hz, H-6)] and [6.36 (1H, d, *J* = 1.9 Hz, H-2’), 6.77 (1H, d, *J* = 8.1 Hz, H-5’), and 6.59 (1H, dd, *J* = 8.1, 1.9 Hz, H-6’)]; two methoxy groups at *δ*_H_ 3.76 (3H, s, H-3-OCH_3_) and 3.73 (3H, s, H-3’-OCH_3_); an oxymethylene at *δ*_H_ 4.29 (1H, dd, *J* = 11.5, 4.7 Hz, H-9a) and 4.07 (1H, t, *J* = 11.5 Hz, H-9b); one oxymethine at *δ*_H_ 4.73 (1H, d, *J* = 10.8 Hz, H-7); one dioxymethine at *δ*_H_ 5.46 (1H, d, *J* = 4.7 Hz, H-9’’); a methine at *δ*_H_ 3.07 (1H, td, *J* = 10.8, 4.7 Hz, H-8); and a disubstituted *trans*-olefin at *δ*_H_ 6.85 (1H, d, *J* = 16.1 Hz, H-7’’) and 6.31 (1H, dd, *J* = 16.1, 4.6 Hz, H-8’’). The 2D structure of **22** was further established by analyses of its 2D NMR data (**Figure 6B**). The HMBC correlations from H-7 (*δ*_H_ 4.73) to C-9 (*δ*_C_ 72.28), C-9’’ (*δ*_C_ 101.33), C-2 (*δ*_C_ 109.78), and C-6 (*δ*_C_ 120.72), from H-9’’ (*δ*_H_ 5.46) to C-7’’ (*δ*_C_ 133.80), C-9 (*δ*_C_ 72.28), and C-7 (*δ*_C_ 84.74); and from H-7’’ (*δ*_H_ 6.85) to C-2’’ (*δ*_C_ 127.12) indicated the presence of two phenylpropanoid moieties. The ring B was located at C-8 on the basis of the HMBC correlations from H-8 (*δ*_H_ 3.07) to C-2′ (*δ*_C_ 111.71) and C-6′ (*δ*_C_ 120.84). In addition, the two methoxy groups were assigned at C-3 and C-3′, respectively, by the HMBC correlations from *δ*_H_ 3.76 (3H, s, H-3-OCH_3_) to *δ*_C_ 146.38 (C-3) and from *δ*_H_ 3.73 (3H, s H-3′-OCH_3_) to *δ*_C_ 146.54 (C-3′). The locations of the methoxy groups were confirmed by NOESY experiment, which revealed NOE correlations between H-3-OCH_3_ (*δ*_H_ 3.76) and H-2 (*δ*_H_ 6.70), as well as H-3′-OCH_3_ (*δ*_H_ 3.73) and H-2′ (*δ*_H_ 6.36) (**Figure 6B**). The significant NOE correlations observed between H-9′′/H-7 and H-9*β* suggested that H-9′′, H-7, and H-9*β* are positioned along the axial bonds within the chair conformation of the dioxane ring, with *β*-orientations designated accordingly (**Figure 6B**). Furthermore, the coupling constant of 10.8 Hz between H-7 and H-8 supported the assignment of an *α*-axial orientation for H-8 (Cheng et al., 2014; R. Guo et al., 2019). Therefore, the relative configuration of **22** was proposed as *rel*-7*S*,8*S*,9′′*S* and its structure was shown in **Figure 6A**.

Cassitamine E (**23**) was yielded as a yellowish gum with the molecular formula of C_1 8_H_18_O_3_ established from the HRESIMS ion peak at m/z 305.1154 [M + Na]^+^ in positive ion mode (calcd. for C_18_H_18_O_3_Na, 305.1151). A comprehensive analysis of the NMR data (Table 1) revealed the presence of a *trans*-cinnamyl alcohol moiety in **23**, as evidenced by disubstituted *trans*-olefin protons at *δ*_H_ 6.65 (1H, dd, *J* = 16.0, 1.6 Hz, H-7) and 6.29 (1H, dt, *J* = 16.0, 6.1 Hz, H-8); one oxymethylen at *δ*_H_ 4.30 (2H, ddd, *J* = 12.3, 6.1, 1.6 Hz, H-9); and protons of a monosubstituted benzen ring at *δ*_H_ 7.40 (2H, d, *J* = 7.5 Hz, H-2,6), 7.35 (2H, d, *J* = 7.5 Hz, H-3,5), and 7.28 (1H, overlapped, H-4). The ^1^H−^1^H COSY correlations of H-7/H-8/H-9 and H-2,6/H-3,5/H-4, along with the key HMBC correlations from H-7 (*δ*_H_ 6.65) to C-2 (*δ*_C_ 126.55) and C-9 (*δ*_C_ 69.19), and from H-8 (*δ*_H_ 6.29) to C-1 (*δ*_C_ 136.31) (**Figure 6B**) confirmed the presence of a *trans*-cinnamyl alcohol moiety in **23**. Additionally, a 4-chromanol unit was identified on the basis of ^1^H−^1^H COSY correlations of H-7′/H-8′/H-9′ and H-3′/H-4′/H-5′/H-6′, and the HMBC correlations from H-5′ (*δ*_H_ 7.24) to C-9′ (*δ*_C_ 70.61) and from H-8′ (*δ*_H_ 2.64/2.18) to C-6′ (*δ*_C_ 119.58) (**Figure 6B**). Finally, the two moieties were linked via a C_9_-O-C_9′_ bond as confirmed by HMBC correlation between H-9 (*δ*_H_ 4.30) and C-9′ (*δ*_C_ 70.61) (**Figure 6B**). The NOESY cross-peak between H-7′ and H-9′ indicated their spatial proximity, suggesting that they are cofacial. Hence, the relative configuration of **23** was proposed as *rel*-7′*R*,9′*R* and its structure was illustrated in **Figure 6A**.

Cassitamine F (**24**) was isolated as a yellowish gum, and its molecular formula of C_18_ H_16_O_2_ was determined from the HRESIMS ion peak at m/z 287.1046 [M + Na]^+^ in positive ion mode (calcd. for C_18_H_16_O_2_Na, 287.1043). The ^1^H NMR spectrum displayed signals of two monosubstituted benzene rings at *δ*_H_ [7.44 (2H, d, *J* = 7.5 Hz, H-2,6), 7.37 (2H, t, *J* = 7.5 Hz, H-3,5), and 7.29 (1H, overlapped, H-4)] and [7.52 (2H, dd, *J* = 7.6, 1.6 Hz, H-2′,6′), 7.41 (2H, t, *J* = 7.6 Hz, H-3′,5′), and 7.40 (1H, overlapped, H-4′)]; one disubstituted *trans*-olefin at *δ*_H_ 7.12 (1H, d, *J* = 15.1 Hz, H-7′) and 7.41 (1H, overlapped, H-8′); one trisubstituted olefin at *δ*_H_ 7.15 (1H, d, *J* = 14.8 Hz, H-9′); one oxymethine at *δ*_H_ 5.98 (1H, s, H-7); and an aldehyde proton at *δ*_H_ 9.52 (1H, d, *J* = 1.2 Hz, H-9). The 2D structure of **24** was supported by the HMBC correlations from H-7 (*δ*_H_ 5.98) to C-6 (*δ*_C_ 125.59), C-9 (*δ*_C_ 195.30), and C-9′ (*δ*_C_ 150.43), from H-9 (*δ*_H_ 9.52) to C-8 (*δ*_C_ 140.36) and C-9′, and from H-7′ (*δ*_H_ 7.12) to C-2′ (*δ*_C_ 127.86), and by the ^1^H−^1^H COSY correlations of H-7′/H-8′/H-9′ (**Figure 6B**). Thus, the structure of **24** was determined as shown in **Figure 6A**.

#### Bioactivity Screening of Isolated Compounds

To identify the bioactive compounds responsible for the autophagy-inducing and anti-inflammatory effects of *C. cassia*, we conducted LC3-HiBiT reporter assays to assess autophagy activation and HEK-TLR4 reporter assays to evaluate TLR4-mediated inflammatory signaling (**Figure 7A-B**). In the LC3-HiBiT assay, (E)-3-(2-Hydroxyphenyl)propenal, Cinnaphenylpropanoid M, Cassitamine B, Cassitamine C, and Cassitamine F significantly increased luminescence, indicating strong autophagy induction (**Figure 7A**). In the HEK-TLR4 reporter assay, (E)-3-(2-Hydroxyphenyl)propenal, Cassitamine C, Cassitamine E, and Cassitamine F significantly inhibited SEAP activity, indicating suppression of TLR4 activation (**Figure 7B**). Interestingly, Cassitamine C and Cassitamine F displayed dual activity, both promoting autophagy and inhibiting TLR4 activation, highlighting their potential as multifunctional bioactive agents that contribute to both autophagy modulation and inflammatory suppression..

**Figure 7.**
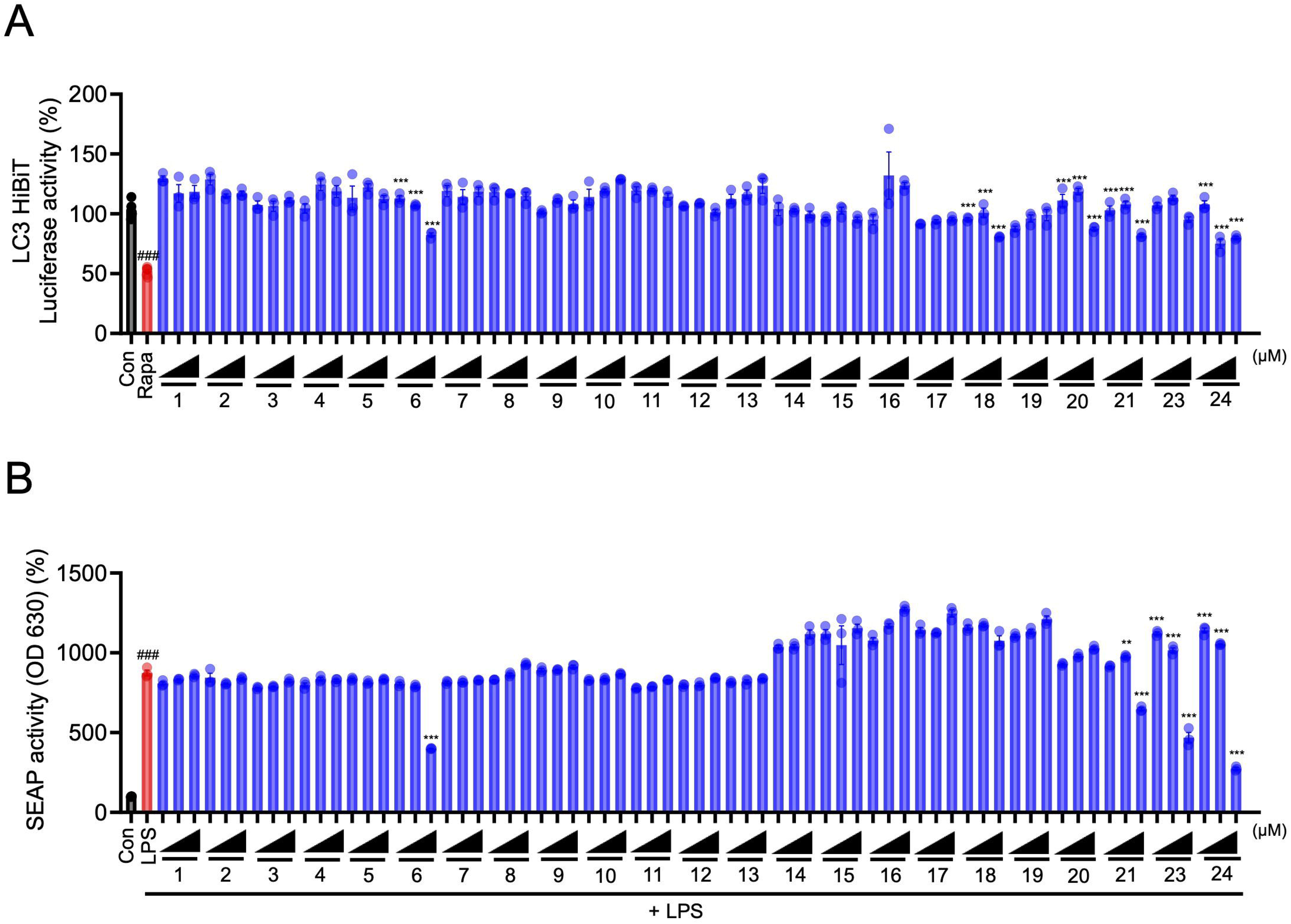
Bioactivity screening of isolated compounds from *C. cassia* extract. (A) LC3-HiBiT reporter assay was used to identify autophagy-inducing compounds. Luminescence (relative light units, RLU) was measured to assess LC3 reporter activity. (B) HEK-TLR4 reporter assay was used to evaluate anti-inflammatory activity. TLR4 activation was assessed by monitoring secreted embryonic alkaline phosphatase (SEAP) activity. Data are presented as mean ± SEM. Significant differences at ###p < 0.001 versus the Control group, ***p < 0.001, **p < 0.01 versus the Vehicle group.

### 3.7. Anti-Inflammatory Effects of Cinassia F in an LPS-Induced Sepsis Model

To further evaluate the in vivo anti-inflammatory potential of the bioactive compounds, mice were subjected to LPS-induced septic shock and treated with Cinassia C, E, or F. Serum cytokine levels of TNF-α, IFN-γ, IL-6, IL-17A, IL-27, and MCP-1 were measured (**Figure 8A-F**). The results demonstrated that Cassitamine F significantly reduced the levels of pro-inflammatory cytokines, suggesting its potent anti-inflammatory activity. In contrast, Cassitamine C and Cassitamine E did not exhibit significant effects on cytokine levels, indicating that Cassitamine F is the key compound responsible for the in vivo anti-inflammatory effects observed in EECE.

**Figure 8.**
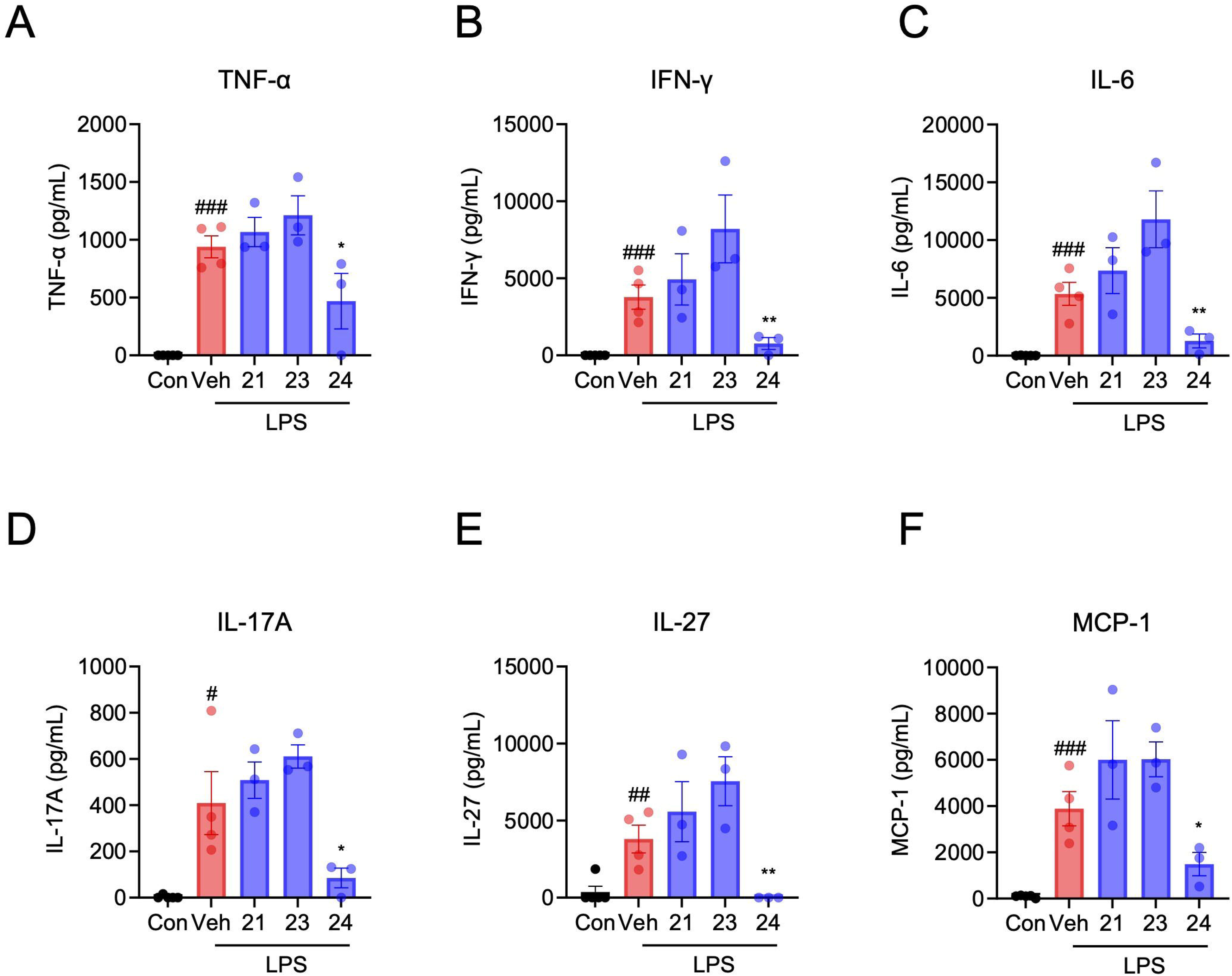
Anti-inflammatory effects of Cinassia E in an LPS-induced sepsis model. Mice were subjected to LPS-induced septic shock and treated with Cinassia B, D, or E to evaluate their anti-inflammatory potential. Serum cytokine levels of (A) TNF-α, (B) IFN-γ, (C) IL-6, (D) IL-17A, (E) IL-27, (F) MCP-1 were measured. Data are presented as mean ± SEM. Significant differences at ###p < 0.001, ##p < 0.01, #p < 0.05 versus the Control group, **p < 0.01, *p < 0.05 versus the Vehicle group.

## 4. Discussion

In this study, we identified EECC as a potent inducer of autophagy and a suppressor of inflammation. High-throughput screening of 1,679 natural product extracts revealed that EECC significantly enhanced autophagy, as confirmed by LC3 HiBiT reporter assays, fluorescence microscopy of RFP-GFP-LC3 fusion protein, and increased LC3-II and Beclin1 expression. Additionally, EECC suppressed TLR activation across multiple pathways and significantly reduced IL-6 and TNF-α productions in LPS-stimulated macrophages, mediated through suppression of NF-κB/AP1 and IRF pathways. The anti-inflammatory effects were autophagy-dependent, as inhibition of autophagy restored NF-κB activation. Bioactivity-guided fractionation led to the identification of 24 compounds, including six novel structures (Cassitamine A-F), among which Cassitamine F exhibited the strongest dual activity as an autophagy inducer and anti-inflammatory agent. In an LPS-induced sepsis model, Cassitamine F significantly reduced pro-inflammatory cytokine levels, highlighting it as a promising therapeutic candidate for inflammatory diseases like sepsis.

Previous studies have shown that cinnamaldehyde, a key component of *C. cassia*, induces autophagy in cancer cells and modulates NF-κB and PERK-CHOP pathways (J. Guo et al., 2024; Kim, 2022), focusing on autophagy-mediated tumor cell death. Our findings confirm that *C. cassia*-derived compounds regulate autophagy but extend this knowledge by demonstrating that autophagy also plays a key role in inflammation suppression. Unlike previous studies that linked cinnamaldehyde-induced autophagy to tumor suppression, our study provides novel insights into the anti-inflammatory effects of autophagy induction. We demonstrated that EECC not only induces autophagy but also suppresses NF-κB/AP1 and IRF pathways in macrophages, leading to a reduction in pro-inflammatory cytokines.

Previous studies have established the anti-inflammatory properties of *C. cassia*, particularly through essential oils, ethanol extracts, and polysaccharides, which have demonstrated protective effects in lung inflammation, gastrointestinal disorders, and diabetes (Jia et al., 2023; F. Liu, Yang, Dong, Zhu, Feng, & Wu, 2024; Shin et al., 2017). These studies demonstrated that various preparations from *C. cassia* effectively suppressed inflammation by modulating immune responses, primarily via the inhibition of NF-κB signaling pathways and reducing cytokine production. Our findings align well with these observations but extend existing knowledge by clearly demonstrating that EECC-mediated anti-inflammatory effects are closely linked to its ability to induce autophagy.

Furthermore our study specifically identified bioactive constituents through bioactivity-guided fractionation, isolating 24 compounds and characterizing six novel molecules (Cassitamine A– F). Notably, we identified Cassitamine F as a potent dual-functional compound that simultaneously enhances autophagy and exhibits substantial anti-inflammatory activity in an LPS-induced sepsis model. Thus, our study broadens the therapeutic scope of *C. cassia*, identifying novel compounds with immunomodulatory activity, particularly Cassitamine F, a promising candidate for inflammatory diseases like sepsis.

In this study, we identified six novel chemical constituents (Cassitamine A–F) from EECC through comprehensive spectroscopic analyses (NMR, HRESIMS, COSY, HMBC, NOESY). These novel compounds exhibit diverse structural features, including phenylpropanoid frameworks, chromene derivatives, aldehyde functionalities, and unique dioxane and chromanol moieties. Cassitamine A, identified as cinnamic acid 2-oxo-2-phenylethyl ester, contains two monosubstituted phenyl groups linked through a trans-olefin and carbonyl functionalities. Cassitamine B and C exhibit chromene and aldehyde groups, respectively, connected to substituted aromatic rings via characteristic olefinic bonds. Cassitamine D is particularly distinctive, with a dioxane ring structure displaying clear stereochemical orientation established by HMBC and NOESY correlations. Cassitamine E comprises a trans-cinnamyl alcohol linked to a 4-chromanol unit, whereas Cassitamine F features two phenyl groups connected via olefinic and aldehyde functionalities. Such structural diversity among these novel isolates not only expands the chemical profile of EECC but also provides new opportunities for the discovery and development of bioactive agents targeting autophagy and inflammatory pathways.

Our findings offer important mechanistic insights into how EECC and its bioactive compounds, particularly Cassitamine F, exert their immunomodulatory effects by inducing autophagy and subsequently modulating inflammatory signaling pathways. Specifically, we observed that EECC significantly induced autophagy, as indicated by changes in autophagic markers LC3-II, Beclin1, and p62. Previous studies have reported that autophagy mediates the degradation of critical NF-κB signaling components, such as IKKβ and NF-κB-inducing kinase (NIK), thereby negatively regulating inflammatory signaling (Trocoli & Djavaheri-Mergny, 2011). Additionally, recent findings highlight that activating selective autophagy, such as mitophagy, can reduce systemic inflammation by removing damaged mitochondria and mitigating oxidative stress (Silwal et al., 2023). Our data support these mechanisms by demonstrating EECC’s suppression of NF-κB/AP1 and IRF pathways in an autophagy-dependent manner, suggesting that EECC and Cassitamine F may similarly promote autophagic clearance of inflammatory signaling components or damaged organelles, consequently preventing excessive inflammation.

Although this study provides significant insights into the anti-inflammatory effects of EECC and Cassitamine F, several limitations should be noted. First, our work identified key autophagy markers (LC3-II, Beclin1, p62), but detailed signaling mechanisms involving specific autophagy regulators, such as mTOR, ULK1, and Beclin1-associated complexes, were not fully elucidated. Moreover, the precise molecular targets and detailed signaling pathways mediating Cassitamine F’s dual autophagy-inducing and anti-inflammatory effects remain to be characterized. Another practical limitation was the insufficient yield of isolated novel compounds, which restricted the scope of experiments and hampered further mechanistic and pharmacological evaluations. Future research addressing these limitations will strengthen the mechanistic understanding and therapeutic potential of EECC and Cassitamine F in inflammatory diseases.

## 5. Conclusion

Our study demonstrates that C. cassia extract and its compound Cassitamine F show anti-inflammatory effects by inducing autophagy. By influencing inflammatory pathways such as TLR, NF-κB, and IRF signaling, EECC reduces cytokine production through an autophagy-dependent mechanism. These results indicate that EECC and Cassitamine F could have therapeutic potential for inflammatory conditions like sepsis, although further studies are needed to confirm its clinical relevance.

## Supporting information on new identified compounds derived from *C. cassia* is provided (Fig. S.1.1. - Fig. S.6.8.)

### CRediT authorship contribution statement

**Geon Park**: Conceptualization, Methodology, Investigation, Visualization, Writing – original draft. **Tam Thi Le**: Conceptualization, Methodology, Visualization. **Tae Kyeom Kang**: Methodology, Resources. **Yuna Jung**: Methodology, Resources. **Wook-Bin Lee**: Conceptualization, Supervision, Writing – original draft, Writing – review & editing. **Sang Hoon Jung**: Conceptualization, Supervision, Writing – review & editing.

## Acknowledgement

This work was supported by an intramural grant (2E33531) from the Korea Institute of Science and Technology (KIST).

## Data Availability Statement

The data that support the findings of this study are available in paper or the supplementary material of this article.

## References

Angus, D. C., & Van der Poll, T. (2013). Severe sepsis and septic shock. New England journal of medicine, 369(9), 840–851.

Chan, Y. Y., Wu, C. H., Wu, S. J., & Wu, T. S. (2002). The constituents and synthesis of cryptamygin-A from the stem bark of. Journal of the Chinese Chemical Society, 49(2), 263–268. <GO to ISI>://WOS:000176991000018.

Cheng, Z. B., Lu, X., Bao, J. M., Han, Q. H., Dong, Z., Tang, G. H., . . . Yin, S. (2014). (±)-Torreyunlignans A–D, Rare 8–9′ Linked Neolignan Enantiomers as Phosphodiesterase-9A Inhibitors from Torreya yunnanensis. Journal of Natural Products, 77(12), 2651–2657. 10.1021/np500528u.

Cohen, J. (2002). The immunopathogenesis of sepsis. Nature, 420(6917), 885–891.

Deretic, V. (2021). Autophagy in inflammation, infection, and immunometabolism. Immunity, 54(3), 437–453.

Deretic, V., & Levine, B. (2018). Autophagy balances inflammation in innate immunity. Autophagy, 14(2), 243–251.

Deretic, V., Saitoh, T., & Akira, S. (2013). Autophagy in infection, inflammation and immunity. Nature Reviews Immunology, 13(10), 722–737.

Dikic, I., & Elazar, Z. (2018). Mechanism and medical implications of mammalian autophagy. Nature reviews Molecular cell biology, 19(6), 349–364.

Duan, L., Rao, X., & Sigdel, K. R. (2019). Regulation of inflammation in autoimmune disease. Journal of immunology research, 2019, 7403796.

Fan, H. X., Huang, G. F., Guo, Q., Ma, J. H., Huang, Y. J., Huang, S. X., . . . Gao, H. (2024). Bioactive Phenylpropanoid Glycosides, Dimers, and Heterodimers from the Bark of Cinnamomum cassia (L.) J.Presl. Journal of Agricultural and Food Chemistry, 72(29), 16263–16275. 10.1021/acs.jafc.4c02129.

Fang, F., Sang, S. M., Chen, K. Y., Gosslau, A., Ho, C. T., & Rosen, R. T. (2005). Isolation and identification of cytotoxic compounds from Bay leaf (Laurus nobilis). Food Chemistry, 93(3), 497–501. 10.1016/j.foodchem.2004.10.029.

Fernandes, R. A., Ranjan, R. S., & Choudhary, P. (2024). K2S2O8-Mediated or Azobisisobutyronitrile-Catalyzed Regioselective Aerobic Oxidative Cleavage of 1-Arylbutadienes to Cinnamaldehydes. Organic Letters, 26(29), 6247–6252. 10.1021/acs.orglett.4c02241.

Guo, J., Yan, S., Jiang, X., Su, Z., Zhang, F., Xie, J., . . . Yao, C. (2024). Advances in pharmacological effects and mechanism of action of cinnamaldehyde. Frontiers in Pharmacology, 15, 1365949.

Guo, R., Shang, X. Y., Lv, T. M., Yao, G. D., Lin, B., Wan, X. B., . . . Song, S. J. (2019). Phenylpropanoid derivatives from the fruit of Crataegus pinnatifida Bunge and their distinctive effects on human hepatoma cells. Phytochemistry, 164, 252–261. 10.1016/j.phytochem.2019.05.005.

Hayashi, K., Tanimoto, H., Zhang, H., Morimoto, T., Nishiyama, Y., & Kakiuchi, K. (2012). Efficient Synthesis of α,β-Unsaturated Alkylimines Performed with Allyl Cations and Azides: Application to the Synthesis of an Ant Venom Alkaloid. Organic Letters, 14(22), 5728–5731. 10.1021/ol302608q.

Ho, J., Yu, J., Wong, S. H., Zhang, L., Liu, X., Wong, W. T., . . . Gin, T. (2016). Autophagy in sepsis: degradation into exhaustion? Autophagy, 12(7), 1073–1082.

Hong, J.-W., Yang, G.-E., Kim, Y. B., Eom, S. H., Lew, J.-H., & Kang, H. (2012). Anti-inflammatory activity of cinnamon water extract in vivo and in vitro LPS-induced models. BMC complementary and alternative medicine, 12, 1–8.

Jia, W., He, X., Jin, W., Gu, J., Yu, S., He, J., . . . Yang, L. (2023). Ramulus Cinnamomi essential oil exerts an anti-inflammatory effect on RAW264. 7 cells through N-acylethanolamine acid amidase inhibition. Journal of Ethnopharmacology, 317, 116747.

Kang, T. K., Le, T. T., Kwon, H., Park, G., Kim, K.-A., Ko, H., . . . Jung, S. H. (2023). Lithospermum erythrorhizon Siebold & Zucc. extract reduces the severity of endotoxin-induced uveitis. Phytomedicine, 121, 155133.

Khamarui, S., Maiti, R., & Maiti, D. K. (2015). General base-tuned unorthodox synthesis of amides and ketoesters with water. Chemical Communications, 51(2), 384–387. 10.1039/c4cc07961b.

Kim, T. W. (2022). Cinnamaldehyde induces autophagy-mediated cell death through ER stress and epigenetic modification in gastric cancer cells. Acta Pharmacologica Sinica, 43(3), 712–723.

Kimura, S., Noda, T., & Yoshimori, T. (2007). Dissection of the autophagosome maturation process by a novel reporter protein, tandem fluorescent-tagged LC3. Autophagy, 3(5), 452–460.

Kiyotsuka, Y., Katayama, Y., Acharya, H. P., Hyodo, T., & Kobayashi, Y. (2009). New General Method for Regio- and Stereoselective Allylic Substitution with Aryl and Alkenyl Coppers Derived from Grignard Reagents. Journal of Organic Chemistry, 74(5), 1939–1951. 10.1021/jo802426g.

Kuo, Y. H., & Yu, M. T. (1996). Two new sesquiterpenes, (-)-15-hydroxycalamenene and (-)- 1-hydroxy-1,3,5-bisabolatrien-10-one, from the heartwood of Juniperus formosana Hay var concolor Hay. Chemical & Pharmaceutical Bulletin, 44(11), 2150–2152. <GO to ISI>://WOS:A1996VV31300027.

Li, C.-y., Liao, L.-j., Yang, S.-x., Wang, L.-y., Chen, H., Luo, P., . . . Huang, Y.-Q. (2024). Cinnamaldehyde: An effective component of Cinnamomum cassia inhibiting Helicobacter pylori. Journal of Ethnopharmacology, 330, 118222.

Li, X., Lu, H.-Y., Jiang, X.-W., Yang, Y., Xing, B., Yao, D., . . . Zhao, Q.-C. (2021). Cinnamomum cassia extract promotes thermogenesis during exposure to cold via activation of brown adipose tissue. Journal of Ethnopharmacology, 266, 113413.

Lim, Y., Kang, T. K., Kim, M. I., Kim, D., Kim, J. Y., Jung, S. H., . . . Seo, M. H. (2025). Massively Parallel Screening of Toll/Interleukin-1 Receptor (TIR)-Derived Peptides Reveals Multiple Toll-Like Receptors (TLRs)-Targeting Immunomodulatory Peptides. Advanced Science, 12(1), 2406018.

Liu, F., Yang, Y., Dong, H., Zhu, Y., Feng, W., & Wu, H. (2024). Essential oil from Cinnamomum cassia Presl bark regulates macrophage polarization and ameliorates lipopolysaccharide-induced acute lung injury through TLR4/MyD88/NF-κB pathway. Phytomedicine, 129, 155651.

Liu, X., Fu, J., Yao, X. J., Yang, J., Liu, L., Xie, T. G., . . . Zhu, G. Y. (2018). Phenolic Constituents Isolated from the Twigs of Cinnamomum cassia and Their Potential Neuroprotective Effects. Journal of Natural Products, 81(6), 1333–1342. 10.1021/acs.jnatprod.7b00924.

Loos, B., Toit, A. d., & Hofmeyr, J.-H. S. (2014). Defining and measuring autophagosome flux—concept and reality. Autophagy, 10(11), 2087–2096.

Malik, P., & Chakraborty, D. (2010). Bismuth(III) Oxide Catalyzed Oxidation of Alcohols with tert-Butyl Hydroperoxide. Synthesis(21), 3736–3740. 10.1055/s-0030-1258221.

Matsuoka, T., Hattori, A., Oishi, S., Araki, M., Ma, B., Fujii, T., . . . Inuki, S. (2023). Establishment of an MR1 Presentation Reporter Screening System and Identification of Phenylpropanoid Derivatives as MR1 Ligands. Journal of Medicinal Chemistry, 66(17), 12520–12535. 10.1021/acs.jmedchem.3c01122.

Matsuzawa-Ishimoto, Y., Hwang, S., & Cadwell, K. (2018). Autophagy and inflammation. Annual review of immunology, 36(1), 73–101.

Mcdonald, T. R., Turner, J. A., Gabbey, A. L., Balasubramanian, P., & Rousseaux, S. A. L. (2024). Synthesis of Borylated (Aminomethyl)cyclopropanes Using C-Bisnucleophiles. Organic Letters, 26(18), 3822–3827. 10.1021/acs.orglett.4c00987.

Mizushima, N., & Levine, B. (2020). Autophagy in human diseases. New England journal of medicine, 383(16), 1564–1576.

Ododo, M. M., Choudhury, M. K., & Dekebo, A. H. (2016). Structure elucidation of β-sitosterol with antibacterial activity from the root bark of Malva parviflora. Springerplus, 5. https://doi.org/ARTN 1210 10.1186/s40064-016-2894-x.

Riga, R., Happyana, N., Quentmeier, A., Zammarelli, C., Kayser, O., & Hakim, E. H. (2021). Secondary metabolites from Diaporthe lithocarpus isolated from Artocarpus heterophyllus. Natural Product Research, 35(14), 2324–2328. 10.1080/14786419.2019.1672685.

Saitoh, T., & Akira, S. (2010). Regulation of innate immune responses by autophagy-related proteins. Journal of Cell Biology, 189(6), 925–935.

Satmbekova, D., Srivedavyasasri, R., Orazbekov, Y., Omarova, R., Datkhayev, U., & Ross, S. A. (2018). Chemical and biological studies on Cichorium intybus L. Natural Product Research, 32(11), 1343–1347. 10.1080/14786419.2017.1343319.

Shibutani, S. T., Saitoh, T., Nowag, H., Münz, C., & Yoshimori, T. (2015). Autophagy and autophagy-related proteins in the immune system. Nature immunology, 16(10), 1014–1024.

Shin, W.-Y., Shim, D.-W., Kim, M.-K., Sun, X., Koppula, S., Yu, S. H., . . . Lee, K.-H. (2017). Protective effects of Cinnamomum cassia (Lamaceae) against gout and septic responses via attenuation of inflammasome activation in experimental models. Journal of Ethnopharmacology, 205, 173–177.

Silwal, P., Kim, Y. J., Lee, Y. J., Kim, I. S., Jeon, S. M., Roh, T., . . . Jo, D. S. (2023). Chemical mimetics of the N-degron pathway alleviate systemic inflammation by activating mitophagy and immunometabolic remodeling. Experimental & Molecular Medicine, 55(2), 333–346.

Trapp, M. A., Kai, M., Mithöfer, A., & Rodrigues, E. (2015). Antibiotic oxylipins from Alternanthera brasiliana and its endophytic bacteria. Phytochemistry, 110, 72–82. 10.1016/j.phytochem.2014.11.005.

Trocoli, A., & Djavaheri-Mergny, M. (2011). The complex interplay between autophagy and NF-κB signaling pathways in cancer cells. American journal of cancer research, 1(5), 629.

Xiang, Y., Zhang, M., Jiang, D., Su, Q., & Shi, J. (2023). The role of inflammation in autoimmune disease: a therapeutic target. Frontiers in immunology, 14, 1267091.

Yan, H. M., Song, Y., Chai, B., Shi, Q. Y., Zhang, Z. X., Li, F. F., . . . Yu, S. S. (2023). Analgesic sesquiterpenes from the roots of Rhododendron micranthum. Tetrahedron, 141. https://doi.org/ARTN 133436 10.1016/j.tet.2023.133436.

Zhang, C., Fan, L., Fan, S., Wang, J., Luo, T., Tang, Y., . . . Yu, L. (2019). Cinnamomum cassia Presl: A review of its traditional uses, phytochemistry, pharmacology and toxicology. Molecules, 24(19), 3473.

