## Supporting information 1.1. - 6.8. for "*Cinnamomum cassia* Extract and Its Novel Isolated Compound Suppress Inflammation via Autophagy Induction in Sepsis"

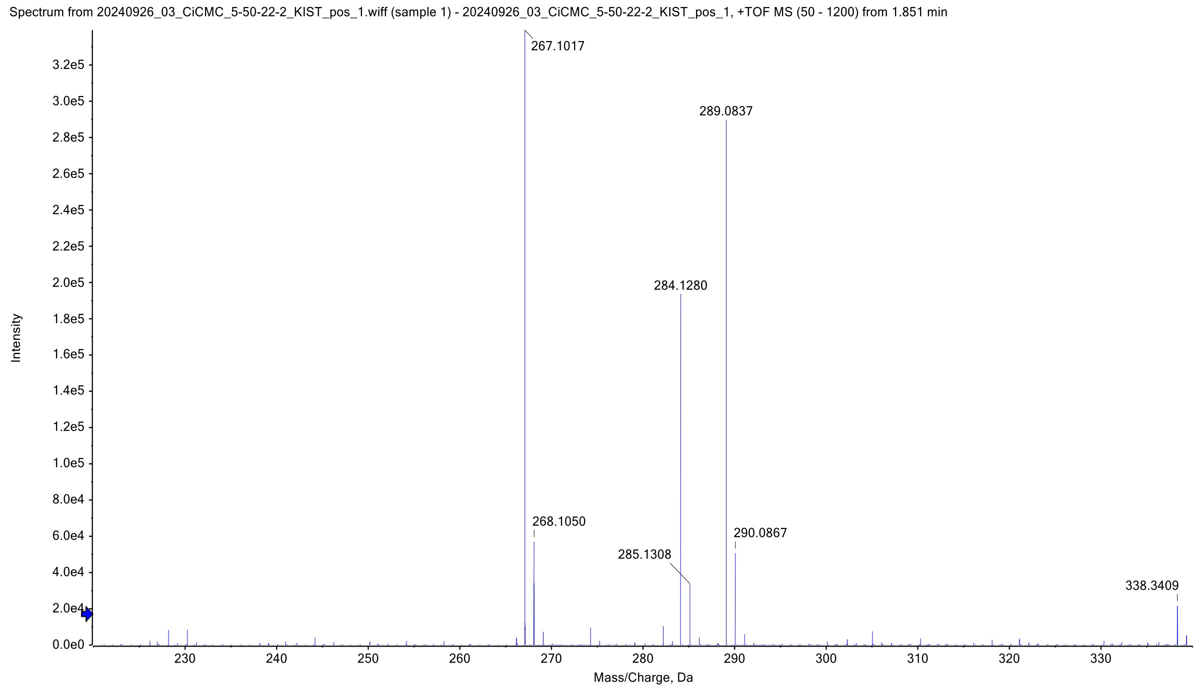

Fig. S.1.1. HR ESI-MS spectrum of **19**

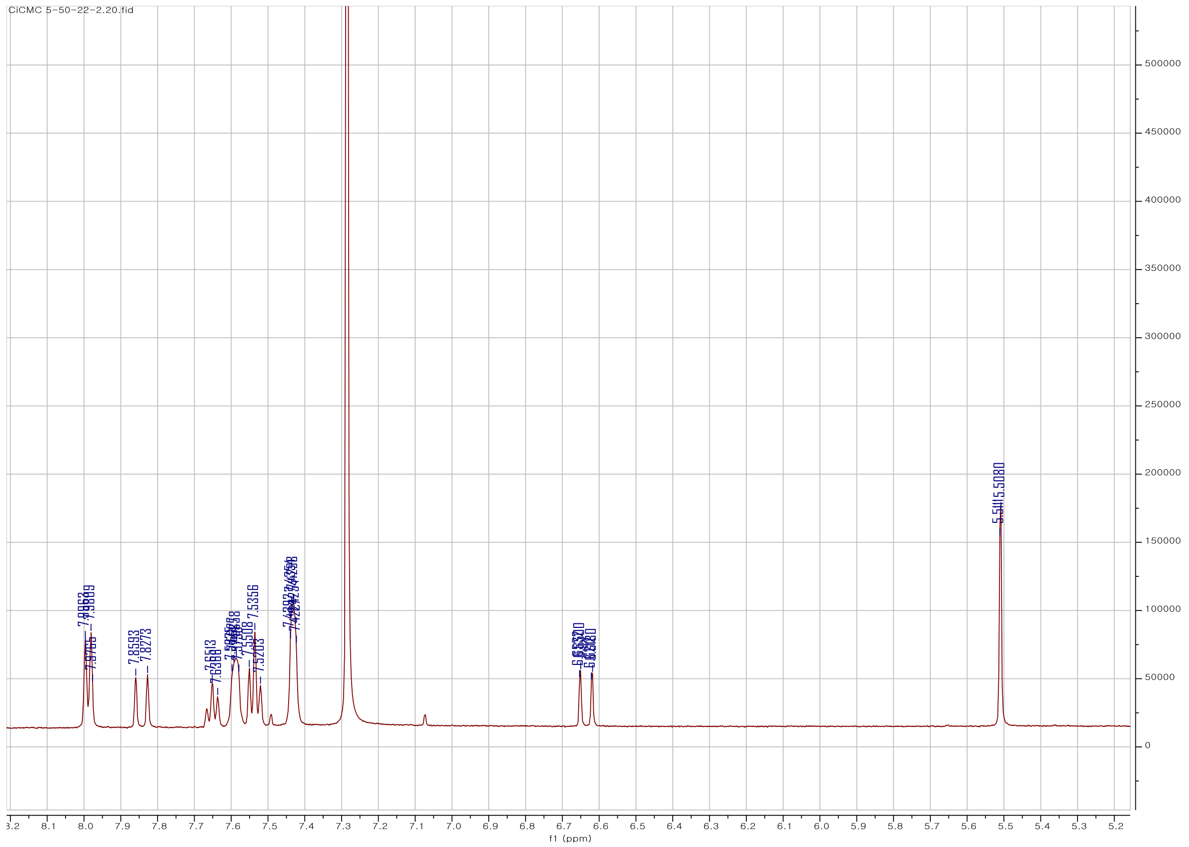

Fig. S.1.2. $^{1}$H-NMR spectrum of **19** in $CHCl_3$-*d* at 500 MHz

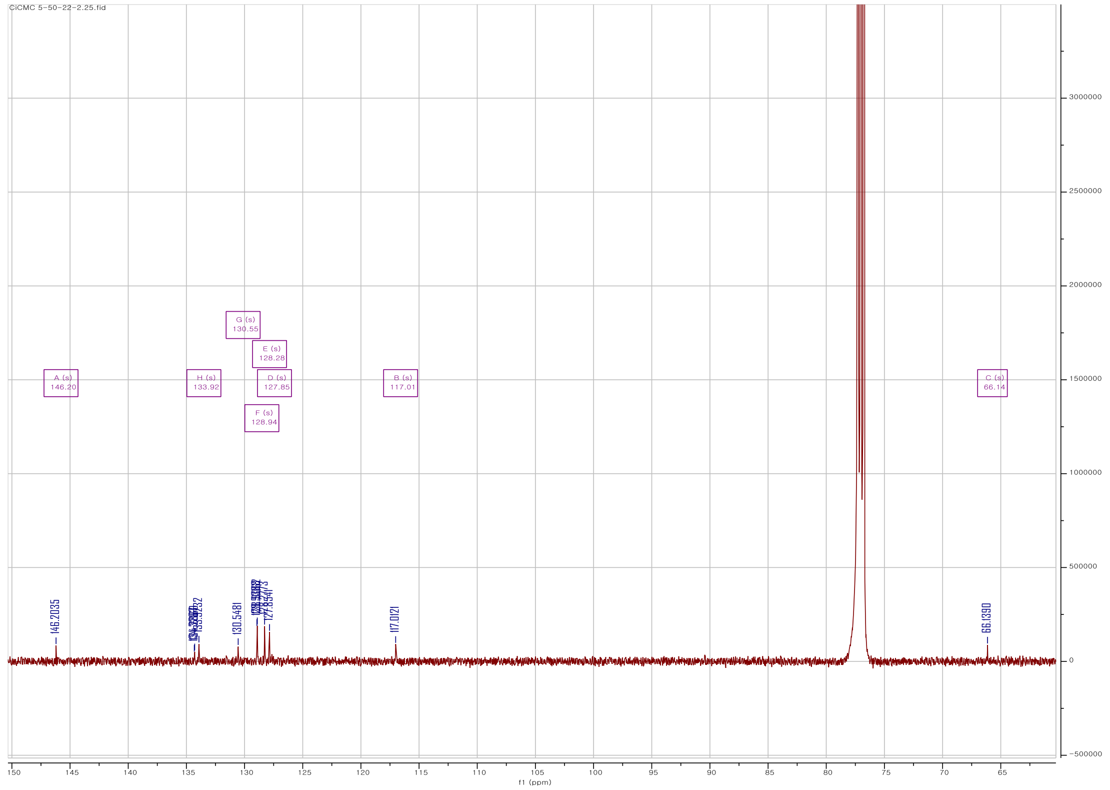

Fig. S.1.3. $^{13}$C-NMR spectrum of **19** in $CHCl_3$-*d* at 125 MHz

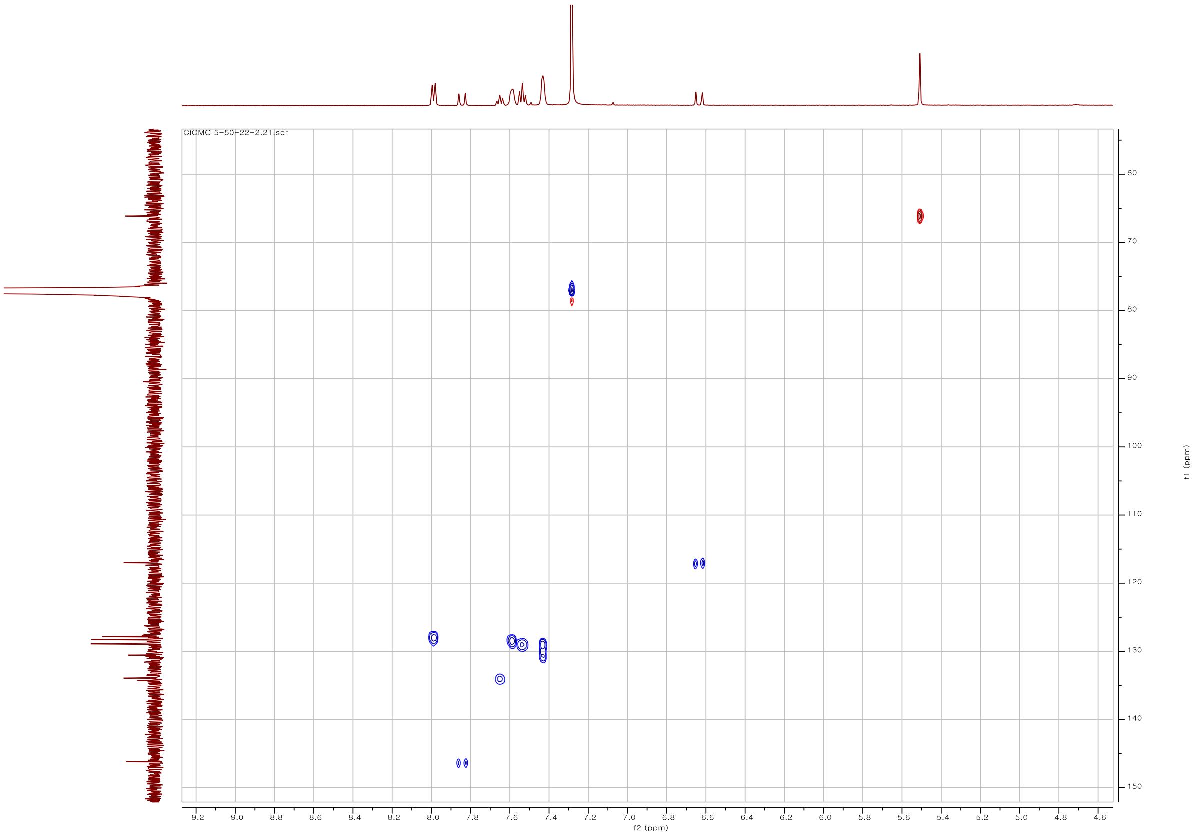

Fig. S.1.4. HSQC spectrum of **19** in $CHCl_3$-*d*

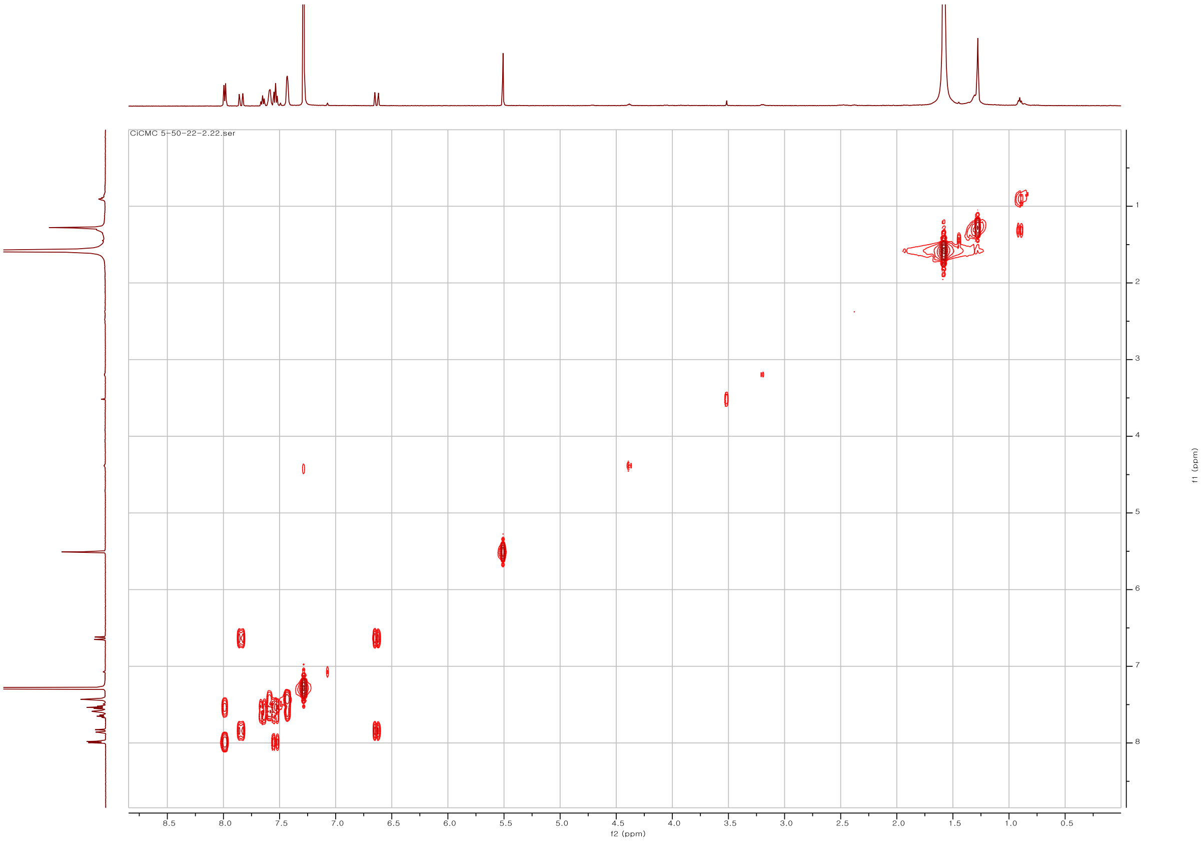

Fig. S.1.5. COSY spectrum of **19** in $CHCl_3$-*d*

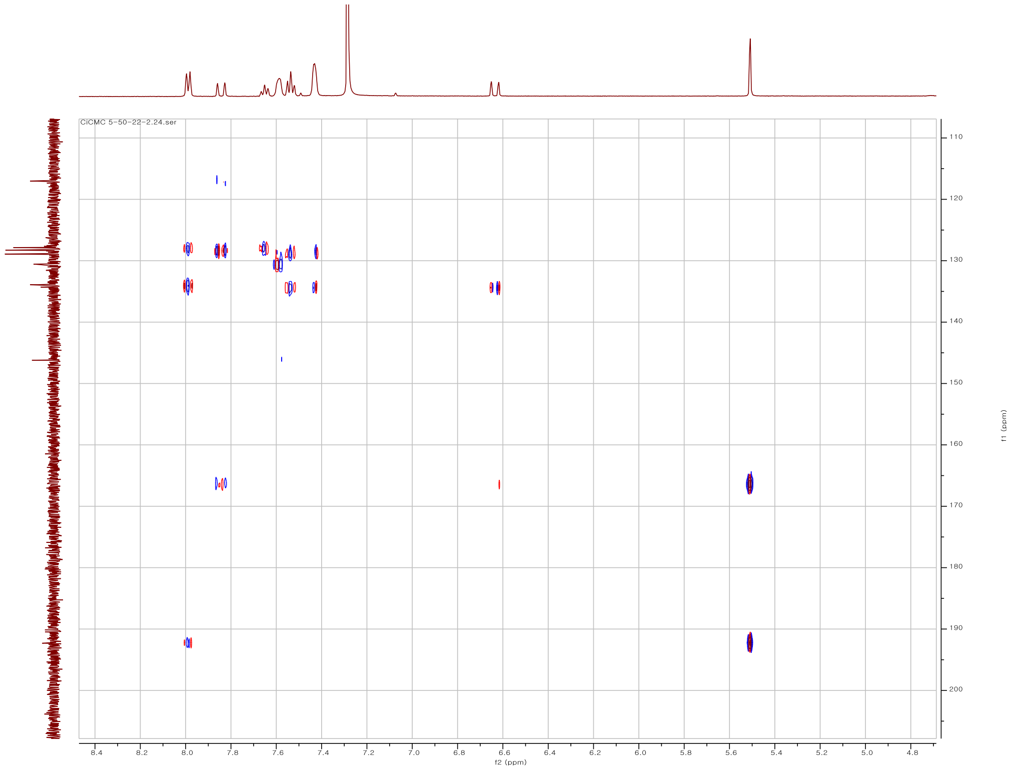

Fig. S.1.6. HMBC spectrum of **19** in $CHCl_3$-*d*

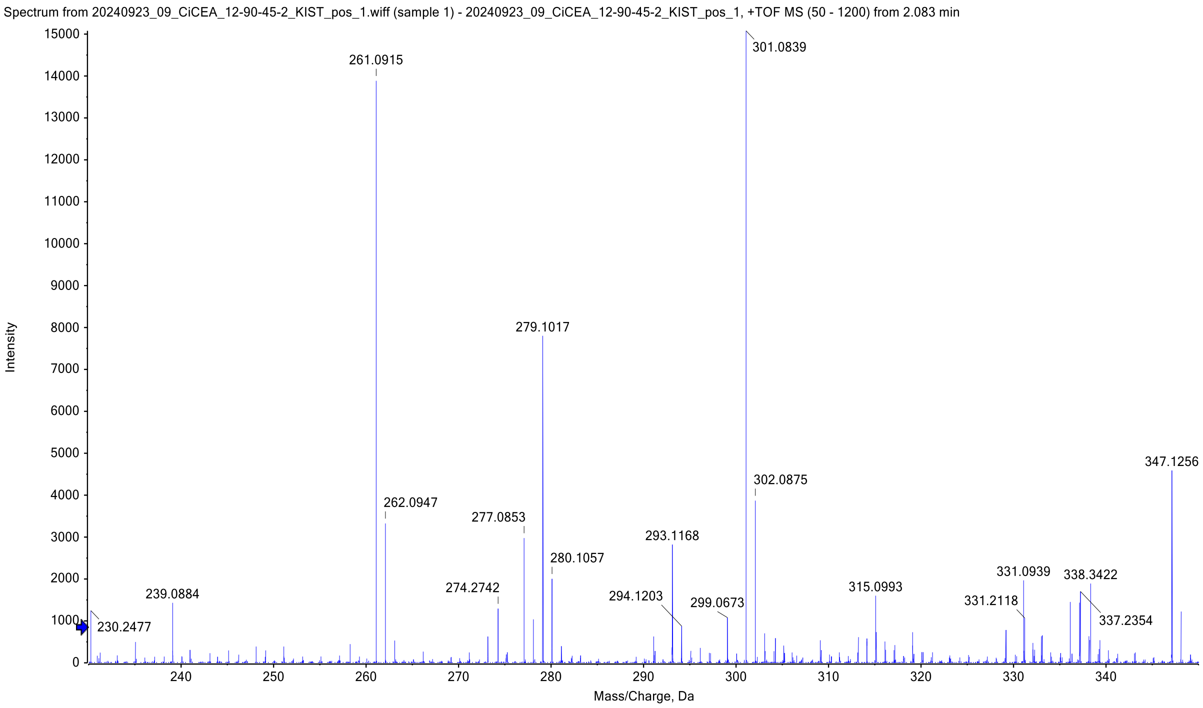

Fig. S.2.1. HR ESI-MS spectrum of **20**

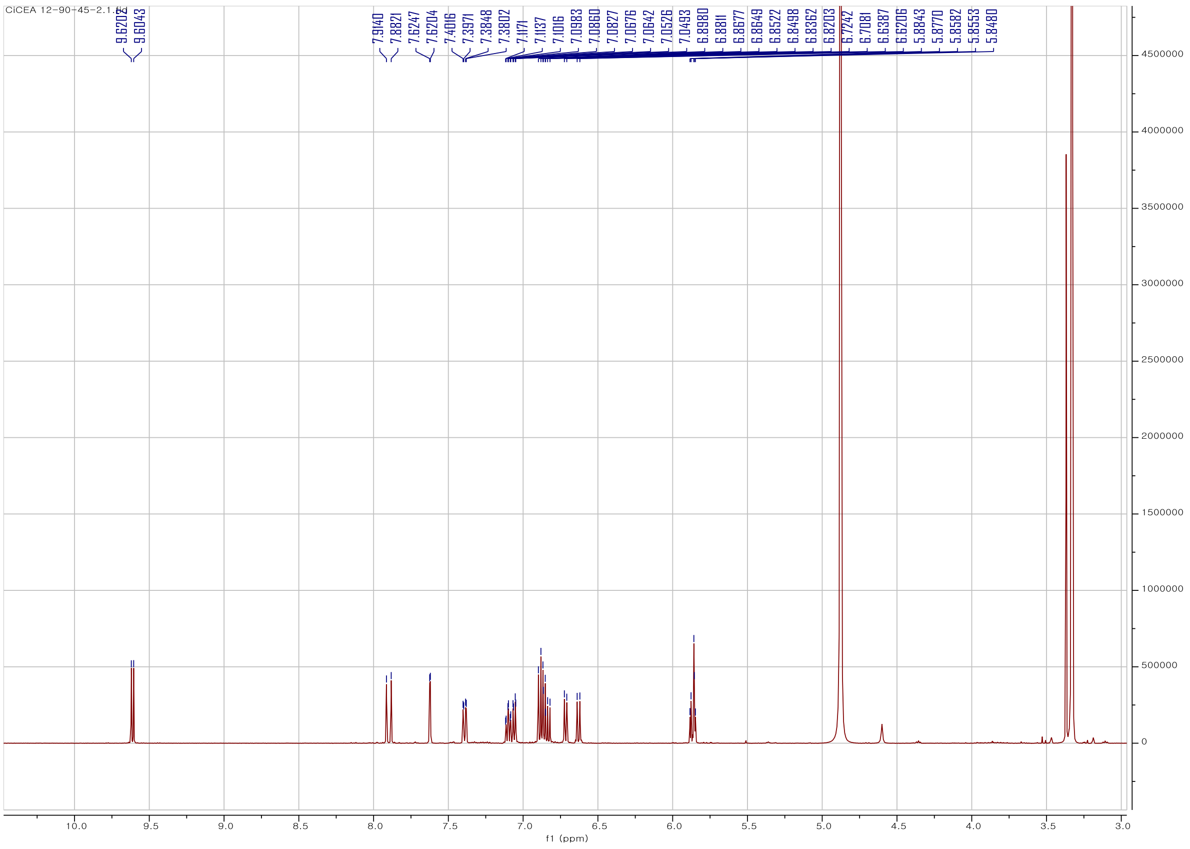

Fig. S.2.2. [1]H-NMR spectrum of **20** in $CH_3OH$-$d_4$ at 500 MHz

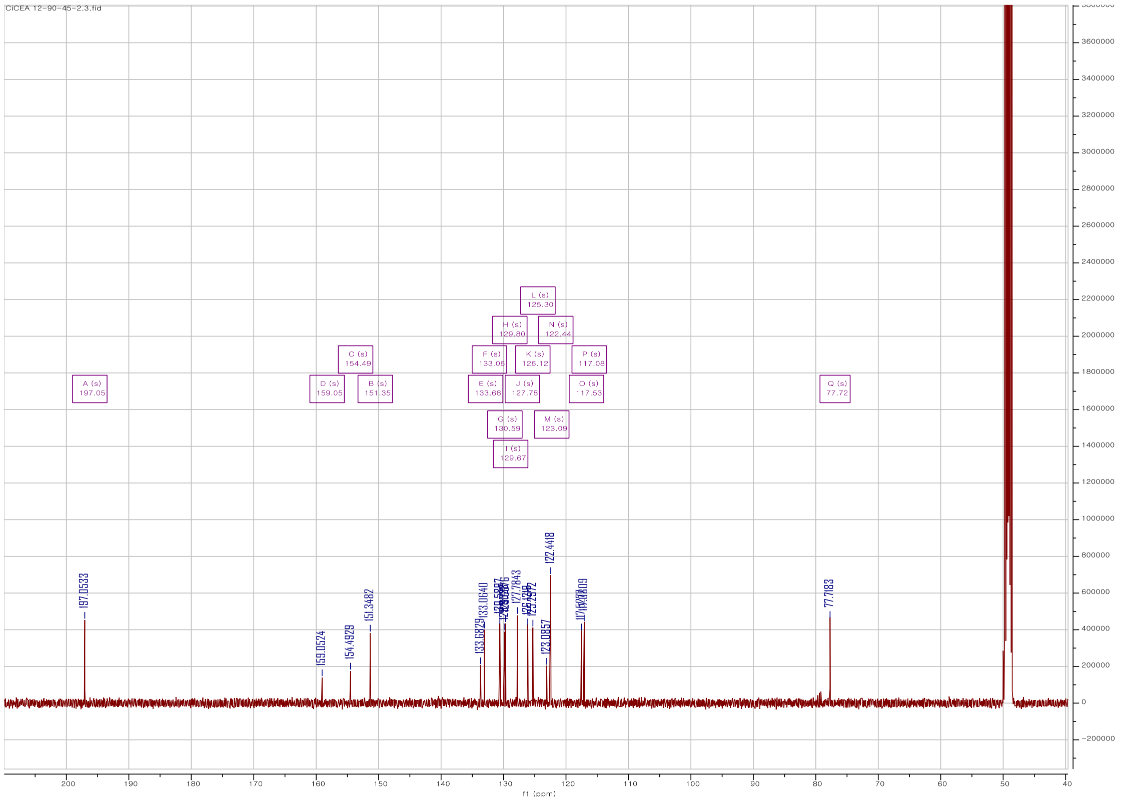

Fig. S.2.3. $^{13}$C-NMR spectrum of **20** in $CH_3OH$-$d_4$ at 125 MHz

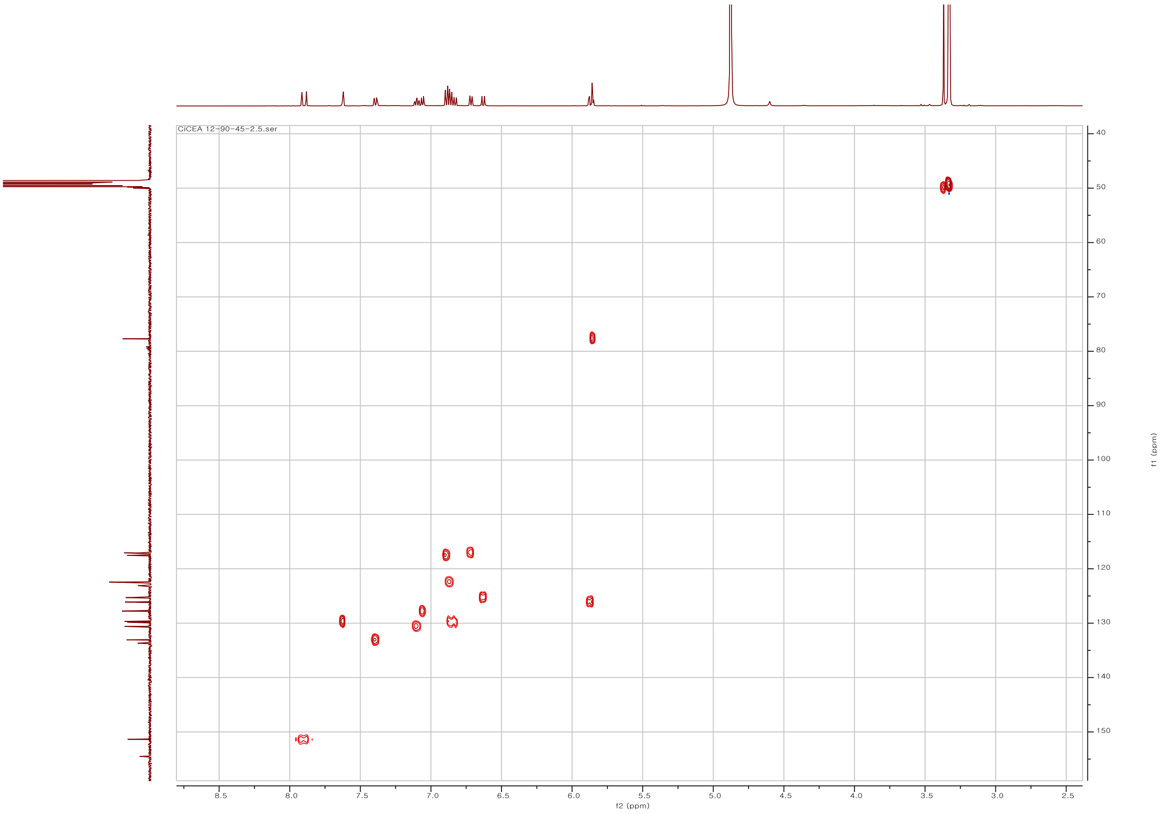

Fig. S.2.4. HSQC spectrum of **20** in $CH_3OH$-$d_4$

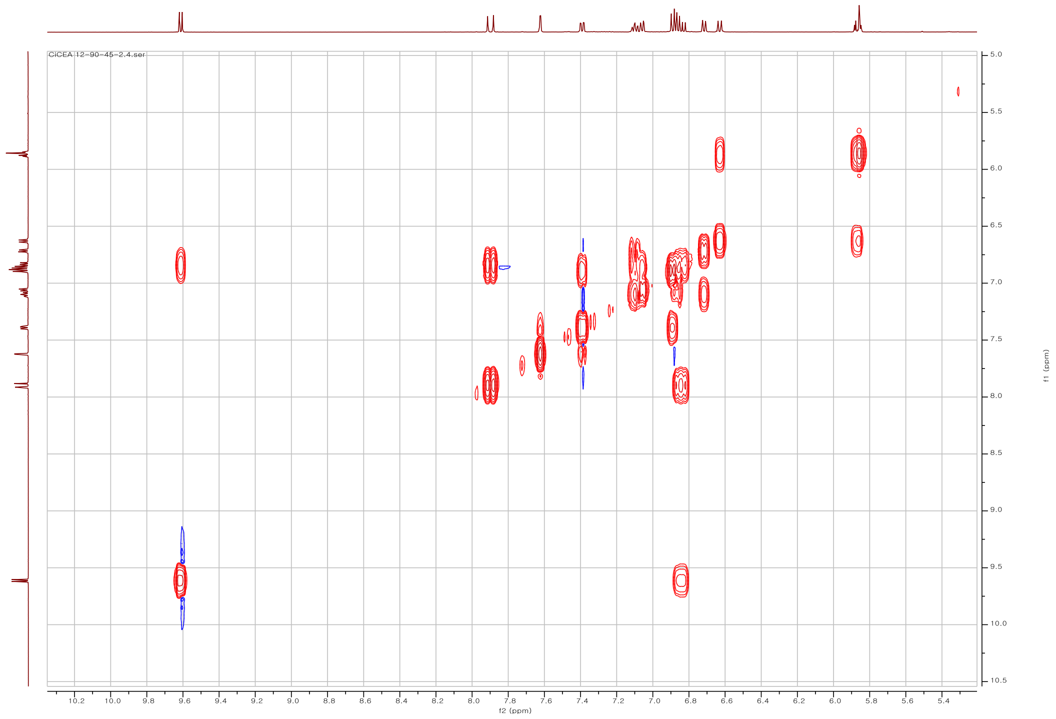

Fig. S.2.5. COSY spectrum of **20** in $CH_3OH$-$d_4$

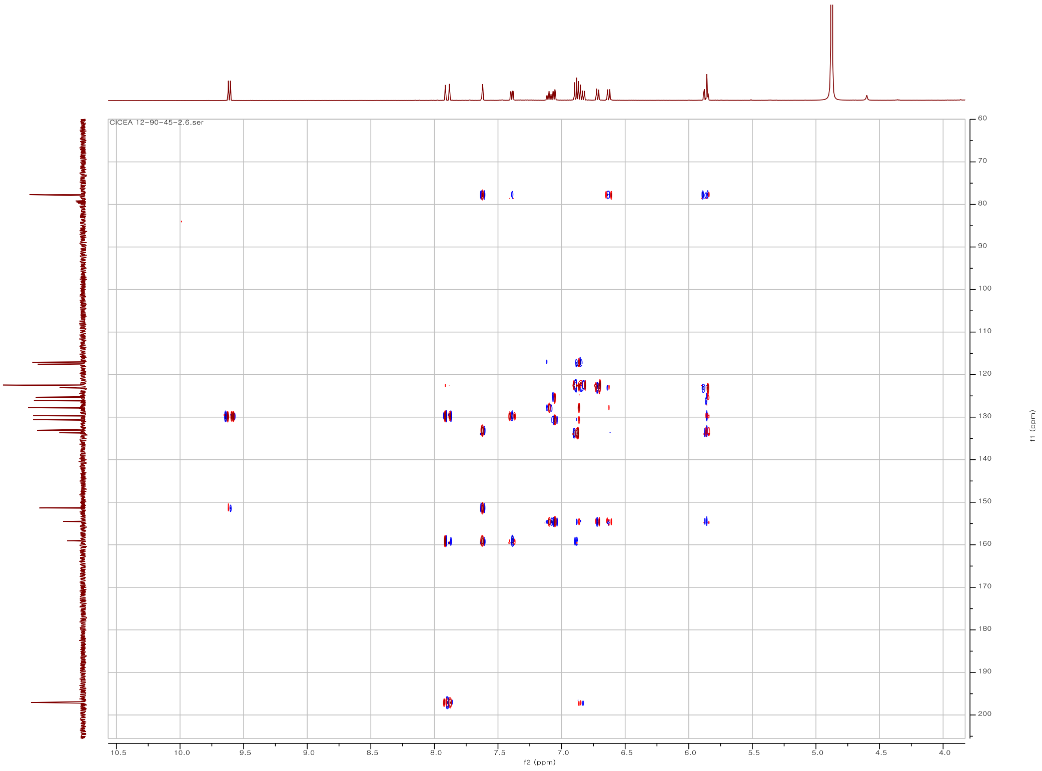

Fig. S.2.6. HMBC spectrum of **20** in $CH_3OH$-$d_4$

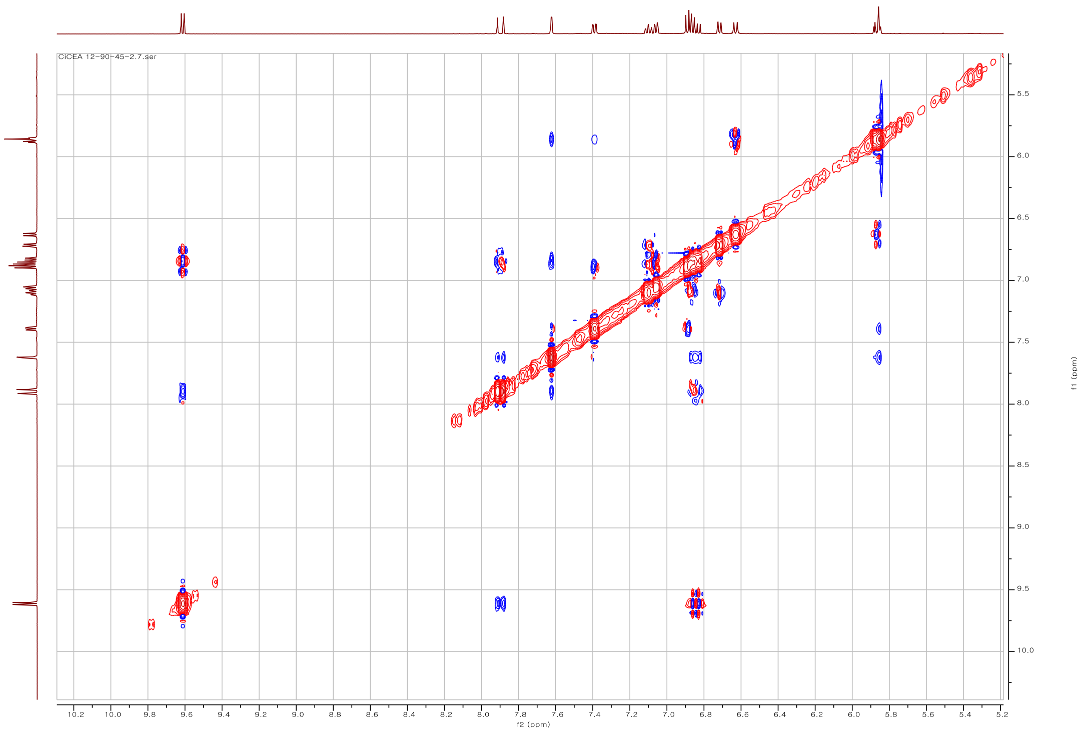

Fig. S.2.7. NOESY spectrum of **20** in $CH_3OH$-$d_4$

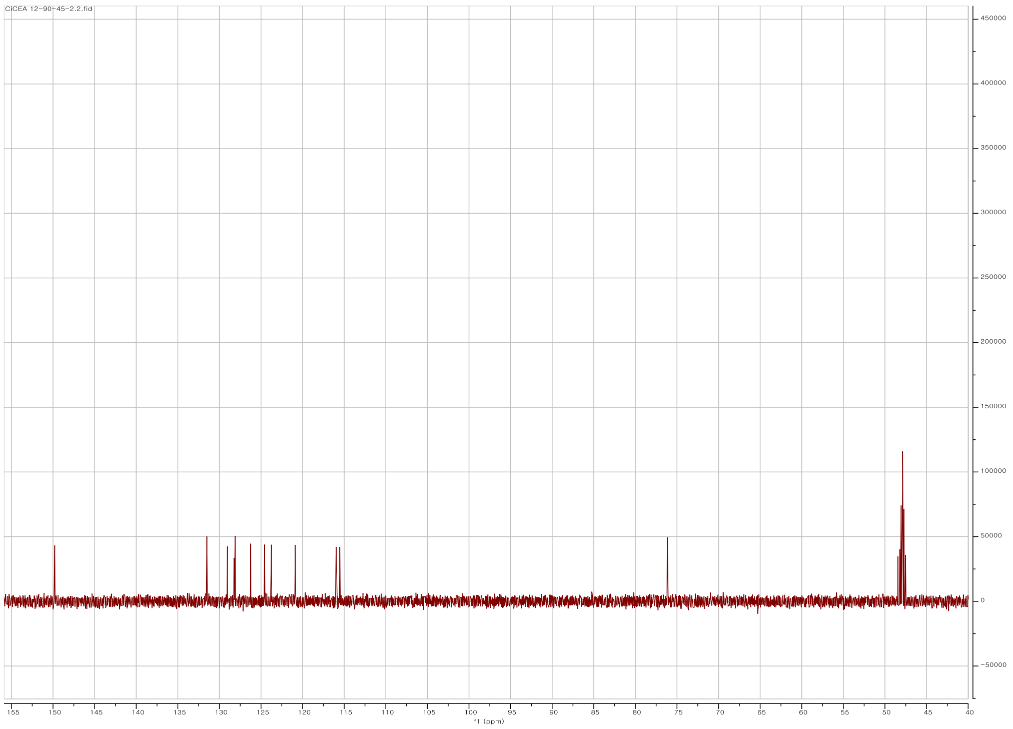

Fig. S.2.8. DEPT135 spectrum of **20** in $CH_3OH$-$d_4$

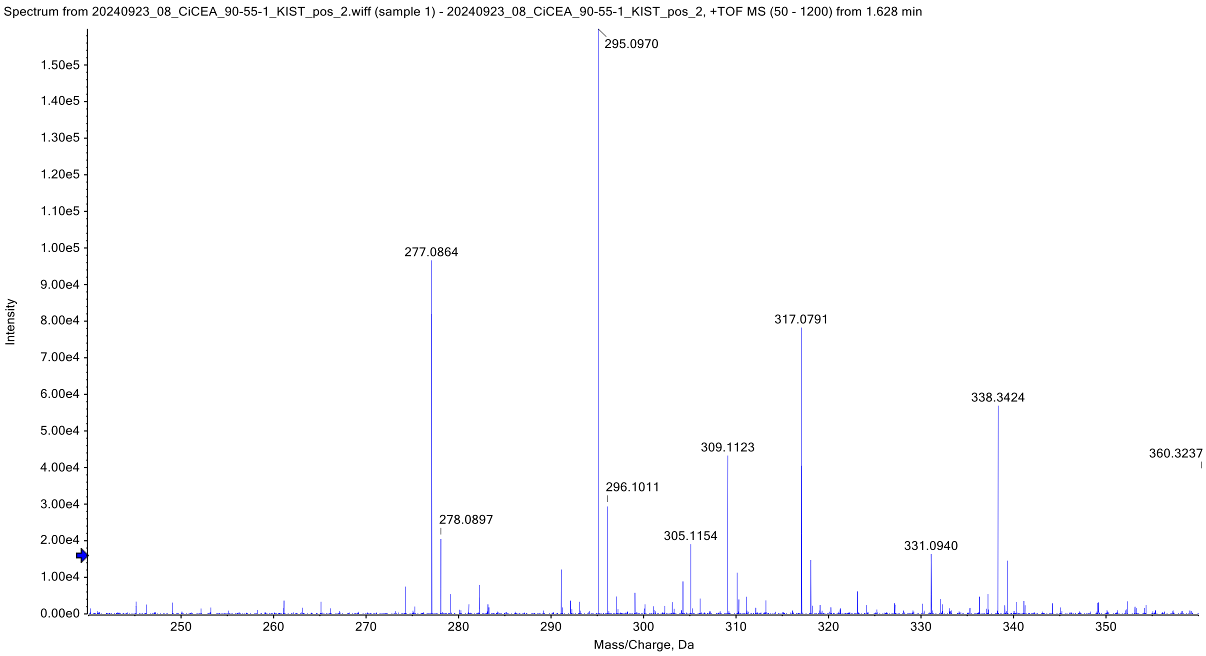

Fig. S.3.1. HR ESI-MS spectrum of **21**

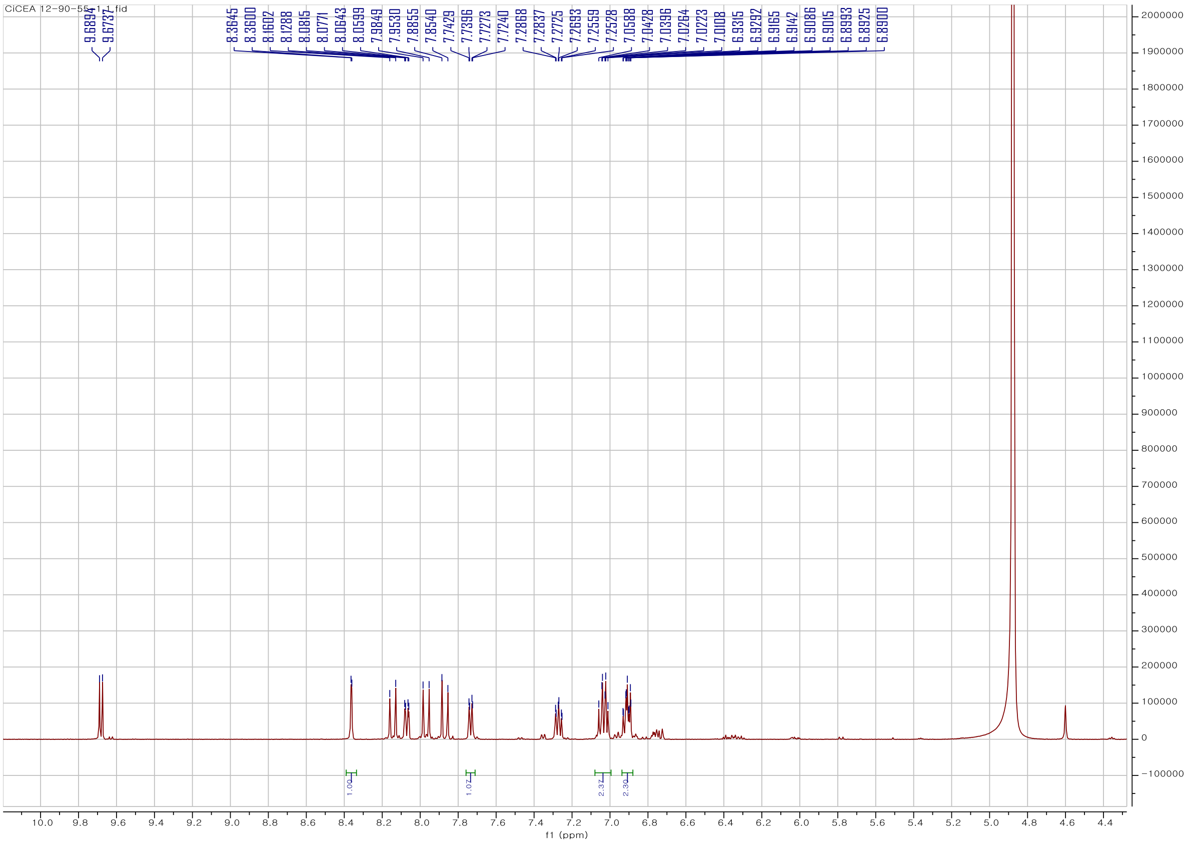

Fig. S.3.2. $^{1}$H-NMR spectrum of **21** in $CH_3OH$-$d_4$ at 500 MHz

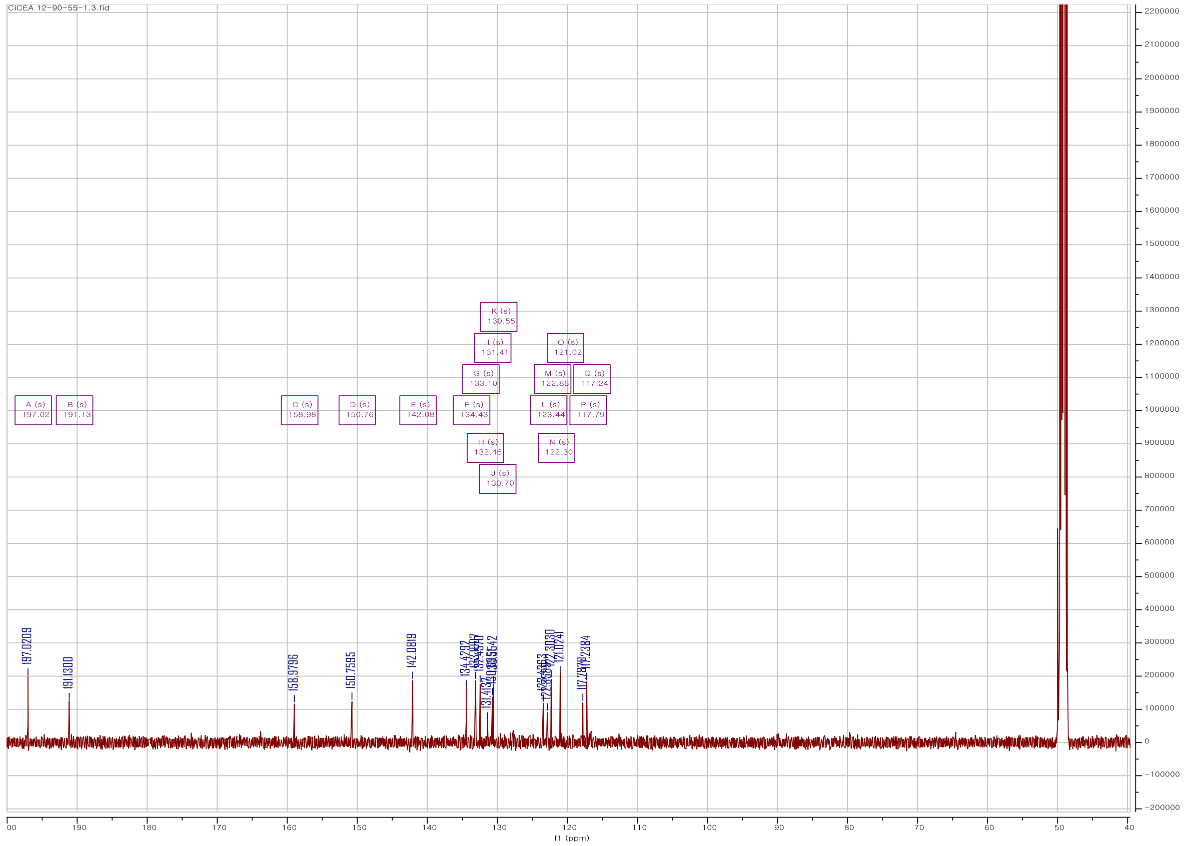

Fig. S.3.3. $^{13}$C-NMR spectrum of **21** in $CH_3OH$-$d_4$ at 125 MHz

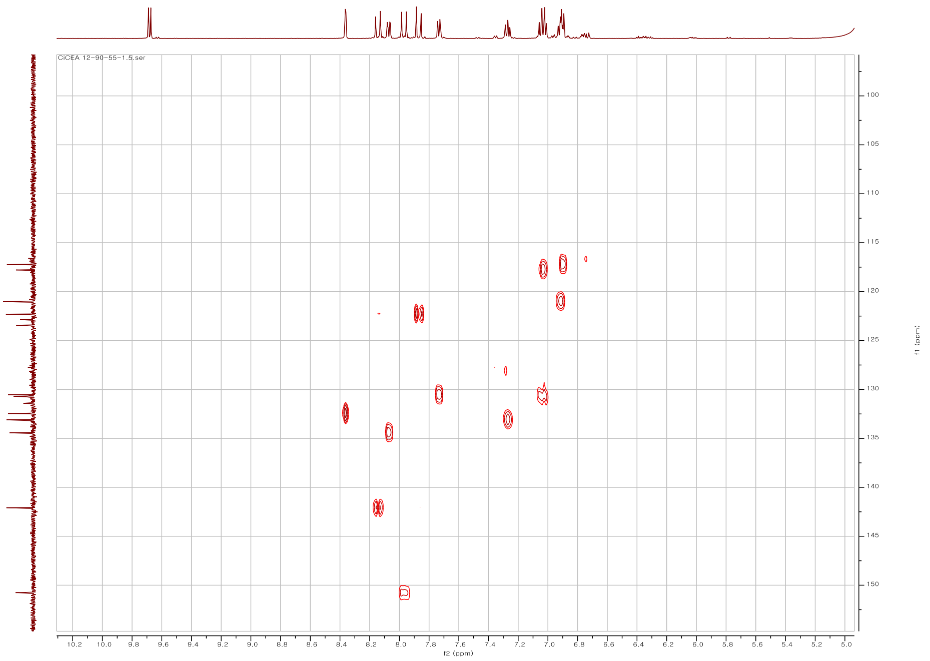

Fig. S.3.4. HSQC spectrum of **21** in $CH_3OH$-$d_4$

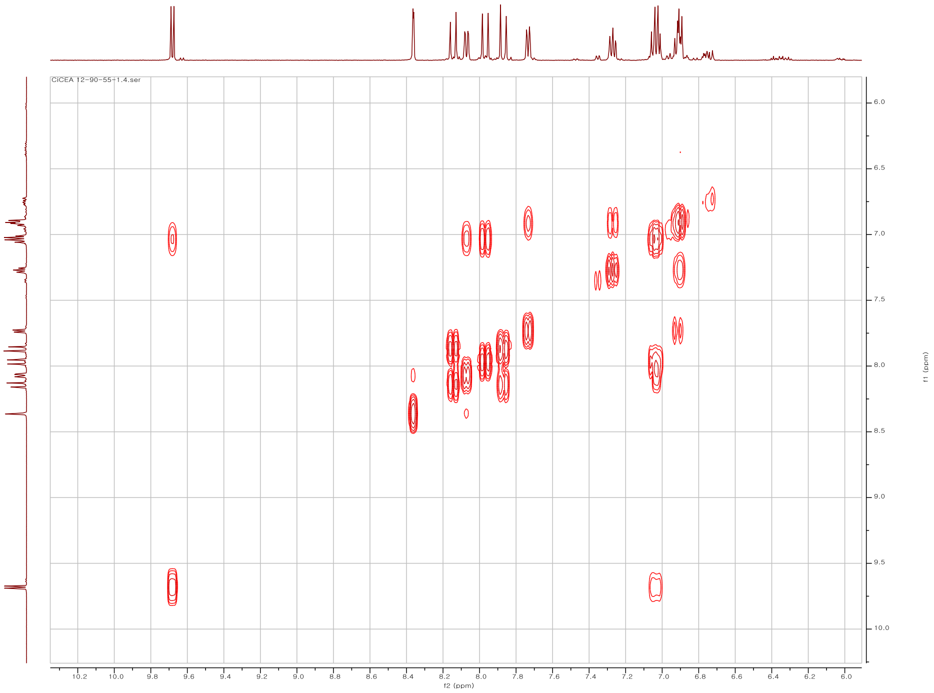

Fig. S.3.5. COSY spectrum of **21** in $CH_3OH$-$d_4$

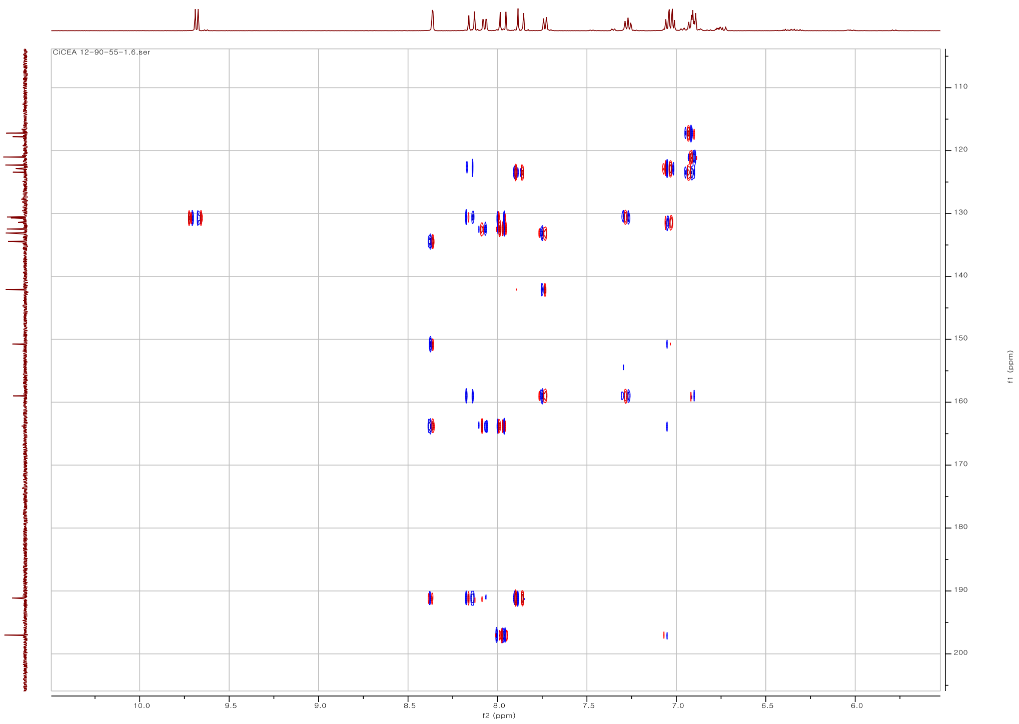

Fig. S.3.6. HMBC spectrum of **21** in $CH_3OH$-$d_4$

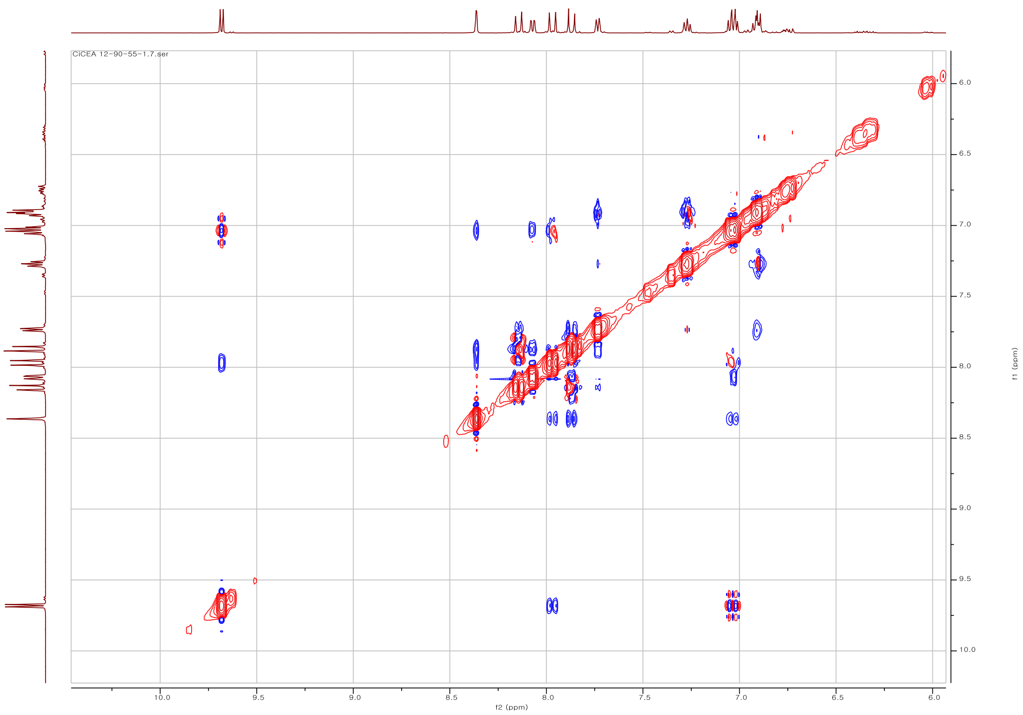

Fig. S.3.7. NOESY spectrum of **21** in $CH_3OH$-$d_4$

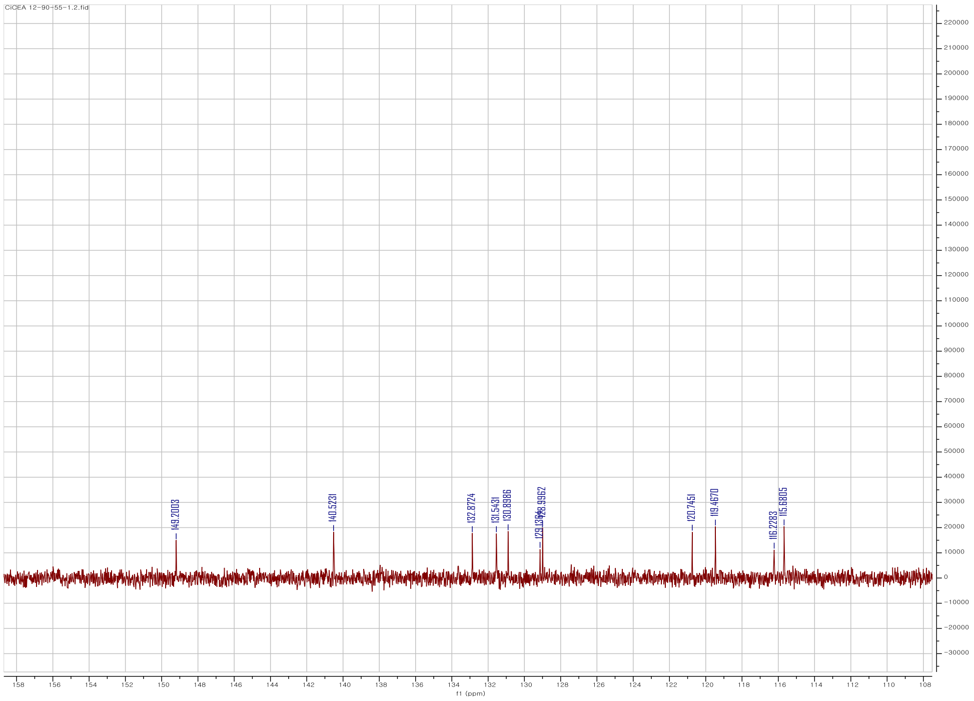

Fig. S.3.8. DEPT135 spectrum of **21** in $CH_3OH$-$d_4$

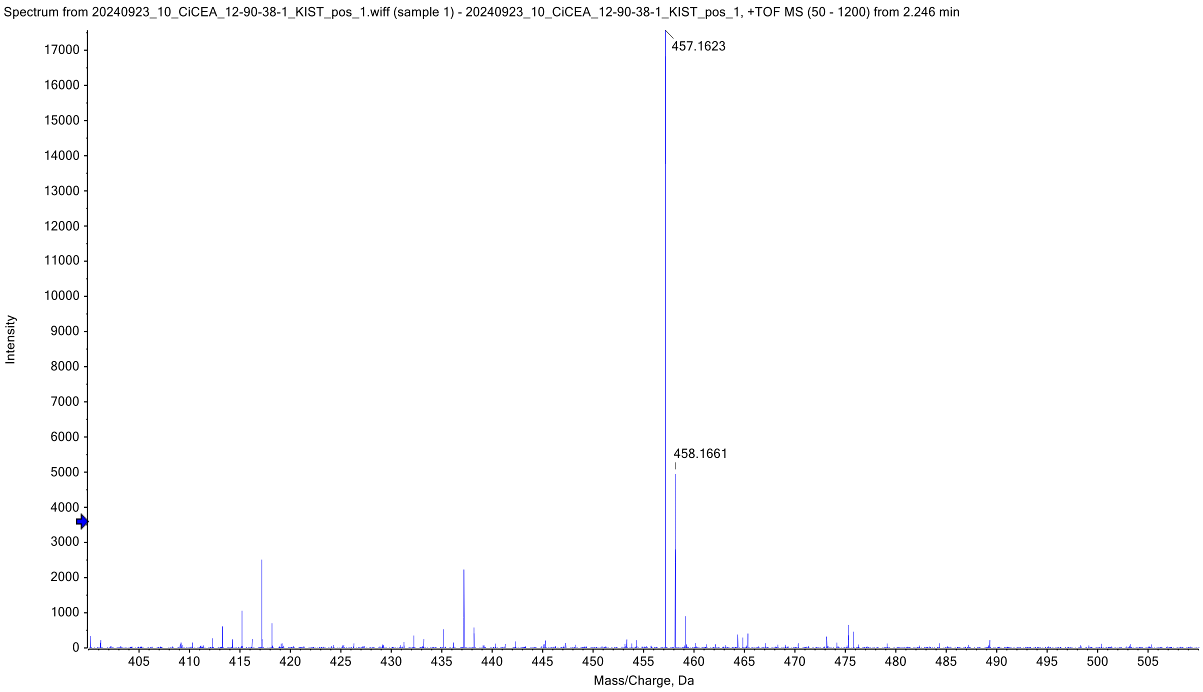

Fig. S.4.1. HR ESI-MS spectrum of **22**

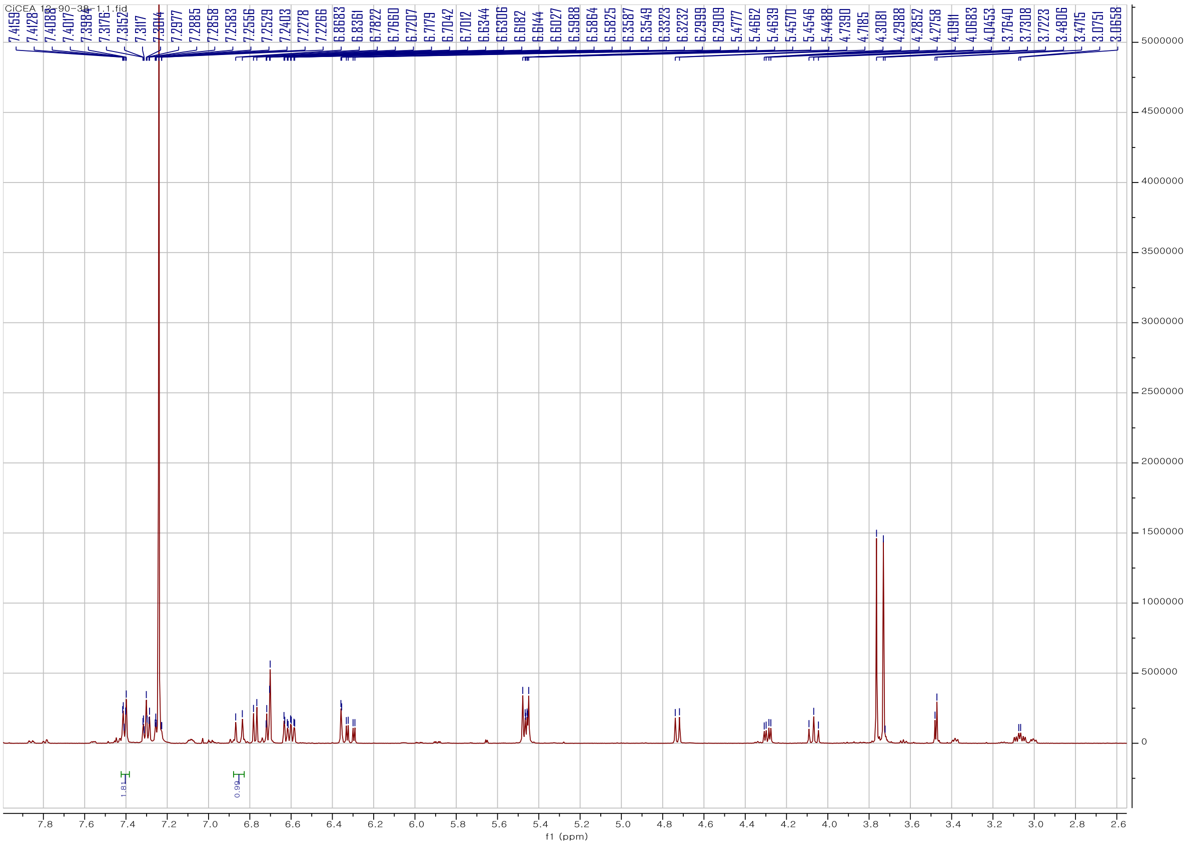

Fig. S.4.2. [1]H-NMR spectrum of **22** in $CHCl_3$-*d* at 500 MHz

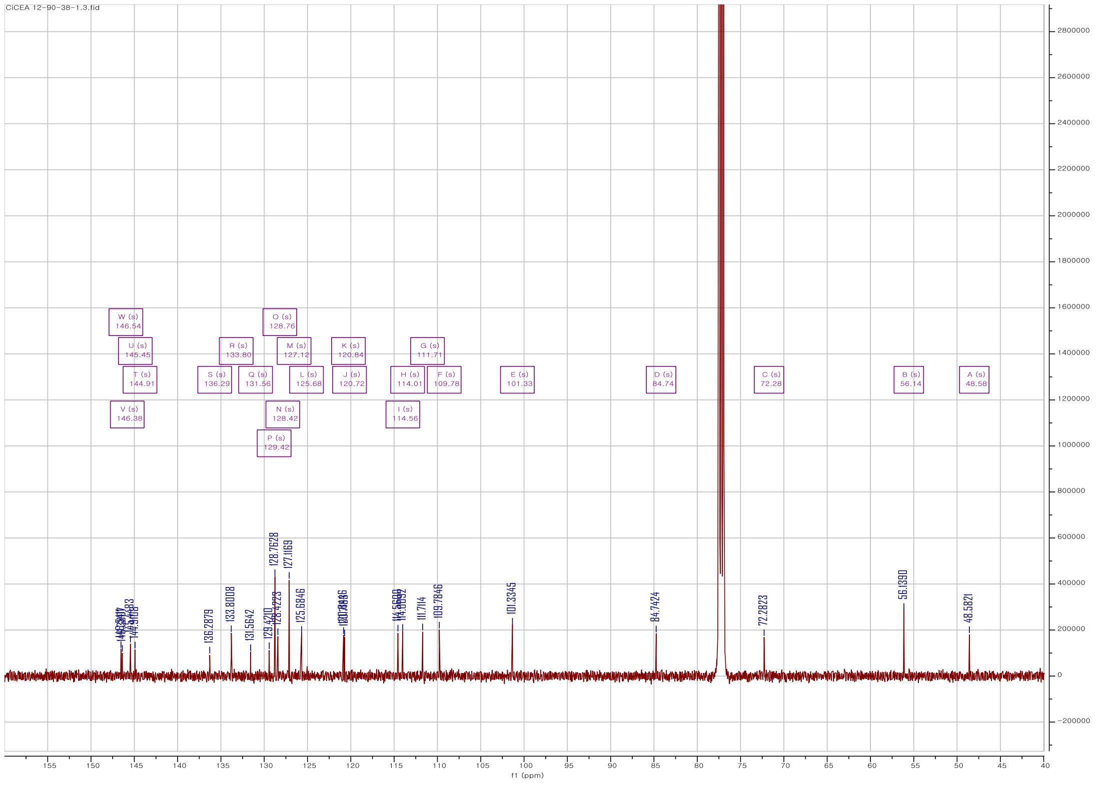

Fig. S.4.3. $^{13}$C-NMR spectrum of **22** in $CHCl_3$-*d* at 125 MHz

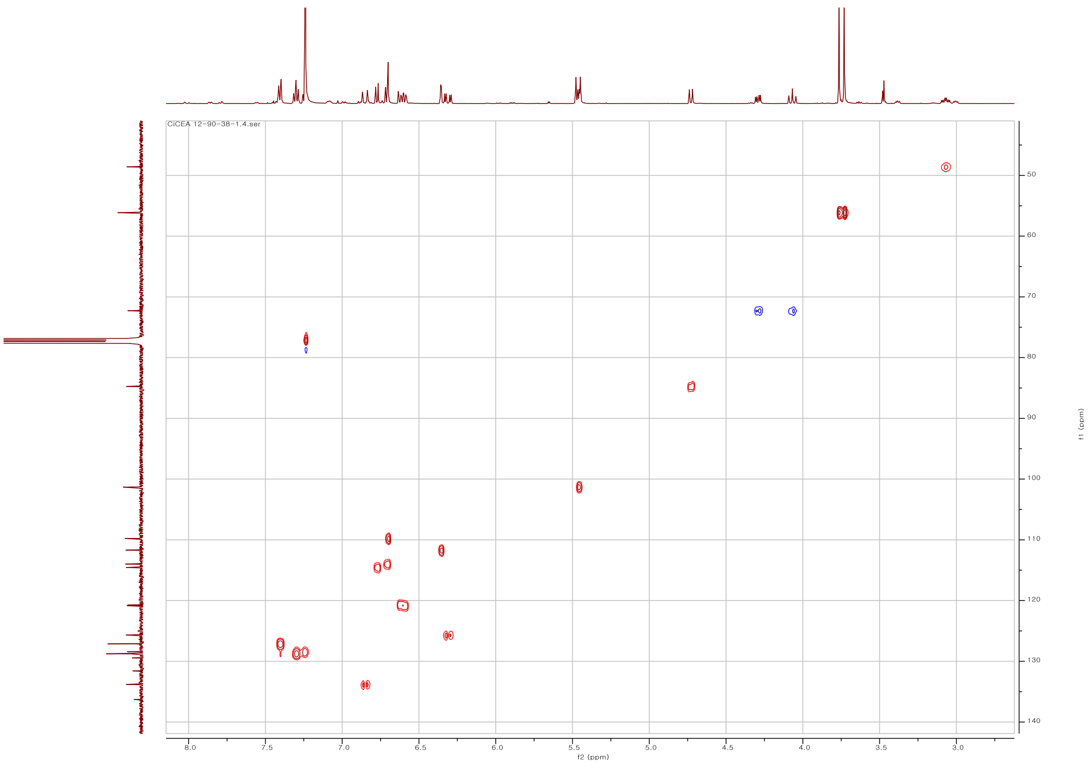

Fig. S.4.4. HSQC spectrum of **22** in $CHCl_3$-*d*

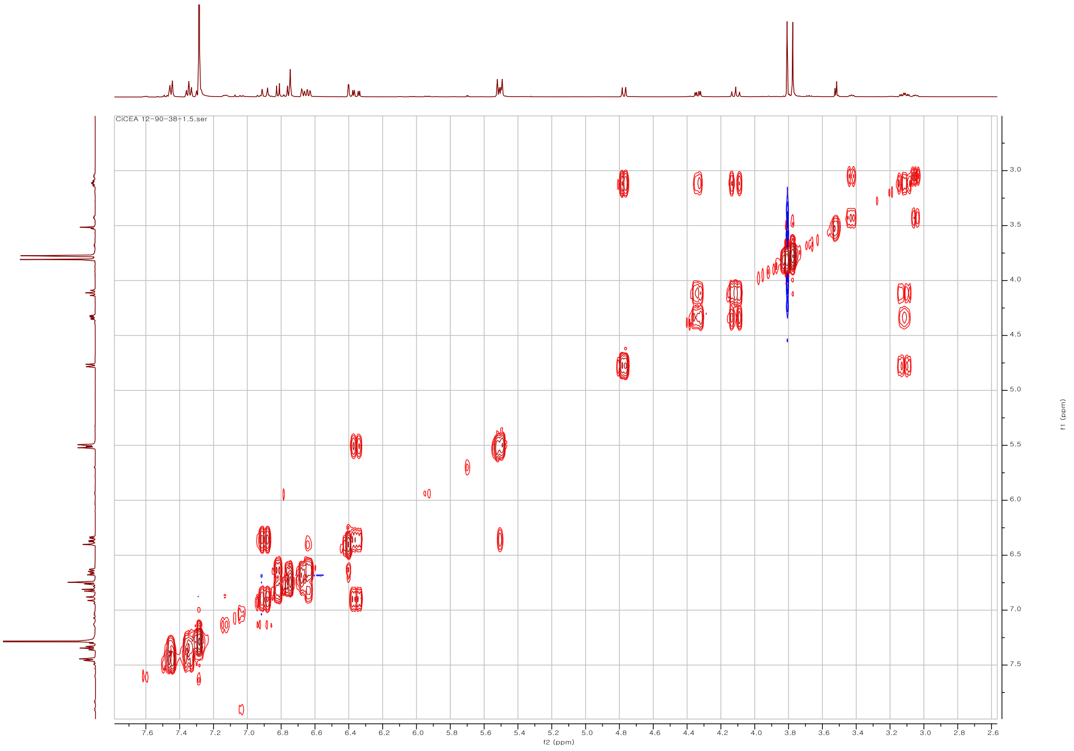

Fig. S.4.5. COSY spectrum of **22** in $CHCl_3$-*d*

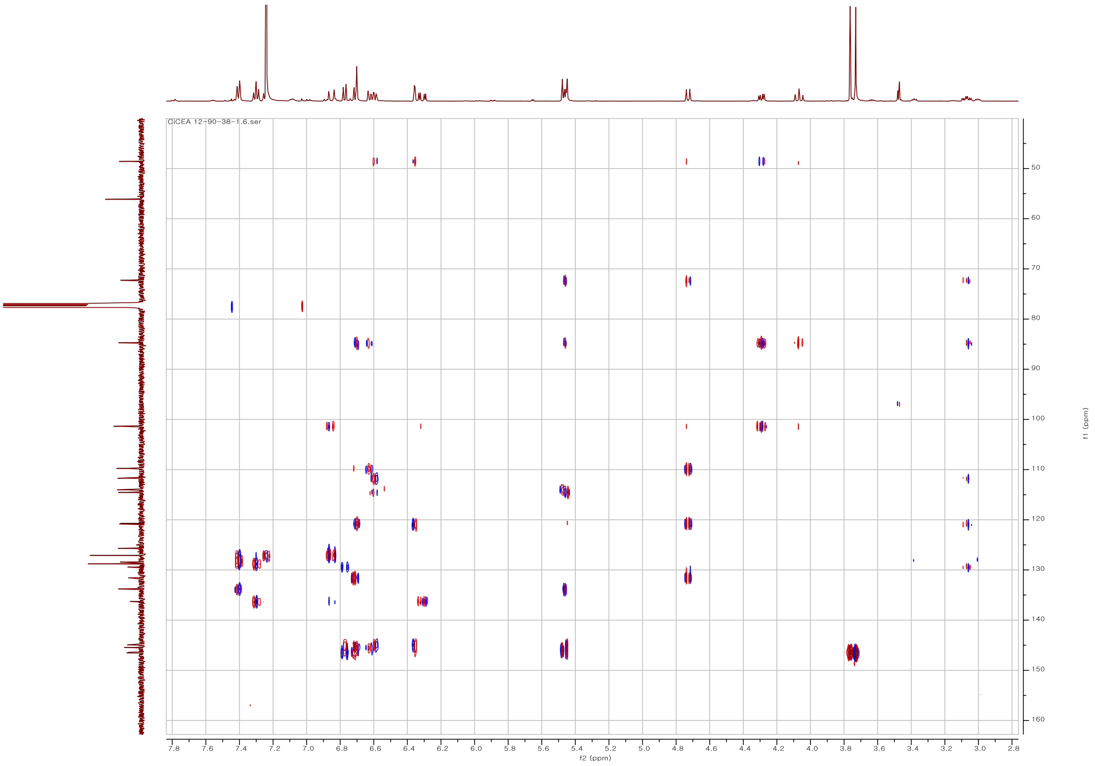

Fig. S.4.6. HMBC spectrum of **22** in $CHCl_3$-*d*

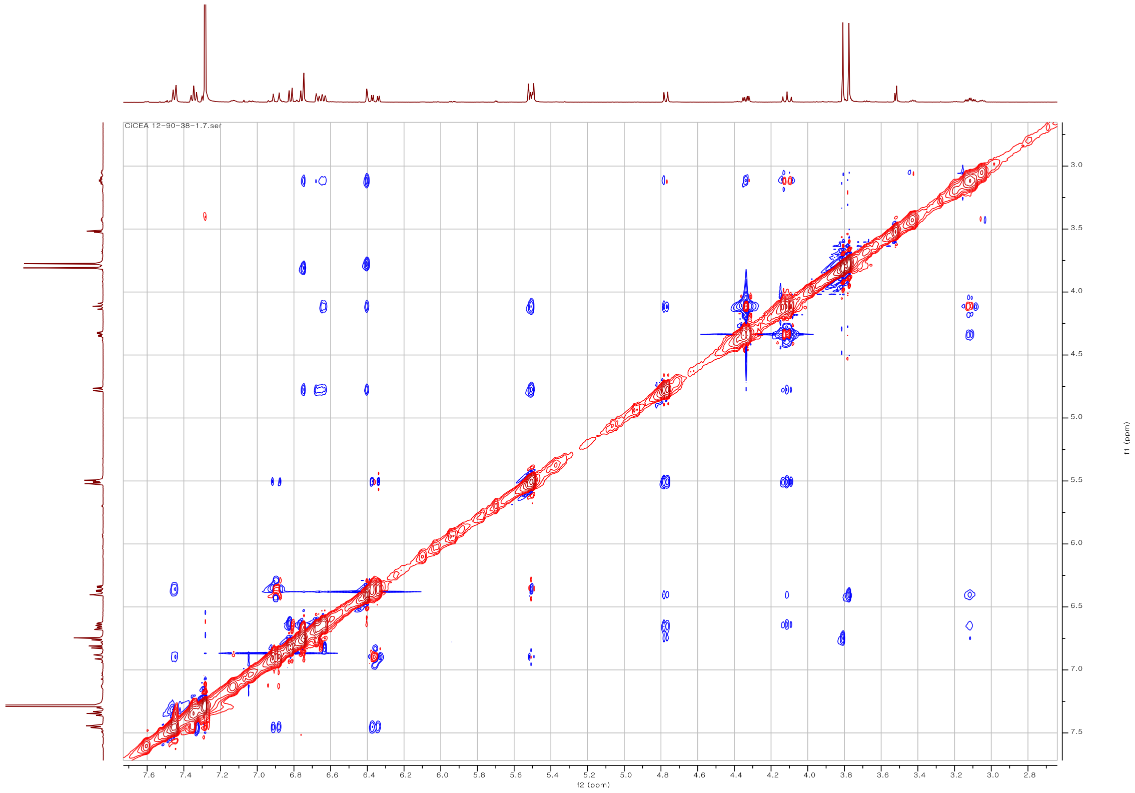

Fig. S.4.7. NOESY spectrum of **22** in $CHCl_3$-*d*

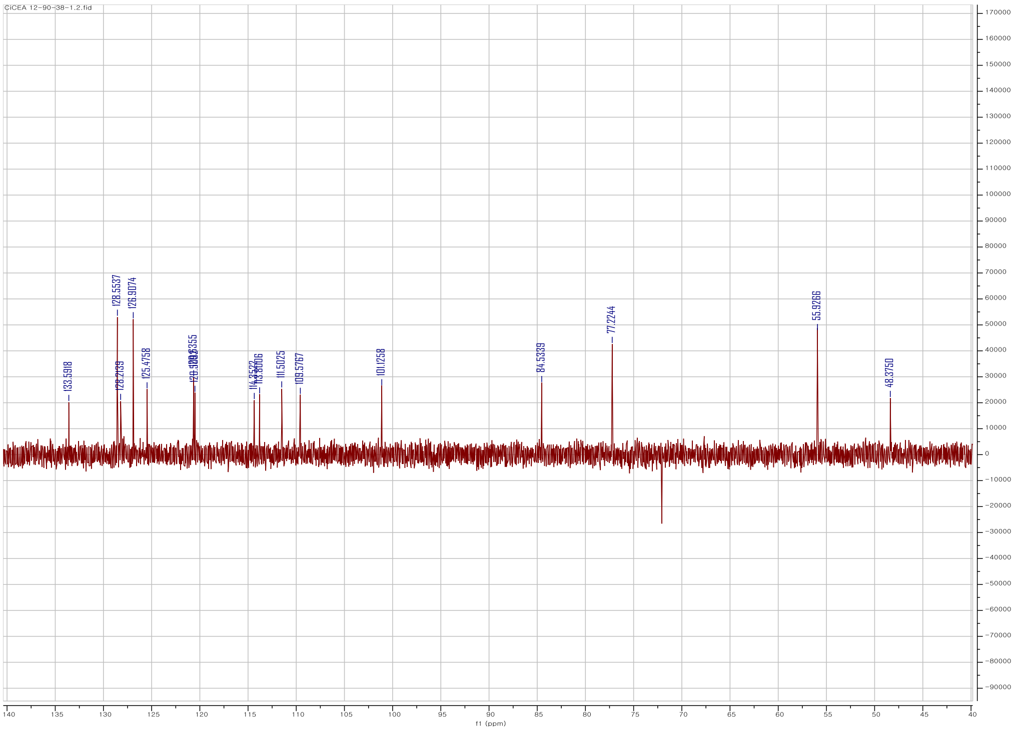

Fig. S.4.8. DEPT135 spectrum of **22** in $CHCl_3$-*d*

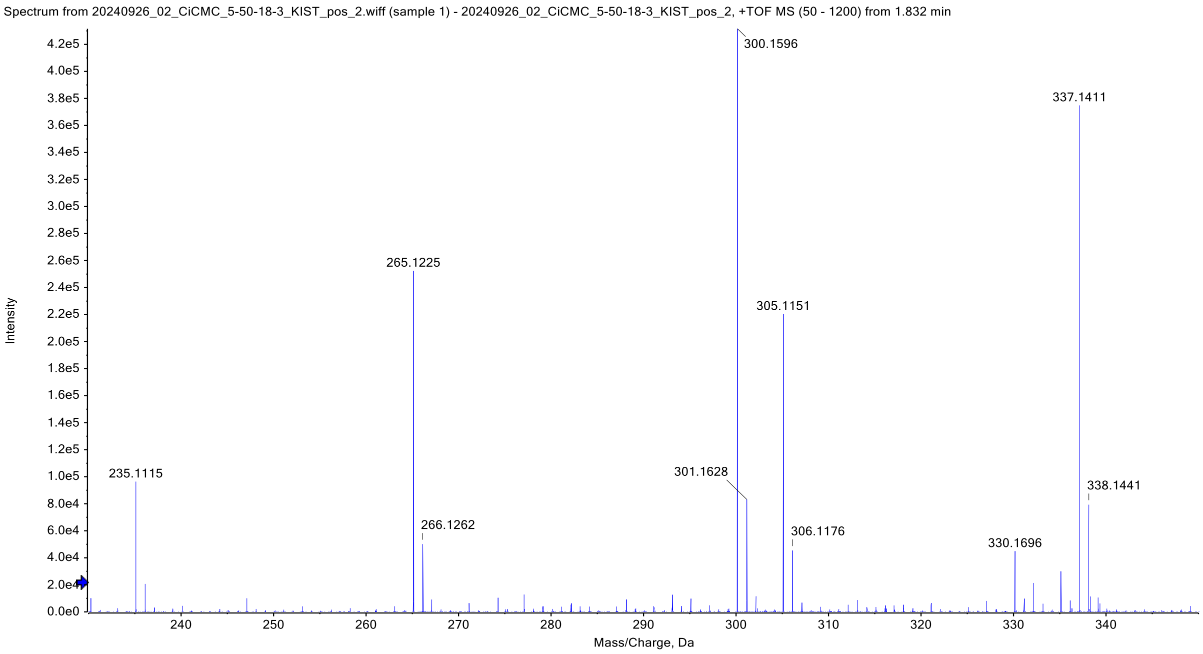

Fig. S.5.1. HR ESI-MS spectrum of **23**

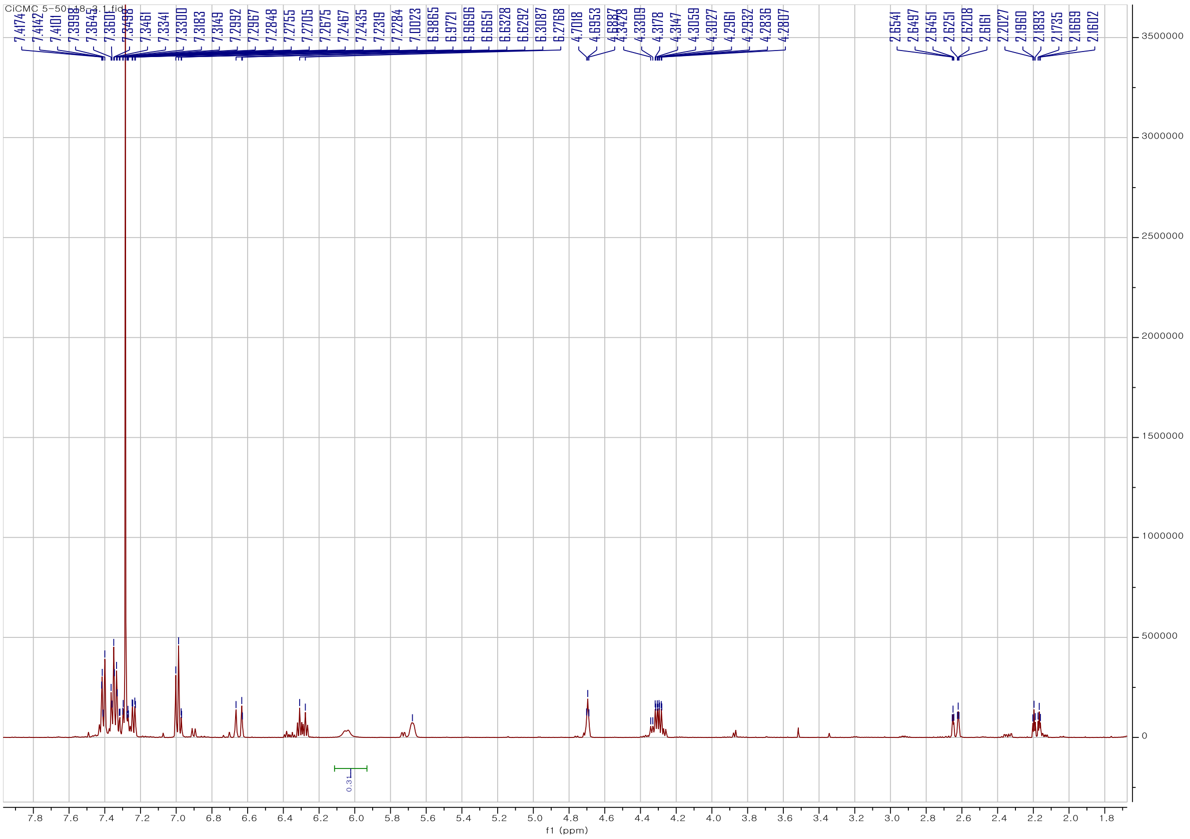

Fig. S.5.2. [1]H-NMR spectrum of **23** in $CHCl_3$-*d* at 500 MHz

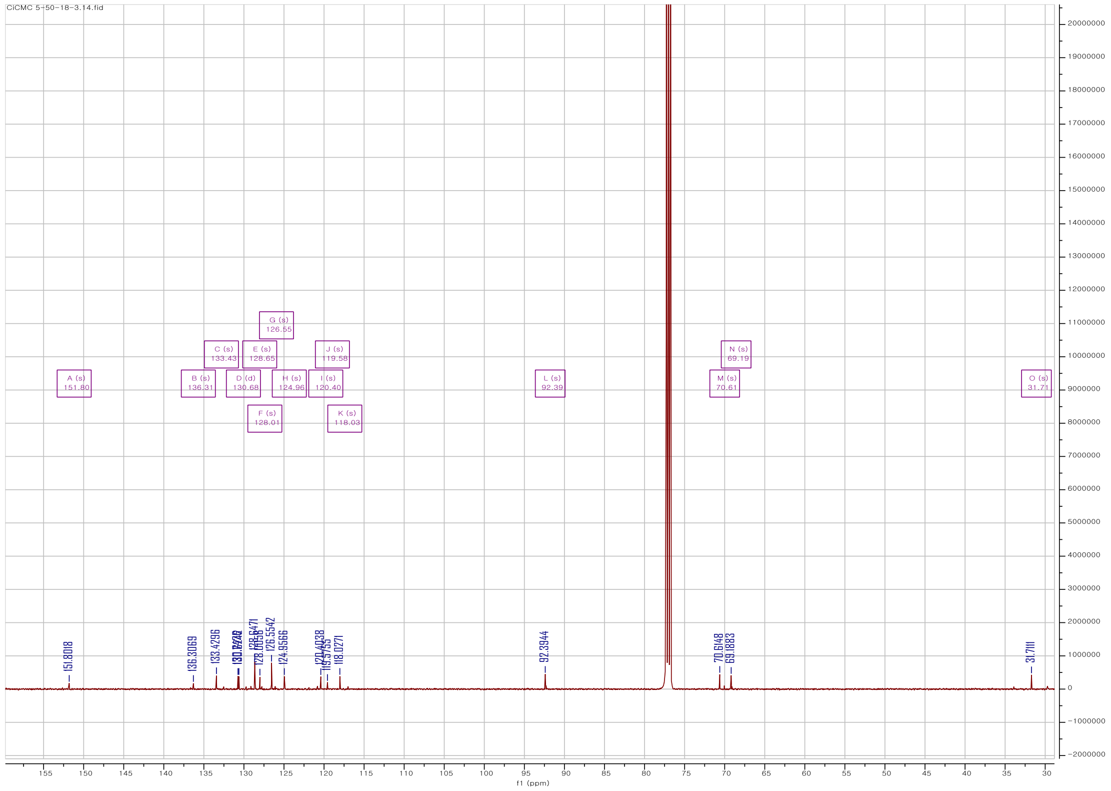

Fig. S.5.3. $^{13}$C-NMR spectrum of **23** in $CHCl_3$-*d* at 125 MHz

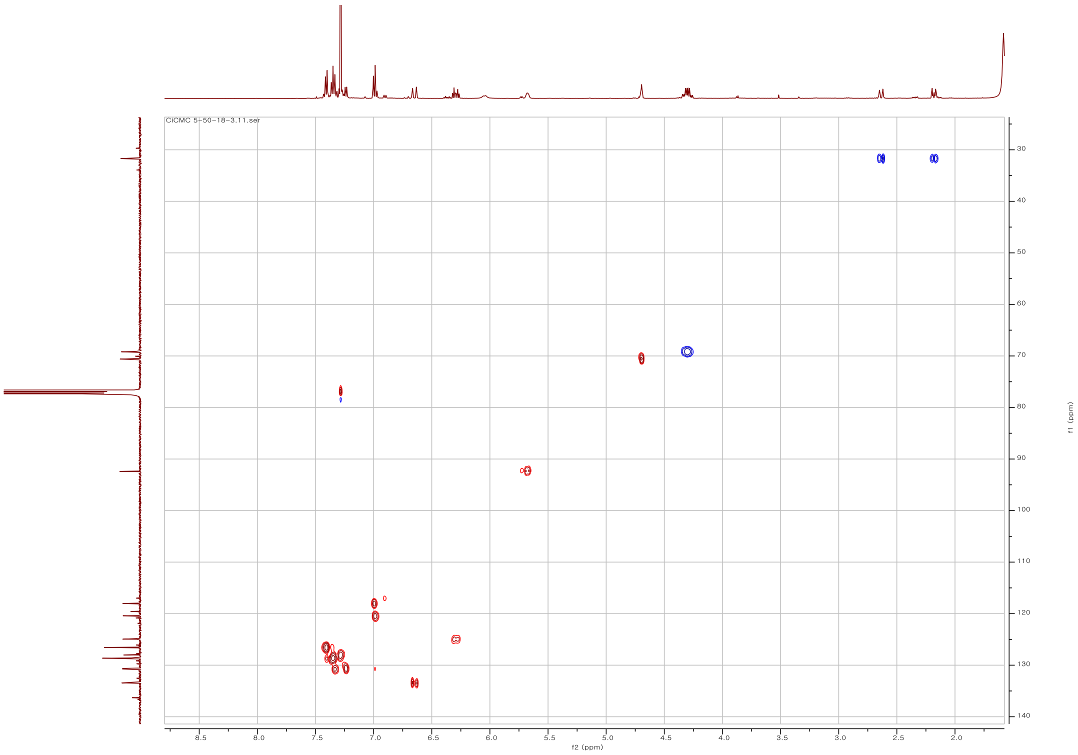

Fig. S.5.4. HSQC spectrum of **23** in $CHCl_3$-*d*

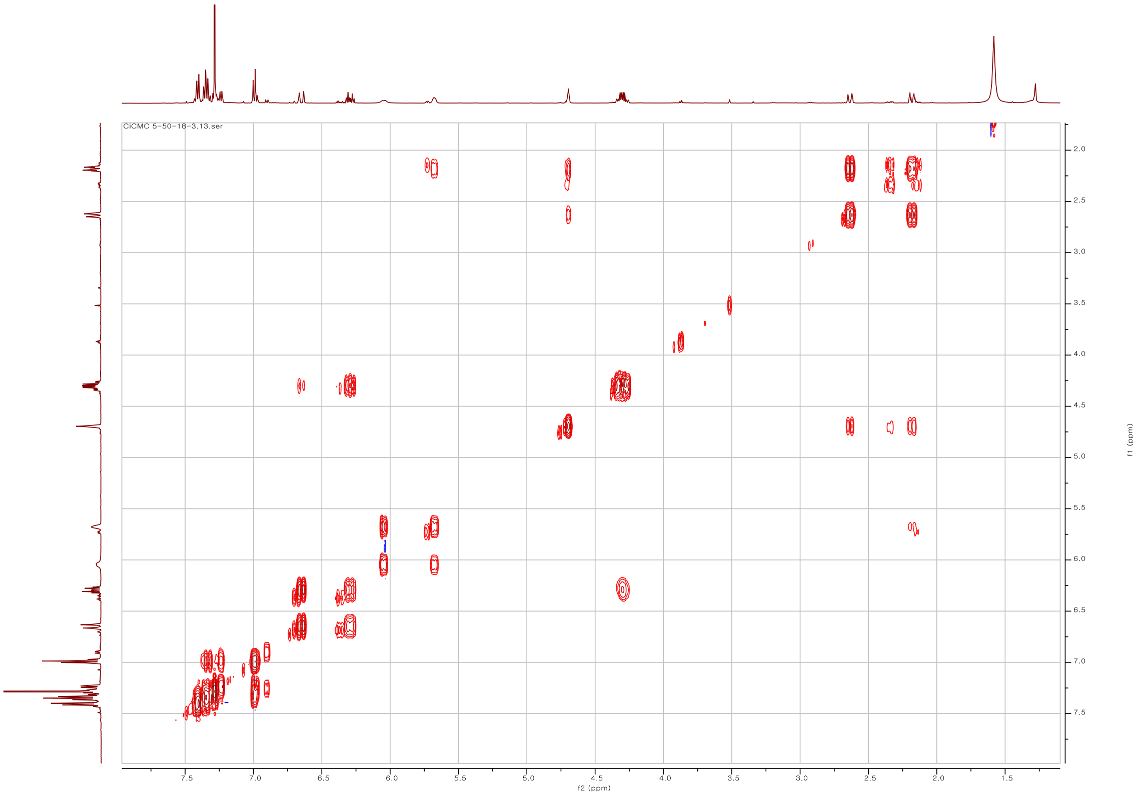

Fig. S.5.5. COSY spectrum of **23** in $CHCl_3$-*d*

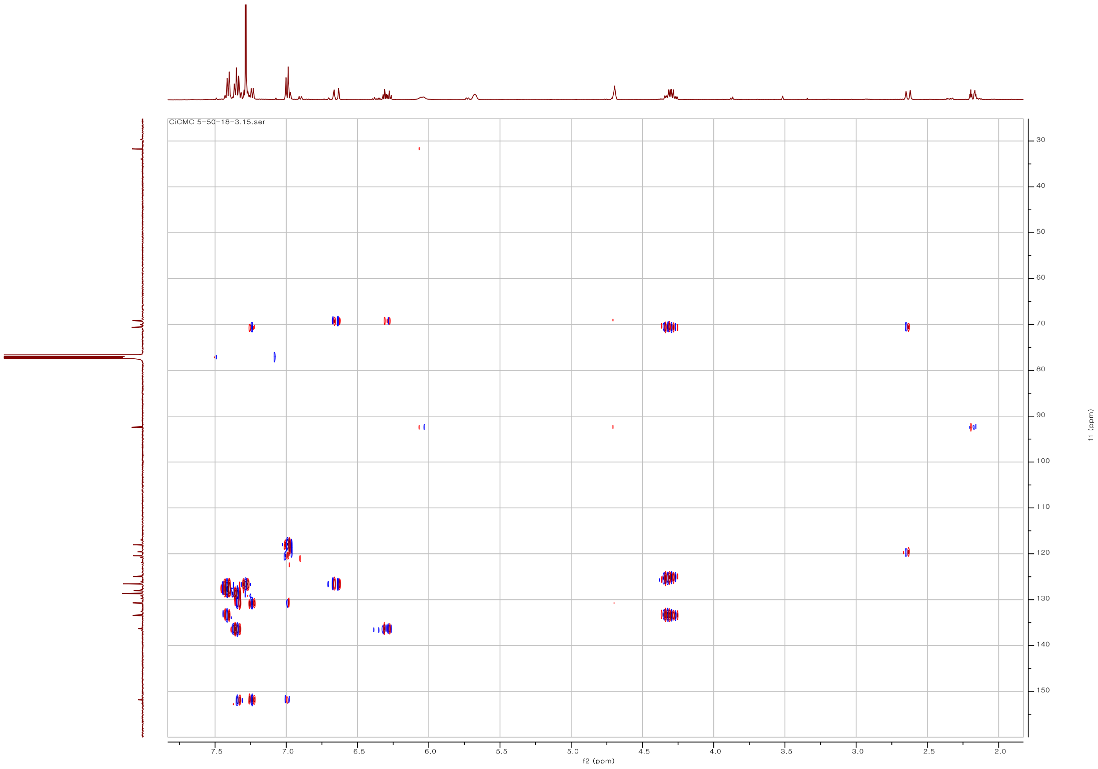

Fig. S.5.6. HMBC spectrum of **23** in $CHCl_3$-*d*

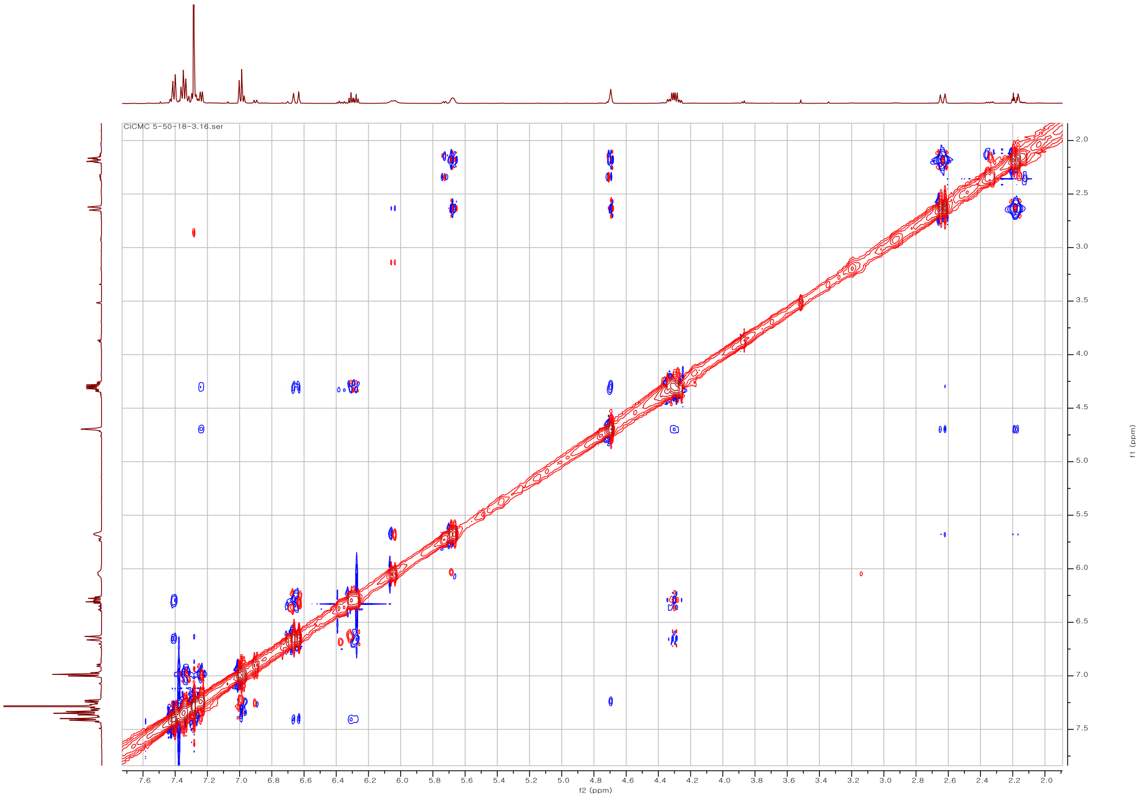

Fig. S.5.7. NOESY spectrum of **23** in $CHCl_3$-*d*

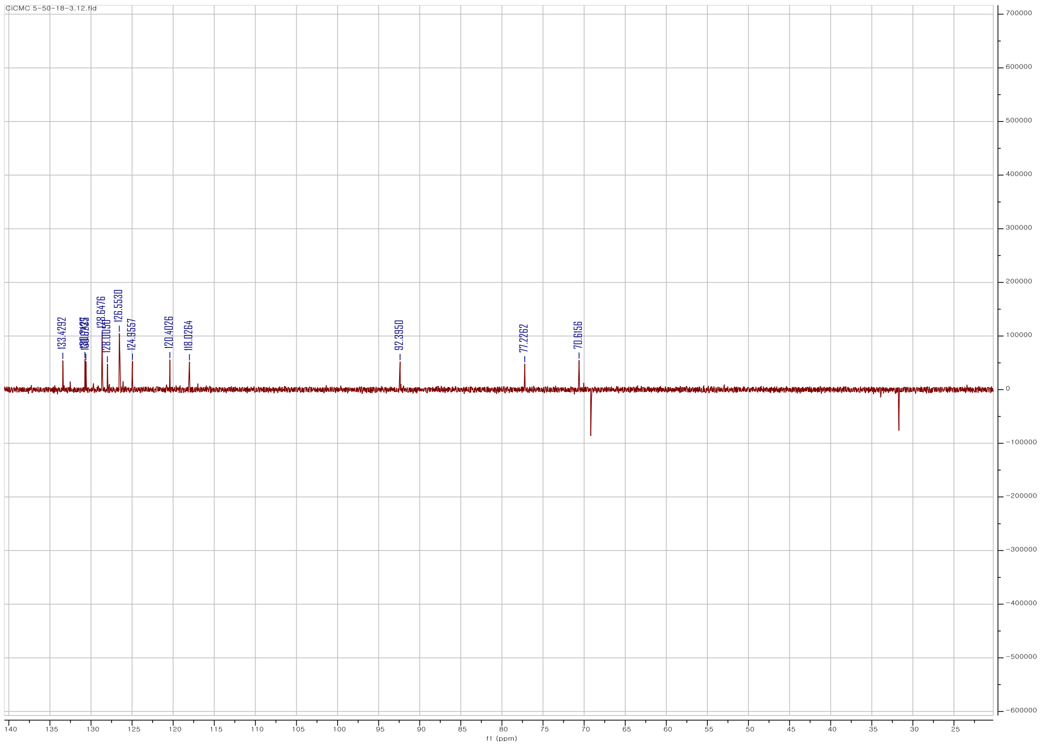

Fig. S.5.8. DEPT135 spectrum of **23** in $CHCl_3$-*d*

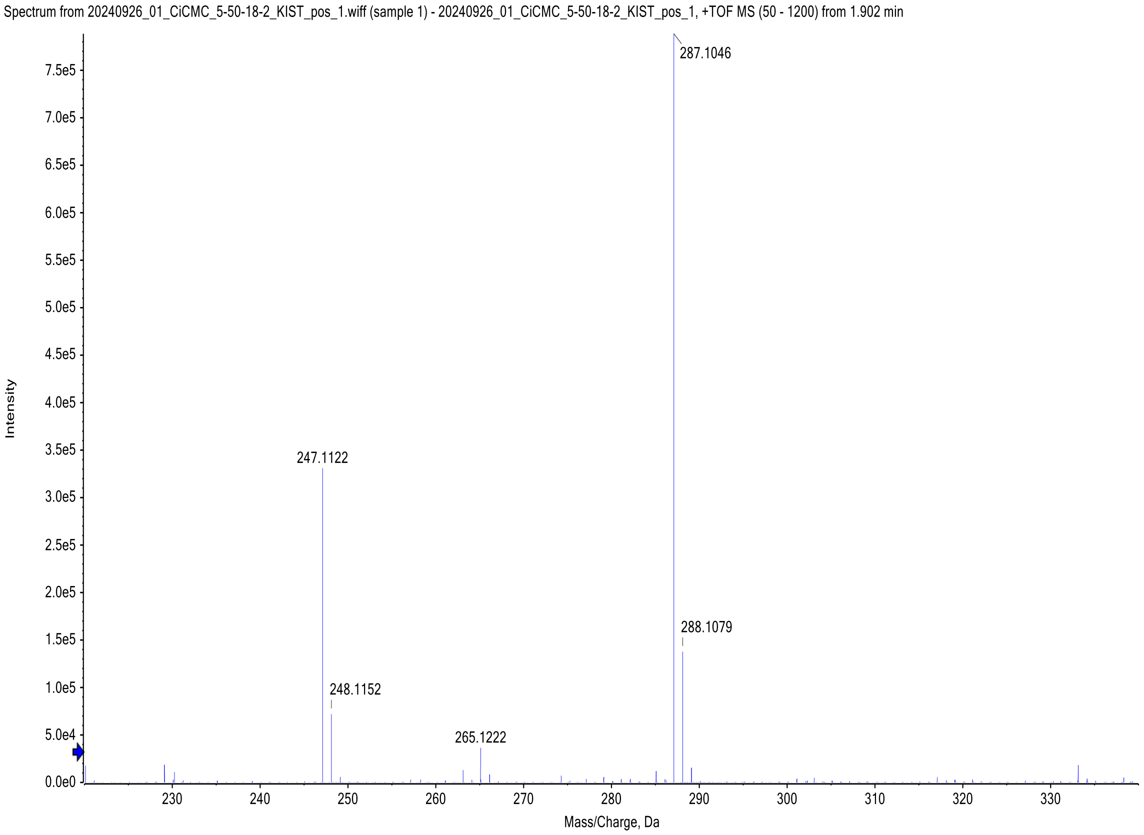

Fig. S.6.1. HR ESI-MS spectrum of **24**

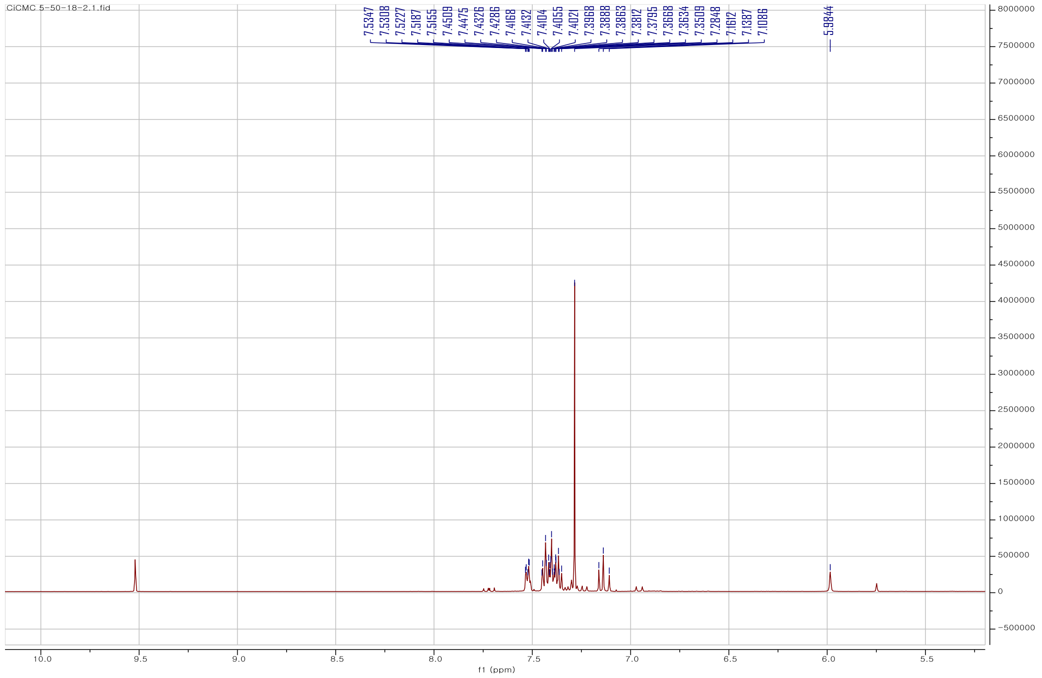

Fig. S.6.2. $^{1}$H-NMR spectrum of **24** in $CHCl_3$-*d* at 500 MHz

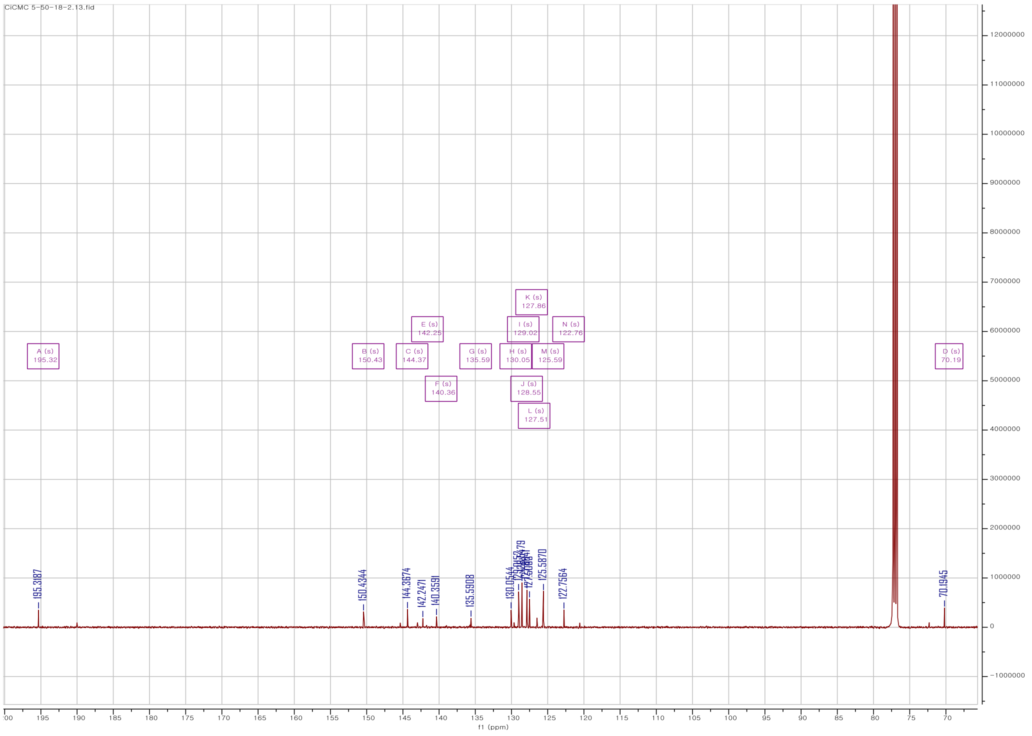

Fig. S.6.3. [13]C-NMR spectrum of **24** in $CHCl_3$-*d* at 125 MHz

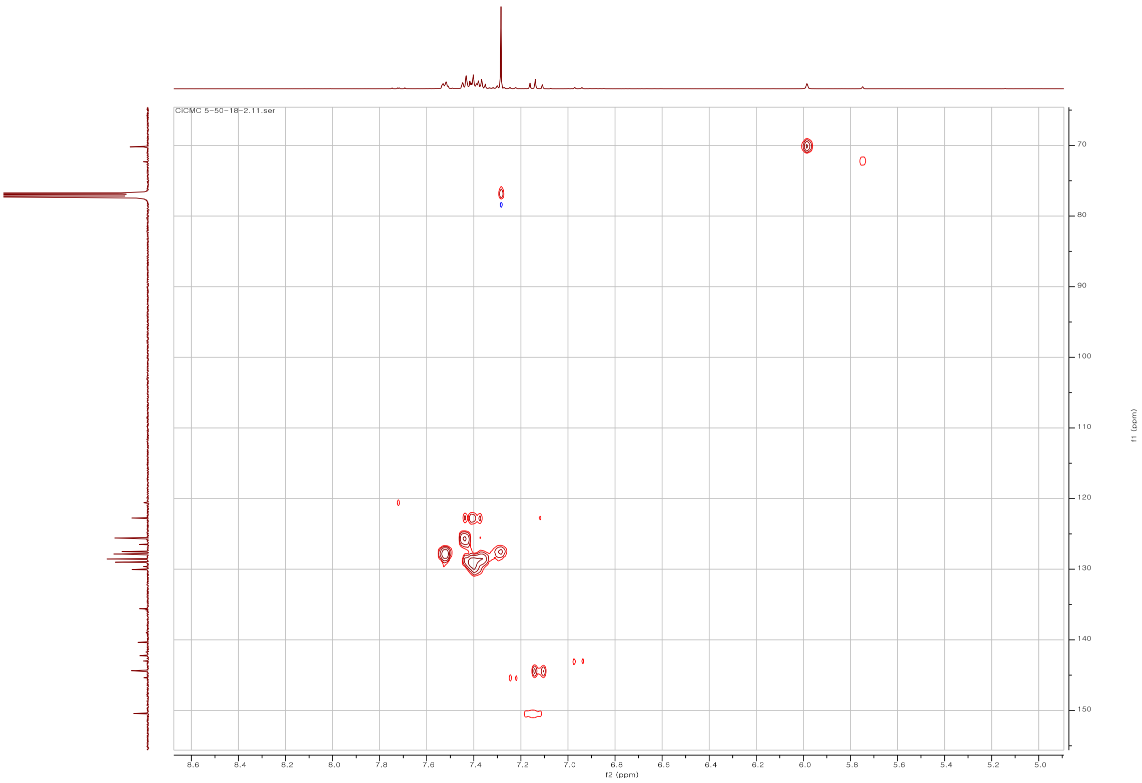

Fig. S.6.4. HSQC spectrum of **24** in $CHCl_3$-*d*

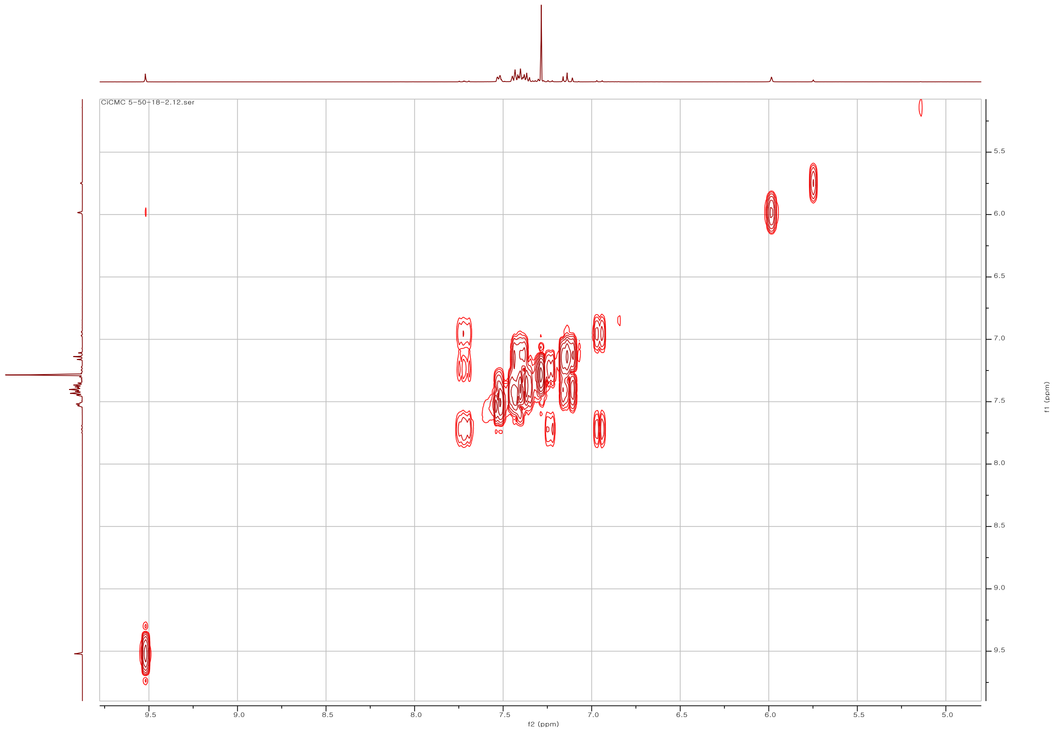

Fig. S.6.5. COSY spectrum of **24** in $CHCl_3$-*d*

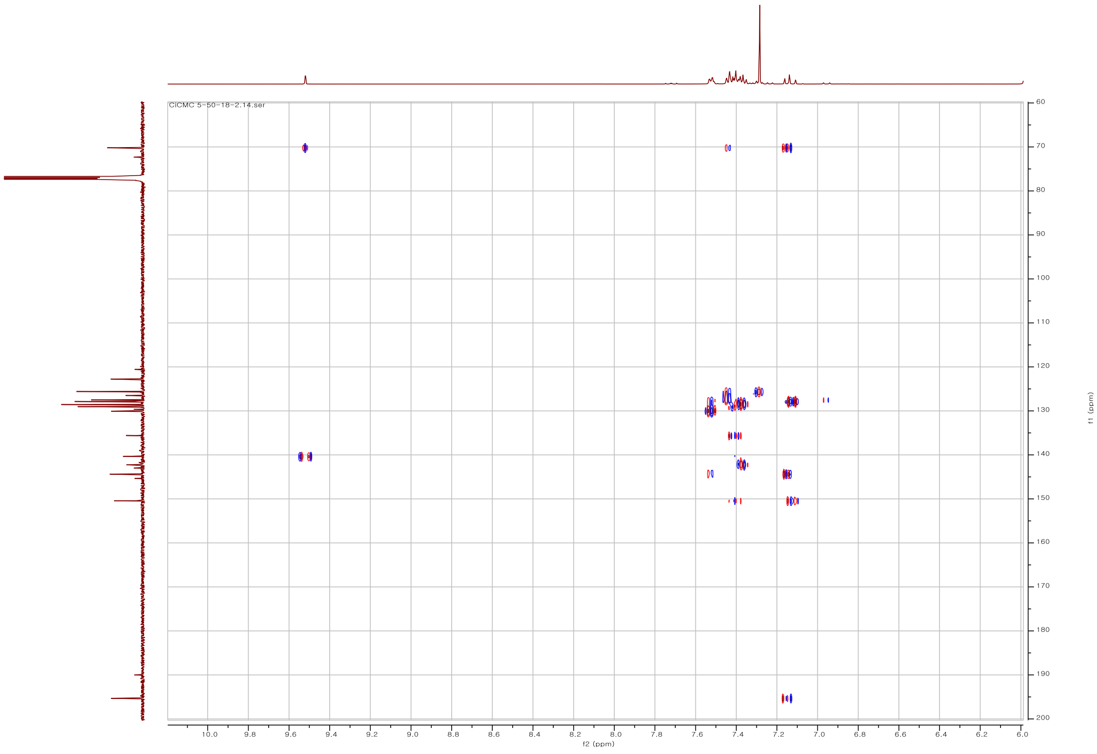

Fig. S.6.6. HMBC spectrum of **24** in $CHCl_3$-*d*

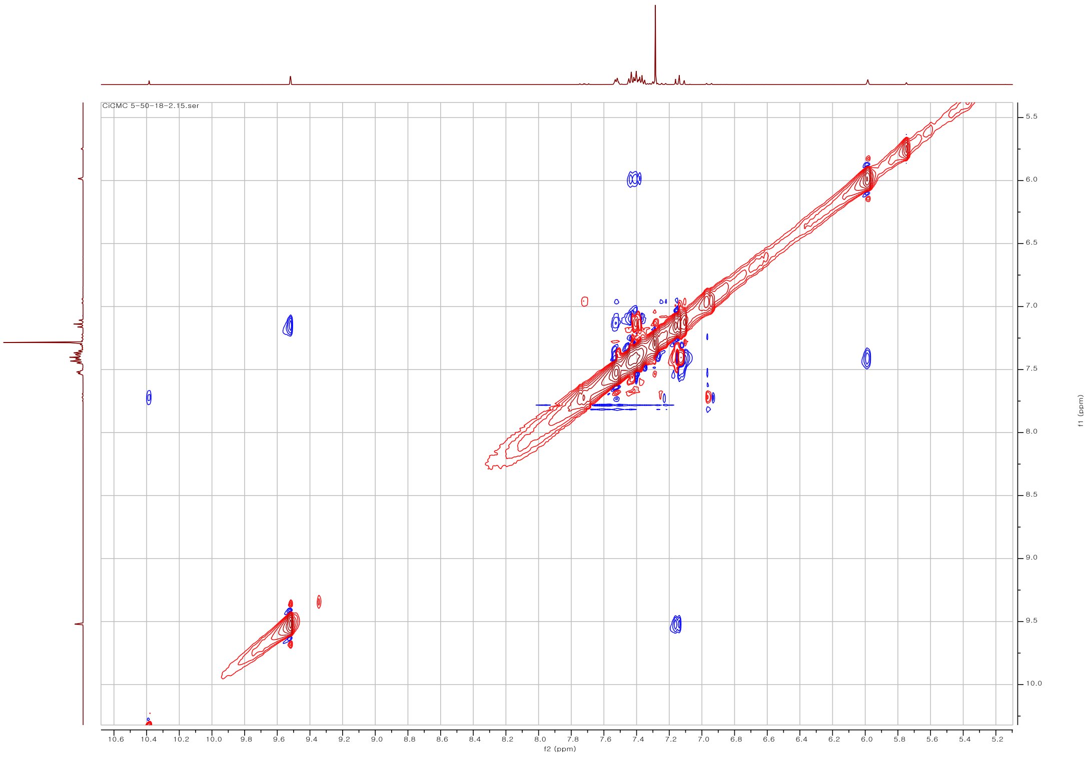

Fig. S.6.7. NOESY spectrum of **24** in $CHCl_3$-*d*

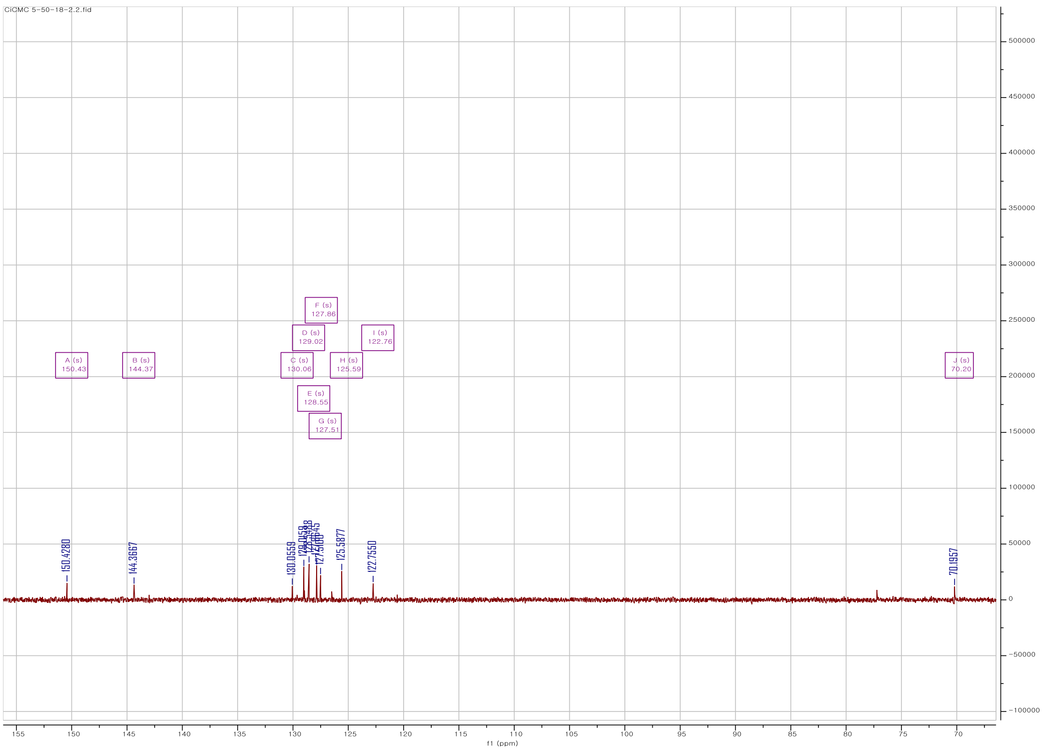

Fig. S.6.8. DEPT135 spectrum of **24** in $CHCl_3$-*d*
